# Strong phylogenetic signal despite high phylogenomic complexity in an Andean plant radiation (*Freziera,* Pentaphylacaceae)

**DOI:** 10.1101/2021.07.01.450750

**Authors:** Laura Frost, Ana M. Bedoya, Laura Lagomarsino

**Author notes:** These co-first authors contributed equally (Laura Frost, Ana M. Bedoya). Corresponding author: Laura Frost Laura Lagomarsino.

## Abstract

The Andes mountains of western South America are a globally important biodiversity hotspot, yet there is a paucity of resolved phylogenies for plant clades from this region. Filling an important gap to our understanding of the World’s richest flora, we present the first phylogeny of *Freziera* (Pentaphylacaceae), an Andean-centered, cloud forest radiation. Our dataset was obtained via hybrid-enriched target sequence capture of Angiosperms353 universal loci for 50 of the ca. 75 spp., obtained almost entirely from herbarium specimens. We identify high phylogenomic complexity in *Freziera*, including the presence of data artifacts. Via by-eye observation of gene trees, detailed examination of warnings from recently improved assembly pipelines, and gene tree filtering, we identified that artifactual orthologs (i.e., the presence of only one copy of a multi-copy gene due to differential assembly) were an important source of gene tree heterogeneity that had a negative impact on phylogenetic inference and support. These artifactual orthologs may be common in plant phylogenomic datasets, where multiple instances of genome duplication are common. After accounting for artifactual orthologs as source of gene tree error, we identified a significant, but non-specific signal of introgression using Patterson’s D and f4 statistics. Despite phylogenomic complexity, we were able to resolve *Freziera* into nine well-supported subclades whose evolution has been shaped by multiple evolutionary processes, including incomplete lineage sorting, historical gene flow, and gene duplication. Our results highlight the complexities of plant phylogenomics, which are heightened in Andean radiations, and show the impact of filtering data processing artifacts and standard filtering approaches on phylogenetic inference.

---

The Andean mountains in South America are one of the most species-rich areas of the world and serve as a center of diversity for many plant groups (Gentry 1982; Mutke and Barthlott 2005). The recent uplift of the Andes has resulted in some of the fastest plant evolutionary radiations reported to date (Madriñán et al. 2013; Hughes 2016), with some greatly phenotypically diverse clades (Hughes and Eastwood 2006; Lagomarsino et al. 2016). A significant portion of Andean plant biodiversity is yet to be described (Ulloa Ulloa et al. 2017) and many, if not most, Andean plant species, have never been included in a phylogeny. However, these phylogenies are fundamental toward understanding the evolutionary patterns that contribute to the origin and diversification of biodiversity in the World’s richest flora.

Establishing well-supported phylogenies for Andean plant groups is challenging for many reasons. Short divergence times between speciation events results in incomplete lineage sorting (ILS), incipient speciation, and introgression all contribute to poor phylogenetic resolution and high gene tree-species tree discordance (Vargas et al. 2017; Morales-Briones et al. 2018; Lagomarsino et al. 2022). This is further complicated by repeated whole genome duplication events throughout the evolutionary history of plants, at both deep and shallow scales (One Thousand Plant Transcriptomes Initiative 2019). There are additional practical limitations for phylogenetic inference in Andean systems. It is difficult to achieve full taxon sampling as species are often narrowly endemic and distributed in remote locations, and members of clades occur in many countries, each with different policies concerning collection and exportation of samples. As a result, achieving dense sampling of Andean-centered lineages commonly requires the use of herbarium specimens as a source of genetic material, which is associated with lower quality and quantity DNA than in freshly-collected or silica-dried leaf tissue (Bakker et al. 2015).

Improvements in methodology in the past decade bring us closer to achieving resolved phylogenies in previously intractable groups. Advancements in genomic sequencing, including hybrid-enriched target sequence capture, allow for the collection of hundreds to thousands of loci, even from degraded DNA from natural history collections (Bakker et al. 2015; McKain et al. 2018). Further, the development of universal probe sets facilitates sequencing of hundreds of loci for any system, regardless of genomic resources available (Johnson et al. 2019). Analytical methods are also increasingly able to accommodate multiple biological sources of gene tree discordance (Ogilvie et al. 2017; Solís-Lemus et al. 2017; Zhang et al. 2018).

Still, a major barrier to phylogenomic inference in many plant clades is the difficulty in efficient identification of orthologous and paralogous sequences from multi-copy loci (Yang and Smith 2014; Morales-Briones et al. 2022). The presence of multiple sequences for a single locus within an individual may result from allelic variation or gene/genome duplication. The latter process has repeatedly taken place during the evolutionary history of plant lineages, and results in the presence of non-orthologous gene copies within the same genome (i.e., paralogs) (Li and Barker 2020). Automated detection of multi-copy genes in phylogenomic datasets could be hindered when sequence data fails to meet contig number, contig length, or read depth thresholds set during assembly, a scenario that is more likely when is DNA obtained from herbarium specimens (Bakker et al. 2015).

The presence of only one copy of a multi-copy gene in a sequenced dataset (i.e., hidden paralogy) may be due to biological processes or to data processing artifacts. “Pseudo-orthologs” are a type of hidden paralogy that results from a biological process: differential loss after gene duplication leads to the presence of a single but non-orthologous copy across species in nature (Smith and Hahn 2021a, 2021b). While there is some evidence that coalescent-based phylogenetic methods are relatively robust to pseudo-orthologs (Smith and Hahn 2021a, 2021b), this finding is applicable only under certain conditions that do not include whole genome duplication followed by rediploidization— a phenomenon common in plants (Li et al. 2021). Hidden paralogy can also be artifactual, including when differential assembly of copies in a multi-copy gene results in the recovery of non-orthologous sequences for a given locus across taxa. We refer to these as “artifactual orthologs.” While it is well-documented that unrecognized paralogy can have negative impacts on species tree inference (Brown and Thomson 2017), the presence of artifactual orthologs in plant phylogenomic datasets and their impact on phylogenetic inference remains underexplored.

We combat the many challenges of Andean plant phylogenomics to infer the first phylogeny and introgression history of *Freziera* (Pentaphylacaceae), a group that constitutes a cloud forest plant radiation that previously lacked any phylogenetic information. *Freziera* includes 75 spp. of trees and shrubs that are widely distributed throughout montane regions of the Neotropics, from southern Mexico to Bolivia, with a center of diversity (61 spp.) in Andean cloud forests (Santamaría-Aguilar and Monro 2019; Fig. 1).There has been at least one whole genome duplication event in an ancestor of Pentaphylacaceae with the inclusive order Ericales (Larson et al. 2020), but ploidy levels in *Freziera* are unknown due to scarcity of genomic resources and chromosome counts. Chromosome counts in Pentaphylacaceae are highly variable (i.e., *Adinandra* [2 spp., n=42], *Cleyera* [1 spp., n=45], *Eurya* [4 spp., n=21, 29, 42], and *Ternstroemia* [2 spp., n=20, 25]; from Chromosome Counts Database [CCDB;(Rice et al. 2015); ccdb.tau.ac.il]).

**Figure 1.**
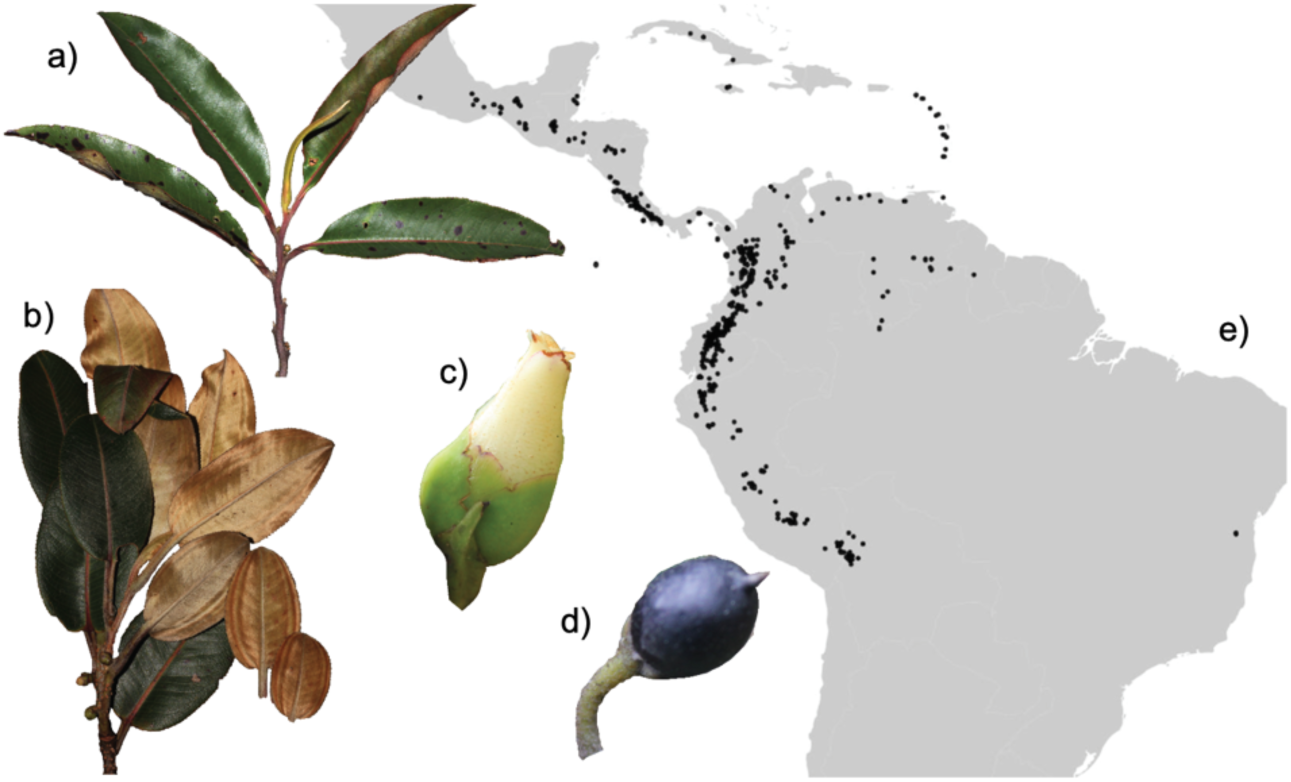
Diversity and distribution of *Freziera*. *Freziera* has significant variation in leaf morphology and pubescence, as illustrated by branches of a) *F. candicans* and b) *Freziera sp.*; meanwhile, c) flower and d) fruit morphology are relatively stable. *Freziera* is an Andean radiation; e) most species are distributed in this mountain chain in western South America, with some species in Central America, the Guiana Shield, the Caribbean, and the Atlantic Forest of Brazil. Photos by L. Lagomarsino.

Using Angiosperms353 (Johnson et al. 2019) target enrichment data derived almost entirely from herbarium specimens, we establish a phylogenetic baseline of *Freziera* despite widespread artifactual orthology and find evidence for historical introgression. We further explore how various types of data curation help remove artifactual orthologs and their impact on species tree inference. We finally provide suggestions to explore complex empirical phylogenomic datasets, especially those with a history of genome duplication and that are obtained with a high reliance on natural history collections.

## Materials and Methods

### Taxon Sampling

Ninety-four accessions representing 55 *Freziera* species—approximately 73% of the species diversity —were sampled for the ingroup. All but five came from herbarium specimens, which had an average age of 31.6 years (range: 7.4– 82.8; Supplementary Table 1). Nine accessions from other Pentaphylacaceae were sampled for the outgroup, including *Eurya japonica* (in the genus sister to *Freziera)*, *Cleyera albopunctata* (a member of tribe Frezierieae), and 7 species of *Ternstroemia* (belonging to the sister tribe Ternstroemieae; (Weitzman et al. 2004; Tsou et al. 2016).

### DNA extraction, library prep, target enrichment, and sequencing

Detailed descriptions of laboratory methods are provided in online Appendix 1. Briefly, DNA extraction followed a modified sorbitol extraction protocol (Štorchová et al. 2000). Library preparation used the KAPA Hyper Prep and KAPA HiFi HS Library Amplification kits with iTru i5 and i7 dual-indexing primers. Target enrichment was carried out using the MyBaits Angiosperms353 universal probe set (Johnson et al. 2019; Hale et al. 2020). DNA libraries were sequenced by Novogene in one Illumina Hiseq 3000 lane with 150 bp paired-end reads. Although most samples come from herbarium specimens, DNA concentration met minimum standards for sequencing.

### Raw data processing

Demultiplexed raw sequence reads were trimmed with illumiprocessor v2.0.9 (Faircloth 2013, 2016), a wrapper for Trimmomatic v0.39 (Bolger et al. 2014). Default settings were used and reads with a minimum length of 40 bp kept. Trimmed reads were assembled into supercontigs (exons and flanking intronic regions) with HybPiper v1.3.1 (Johnson et al. 2016) using the Angiosperm353 target file as reference (Johnson et al. 2019). In addition, we used a taxon-specific target file for Pentaphylacaceae using Easy353 (Zhang et al. 2022), a reference-guided assembly tool for recovery of Angiosperms353 gene sets. We used the transcriptome of *Ternstroemia gymnanthera* (One Thousand Plant Transcriptomes Initiative 2019) for sequence retrieval with Easy353. Additional details of data processing prior to final gene tree inference are available in online Appendix 1.

### Gene tree inference

Preliminary gene trees from the HybPiper v1.3.1 supercontig alignments were generated from aligned sequences for the 322 loci lacking paralog flags with RAxML v.8.2.12 (Stamatakis 2014) under the GTRCAT nucleotide substitution model with 200 rapid bootstraps. The resulting trees were processed with TreeShrink v1.3.3 (Mai and Mirarab 2018) to detect and remove unusually long branches (i.e., potential cross contaminants). Processing in TreeShrink was performed on a “per-gene” and “all-gene” basis. The identified branches (most of which corresponded to samples with few sequenced reads; Supplementary Table S1) were removed from final alignments. We then removed alignments with fewer than 25 ingroup samples from further analyses. Accessions present in <10% of processed gene trees were trimmed using the R package ape (Paradis and Schliep 2019a). Gene trees were inferred with IQ-TREE multicore v2.1.1 (Nguyen et al. 2015) including model section via ModelFinder (Kalyaanamoorthy et al. 2017), tree search, ultrafast bootstraps <u>(Hoang et al. 2018)</u>, and Shimodaira-Hasegawa-like approximate likelihood-ratio test (SH-aLRT; (Guindon et al. 2010; Anisimova et al. 2011).

### Paralog warnings and detection of artifactual orthologs

Despite assembling a single sequence per sample for the majority of loci and raising paralog warnings for <9% loci with HybPiper v1.3.1 (Supplementary Table S1), visual observation of gene trees and alignments suggested the presence of undetected paralog sequences (i.e., artifactual orthologs) in the dataset. Evidence for this included split clades of outgroup species and distinctive motifs in alignments that reflected higher sequence divergence than expected by allelic variation alone (Fig. 2b). Gene trees without paralog warnings were first examined by-eye in FigTree v1.4.3 (Rambaut 2014) and sorted as putative orthologs or artifactual orthologs. Artifactual orthologs are identified when different accessions for the same ingroup species, each represented by only a single sequence in the alignment, cluster into two subclades separated by relatively long internal branches (Fig. 2b). Visual detection of artifactual paralogs from gene trees in phylogenomic datasets thus benefits from sampling multiple individuals per species. We grouped loci with no paralog warnings from HybPiper v1.3.1. and confirmed by-eye as putative orthologs into a gene tree set called *orthologs*.*by.eye* (Table 1).

**Figure 2.**
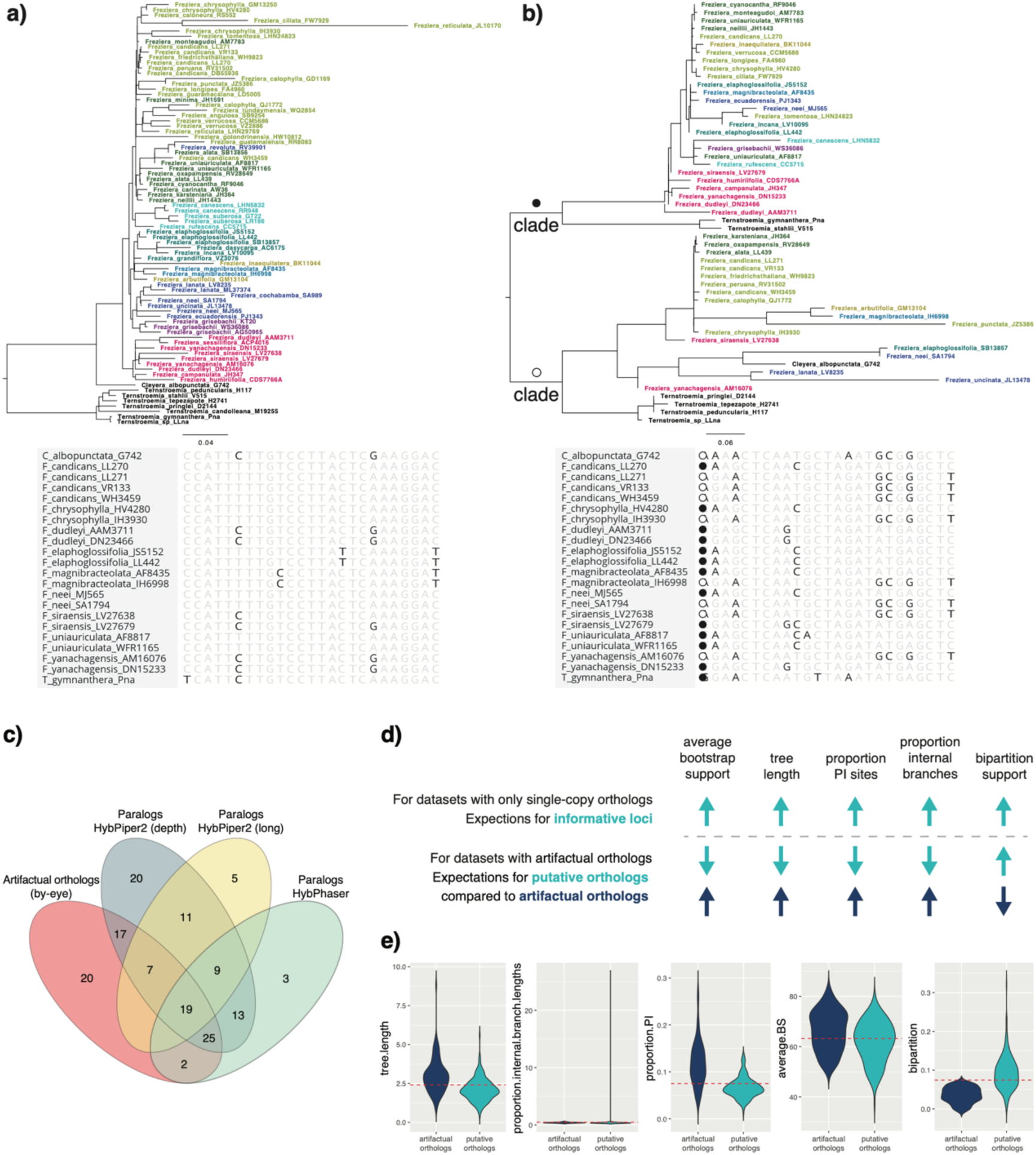
Exemplary gene trees and alignment subsets for a) a putative ortholog, and b) an artifactual ortholog. Putative ortholog gene trees exhibit monophyletic genera for outgroups and appropriate relationships to the ingroup, a relatively shallow backbone in the ingroup, and relationships similar to the preliminary species tree. A relatively low level of variation is present in alignments for putative orthologs, which is fitting of a locus in a universal probe set being applied within a genus. Artifactual ortholog gene trees often have deep divergences between subsets of the ingroup, including between samples from the same species. Within those subclades, patterns of relationships from the preliminary species tree are repeated. In some cases, as in b), outgroups are also non-monophyletic and subsets of outgroup samples are recovered as sister to the ingroup subclades. Alignments for artifactual orthologs exhibit relatively high variation with observable motifs between the suspected copies, as indicated by the black and white dots. c) Overlap between artifactual orthologs identified by-eye, and paralog warnings raised by automated paralogy detection strategies. d) Expectations for informative, single-copy orthologs change for many gene tree filtering metrics when data sets include artifactual orthologs; arrows indicate whether expected values for metrics are high or low. e) Impact of gene tree filtering strategies on removal of artifactual and putative orthologs as identified by-eye. The dashed lines indicate the various thresholds above which gene trees were kept in filtered datasets, as indicated in Table 1.

**Table 1.** Descriptions of the filtering criteria used, names applied to each of the datasets, number of loci removed and selected by each, and the impact of filtering criteria on gene tree discordance, branch support, and species tree topology. Values for datasets that performed equally or better than the orthologs.unfiltered dataset for each metric are bolded. Major topological conflicts between species trees and a consensus topology of the 11 filtered datasets (Fig. 3b) are shown, including instances where the species tree is compatible (+) or in disagreement (-) with the consensus topology. Datasets from which the Elaphoglossifolia group was not resolved as monophyletic are indicated with “n/a”. Asterisks mark disagreement from the alternative placement of only one species. The “Branching order of the Elaphoglossifolia group” includes conflict in the placement of F. magnibracteolata.

| Description of filtering criteria (for keeping loci) | Putative orthologs dataset name after filtering | No. of loci removed by filtering criteria | No. of loci that passed filtering criteria | Normalized Quartet Score | Average branch support | Proportion of highly supported branches (PP > 0.97) | Two clades in Candicans group | Monophyletic Elaphoglossifolia group | Branching order of Elaphoglossifolia group | Placement of Arbutifolia clade |
| --- | --- | --- | --- | --- | --- | --- | --- | --- | --- | --- |
| loci not flagged as paralogs by HybPiper v.1.3.1 | *orthologs.unfiltered* | 31 | 313 | 0.567 | 0.774 | 0.390 | + | – | n/a | + |
| loci identified as putative orthologs after by-eye inspection of gene trees and alignments | *orthologs.by.eye* | 131 (90) | 182 | **0.634** | **0.792** | 0.377 | + | + | + | + |
| loci with 1 contig at ≥75% reference sequence (no long paralog warnings from HybPiper v.2.0) | *orthologs.HybPiper2.long* | 51 | 262 | **0.582** | **0.780** | 0.377 | + | + | + | + |
| loci without paralog warnings by depth or length from HybPiper v.2.0 | *orthologs.HybPiper2.no.warnings* | 126 | 187 | **0.619** | **0.783** | 0.351 | + | + | + | + |
| loci that have a lower proportion of SNPs than 1.5x the interquartile range above the 3rd quartile for any given sample | *HybPhaser* | 71 | 242 | **0.597** | **0.783** | 0.364 | + | + | + | + |
| orthology inference via gene tree pruning using homologs with monophyletic, non-repeating outgroups | *MO* | 91 | 222 | **0.662** | 0.664 | 0.221 | + | + | – | – |
| alignment contains ≥ 7.5% parsimony informative sites in ingroup (average value across all alignments) | *proportion.PI* | 153 | 160 | 0.528 | 0.715 | 0.247 | – | – | n/a | + |
| gene trees with internal branch lengths comprising an above average proportion of the total tree length across all gene trees | *proportion.internal.branch.lengths* | 164 | 148 | 0.542 | 0.685 | 0.234 | – | + | + | + |
| gene trees with average bootstrap support across all nodes above the average value across all gene trees | *average.BS* | 150 | 163 | 0.560 | 0.727 | 0.234 | + | + | – | + |
| gene tree with total tree length above the average value across all gene trees | *tree.length* | 173 | 139 | 0.524 | 0.688 | 0.260 | – | – | n/a | – |
| gene trees with above average bipartition support relative to the species tree inferred from the *orthologs.unfiltered* dataset | *bipartition* | 147 | 166 | **0.601** | **0.792** | **0.390** | + | + | + | + |

We compared our by-eye identification of artifactual orthologs with two automated paralogy detection strategies: HybPiper v2.0 (Johnson et al. 2016) and HybPhaser (Nauheimer et al. 2021). Both versions of HybPiper could detect paralogs if multiple assembled contigs each cover ≥75% of a reference sequence; this returns a long paralog warning in HybPiper2. HybPiper2 also raises paralog warnings when multiple contigs are assembled across ≥75% of the reference sequence even when individual contigs are <75% the full sequence length; this returns a paralog-by-depth warning. In addition, HybPiper2 reports stitched contigs (i.e., those derived from multiple SPAdes contigs), which can be used to identify chimeric sequences (i.e., stitched contigs in which the two separate sequences are derived from different gene copies); HybPiper2 returns chimera warnings when a sufficient number of read pairs from stitched contigs map to different SPAdes contigs and have sequence mismatch above a given threshold for one read but not the other (see HybPiper2 documentation for additional details; https://github.com/mossmatters/HybPiper/). HybPhaser is an automated strategy to identify hybrids, polyploids, contamination, and paralog sequences by identifying samples with a high proportion of heterozygous loci and allele divergence, and loci with a high proportion of SNPs (Nauheimer et al. 2021).

In addition, we applied a method for automated orthology detection (i.e., gene tree pruning) using homologs with monophyletic outgroup (MO; Table 1; Yang and Smith 2014; Morales-Briones et al. 2021a). MO identifies clusters in gene trees with monophyletic outgroups and searches from root to tips for duplications. When duplicated taxa are found on either side of a bifurcation, the subtree with the fewest ingroup taxa is pruned. MO does not directly identify artifactual orthologs but has the potential to remove their effect by selecting orthogroups and allowing the inclusion of more loci in a final dataset.

We generated five sets of gene trees to compare artifactual orthology identification using the above automated pipelines; a description of filtering criteria and thresholds for each dataset are in Table 1. Loci that received a paralog warning in HybPiper v1.3.1 were excluded to generate the *orthologs.unfiltered* dataset. The remaining filtered datasets (i.e., *orthologs.HybPiper2.long*, *orthologs.HybPiper2.no.warnings, HybPhaser,* and *MO*) are subsets of the *orthologs.unfiltered* dataset. For the *HybPhaser* dataset, we did not filter loci by missing data (we kept loci recovered in at least 0.01 proportion of taxa) to ensure that only loci with excess heterozygosity (loci with more than 1.5× the interquartile range above the third quartile), as expected in putative paralogs, were removed. We kept ortholog groups with ≥25 ingroup taxa in the *MO* dataset for comparison with the other filtering strategies.

### Impact of standard gene tree filtering strategies on the presence of artifactual orthologs

We explored how standard gene tree filtering affects the presence of artifactual orthologs (as identified by-eye) in our dataset. Gene tree filtering is typically applied to minimize gene tree estimation error, gene tree discordance (Molloy and Warnow 2018), and for phylogenomic subsampling (Mongiardino Koch 2021) but applying these metrics may also favor the selection of artifactual orthologs in a dataset (Fig. 2d,e). Gene trees were filtered using five common empirical criteria for phylogenomic subsampling (Table 1): one alignment-based metric (*proportion.PI*), two tree length metrics (*tree.length* and *proportion.internal.branch.lengths*), and two gene tree support metrics (*average.BS* and *bipartition*). Due to the presence of non-orthologous copies in alignments including artifactual orthologs, a high degree of genetic variation is expected relative to true orthologs, increasing the proportion of informative sites and, because of well-supported clusters of non-orthologous sequences, we expected an elevated average bootstrap support in gene trees (Fig. 2d). Filtering by bootstrap support is standard practice, collapsing low supported nodes in gene trees prior to species tree inference to remove gene tree error (Zhang et al. 2018). The two tree length criteria were selected because, while a high percentage of internal branch length can signal high phylogenetic signal (Shen et al. 2016), it can also indicate biological pseudo-orthologs (Smith and Hahn, 2016b) and likely, artifactual orthologs (Fig. 2). Finally, filtering by bipartition support may remove artifactual orthologs by favoring loci that are more concordant with the inferred species tree (Smith et al. 2018; Fig. 2e).

Standard gene tree filtering strategies were applied to the *orthologs.unfiltered* dataset. The *proportion.PI* was obtained using AMAS (Borowiec 2016) with outgroups removed. Estimates of *proportion.internal.branch.lengths* were extracted from IQ-TREE ‘*.iqtree’ output files. Bootstrap support values were extracted from IQ-TREE maximum likelihood tree files and averaged to calculate *average.BS*. For *tree.length*, input gene trees were rooted using pxrr in phyx (Brown et al. 2017) and tree length was calculated with the script ‘get_var_length.py’ of SortaDate (Smith et al. 2018) excluding outgroups. *Bipartition* support was calculated with SortaDate (Smith et al. 2018)‘get_bp_genetrees.py’ script(Smith et al. 2018) against a species tree generated with ASTRAL-III (Zhang et al. 2018) from the *orthologs.unfiltered* dataset. Cutoff thresholds for each filter were selected near the average value in our dataset to keep a similar number of loci for each criterion.

### Phylogenetic relationships and introgression in Freziera

#### Impact of filtering criteria on species tree inference

Species trees were inferred in ASTRAL-III from the 11 datasets listed in Table 1. A majority-rule consensus tree was generated from the species trees for the 11 filtered datasets (*consensus* function of the R package *ape*; Paradis and Schliep 2019b) to identify major clades that were concordant across datasets. Gene tree discordance was assessed for all species tree topologies using the final normalized quartet score (NQS; proportion of quartet trees in gene trees that are present in the species tree (Mirarab et al. 2014b) estimated with ASTRAL-III. Impact of filtering criteria on branch support was assessed by calculating the average branch support and proportion of well-supported branches (ppl≥0.97; Rabiee and Mirarab 2020) across the 11 resulting species trees. We inferred a second consensus tree from the species trees generated from the datasets that showed the greatest improvements to both normalized quartet score and average branch support (*bipartition, by.eye, HybPhaser,* and *orthologs.HybPiper2.no.warnings*), which also had substantial reductions in the number of artifactual orthologs (Fig. 2c). The resulting tree was compared to the *orthologs.unfiltered* dataset to explore the impact that the removal of paralogs, including artifactual orthologs, has on species tree inference.

#### Signals of introgression in Freziera

To assess genomic evidence of introgression in species of *Freziera*, we calculated Patterson’s D and f4 statistics using the *Dtrios* function in Dsuite v.0.4 r43 (Malinsky et al. 2021). The input VCF file was generated with dDocent (Puritz et al. 2014) by mapping all *Freziera* trimmed reads to the sequences of *T. tepezapote* for the loci in the *bipartition*, *by.eye*, and *hybiper2.no.warnings* datasets. We called SNPs using default values for all mapping parameters. The resulting VCF files were filtered with vcftools (Danecek et al. 2011). We retained SNPs with <50% missing data and retained only one SNP per 100 bp window to decrease the likelihood of including linked SNPs. The statistical significance of the D and f4 was assessed using block jackknife on windows of 75–78 SNPs followed by Benjamini-Hochberg correction as implemented in R (Benjamini and Hochberg 1995) to assess family-wise error rate (following (Malinsky et al. 2018). The D and f4 statistics were estimated for all possible trios across three datasets (13,244, 15,180, 14,190 trios for the *bipartition*, *by.eye*, and *hybiper2.no.warnings* respectively). The ASTRAL-III species trees inferred from each of the three datasets were specified in Dsuite so that the D and f4 estimated values were arranged according to the tree.

The relatively small number of loci in our dataset limited the power to detect introgression along a phylogeny. The f-branch statistic, which accounts for the correlation of D and f4 due to shared ancestry among multiple potential introgression donor species (Malinsky et al. 2018), allows for a better interpretation of introgression patterns across a tree (Malinsky et al. 2021). However, simulation analyses have shown that for a relatively small number of unlinked SNPs (<10,000), the proportion of cases where the strongest inferred f-branch signal corresponds to the correct simulated gene flow recipient and donor branches is <20% (Malinsky et al. 2021). Due to the limited number of unlinked SNPs in our targeted sequence capture dataset, we did not apply this metric.

## Results

### Sequence capture efficiency

Voucher information for accessions and per sample data for read trimming and contig assembly are available in Supplementary Table S1. Of the 102 accessions for which target enrichment libraries were prepared and sequenced, 22 (21 *Freziera* and 1 outgroup) were excluded either because no sequences were assembled for any of the 353 target loci, or because they were identified to have exceedingly long branches with TreeShrink (most of the latter corresponded to samples with few sequenced reads; Supplementary Table S1). Target enrichment efficiency across all sequenced samples is shown in Supplementary Figure S1. Seventy-two accessions representing 50 of the 75 species of *Freziera* and 8 members of the outgroup remained (see methods; Supplementary Table S1), which had assembled sequences for 52–348 genes (average: ∼296). Standard assembly statistics from both versions of HybPiper are in Supplementary Tables S1-S2. No loci were flagged by HybPiper2 as putatively chimeric. However, all but 13 loci included at least one sequence with stitched contigs (Supplementary Table S3). All sequences without stitched contigs correspond to regions spanning a single exon. Off-target data collection was insufficient to assemble plastomes.

### Paralog warnings and detection of artifactual orthologs

The 11 paralogy detection and gene tree filtering criteria that we applied generated datasets with 140–313 loci (Table 1). Results of artifactual ortholog detection by-eye and from automated strategies are shown in Fig. 2a and Supplementary Table S3. HybPiper v.1.3.1 raised paralog warnings for 31 loci (i.e., >1 contig was assembled for at least one sample for a given locus). Of the 322 loci that were not flagged as paralogs, nine were removed because they contained fewer than 25 ingroup samples, leaving 313 with a single assembled contig per species. By-eye inspection of these 313 loci resulted in the identification of 90 artifactual orthologs. Loci flagged as potentially paralogous in HybPiper v.1.3.1 also had long paralog warnings and paralog warnings by depth with HybPiper2. Of the 313 loci without paralog warnings from HybPiper v.1.31, 51 had long paralog warnings, of which 46 also received paralog warnings by contig depth with HybPiper2. An additional 75 loci were not flagged as long paralogs but did receive paralog warning by contig depth for at least one sample, totaling 121 loci with warnings by contig depth and 126 loci with any kind of paralog warning issued by HybPiper2. HybPhaser identified 73 loci as putative paralogs (Supplementary Table S3; online Appendix 2), most of which also had paralog warnings by depth. There was substantial overlap of loci identified by these methods (Fig. 2c). Three of the methods we compared— HybPiper2 paralog-by-depth warnings, HybPiper2 paralog-by-length warnings, and HybPhaser— flagged 70 (78%) of the loci that we identified as a artifactual orthologs in our by-eye assessment (Fig. 2c). Of these, HybPiper2 paralog-by-depth warnings had the greatest overlap with our by-eye assessment (Fig. 2c; Supplementary Table S3). Assembly with a taxon-specific target file resulted in similarly low levels of paralog detection as the universal target file: <8% of assembled loci recovered >1 long contig for at least one sample (Supplementary Table S4). There was a correlation between the number of reads mapped and the number of paralog warnings from HybPiper in a sample (Supplementary Fig. S3).

### Impact of standard gene tree filtering strategies on the presence of artifactual orthologs

Summary statistics for alignments and gene trees are available in Supplementary Table S3. The impact of gene tree filtering strategies on removal of artifactual and putative orthologs (as identified by-eye) is shown in Fig. 2e. Putative orthologs are removed with all filtering strategies, and only filtering gene trees by bipartition support resulted in the removal of most artifactual orthologs. As commonly applied, the remaining four gene tree filtering criteria all resulted in datasets in which there were either similar proportions of putative orthologs and artifactual orthologs in the final datasets (*average.BS, proportion.internal.branch.length*), or that favored the inclusion of artifactual orthologs (*tree.length*, *proportion.PI*).

### Impact of data curation on phylogenetic performance

We found that removing artifactual orthologs from our datasets improved phylogenetic performance metrics, and that, with the exception of filtering by bipartition support, gene tree filtering did not. Datasets in which curation reduced the proportion of artifactual orthologs (*orthologs.by.eye, orthologs.HybPiper2, HybPhaser*, and *bipartition*) had the highest normalized quartet scores and average branch support (Table 1). Except for *bipartition*, these values were consistently lower in datasets resulting from standard gene tree filtering methods (Table 1), likely due to an increased proportion of artifactual orthologs relative to the unfiltered dataset (Fig. 2b). Relative to the unfiltered dataset, the number of highly supported branches was similar in datasets in which artifactual orthologs were filtered out, and lower in datasets that underwent conventional gene tree filtering, again apart from *bipartition* (Table 1). While there was some impact on removing artifactual orthologs on branch support, it is possible that these results were relatively small due to the high degree of gene tree heterogeneity in our dataset (Table 1).

We also found consistent impacts of removing artifactual orthologs on the species tree topology. Datasets where artifactual orthologs were removed (*by.eye, HybPiper2.long, HybPiper2.depth, HybPhaser, bipartition*) consistently recovered the Elaphoglossifolia group as monophyletic. Contrastingly, species tree topologies generated by gene tree filtering differed from any of those generated from datasets that were curated to remove artifactual orthologs, and sometimes introduced relationships that were found in no other analyses, including of the unfiltered dataset (e.g., non-monophyletic Callophylla and Karsteniana clades and placement of the Arbutifolia clade; Table 1; Fig. 3b).

**Figure 3.**
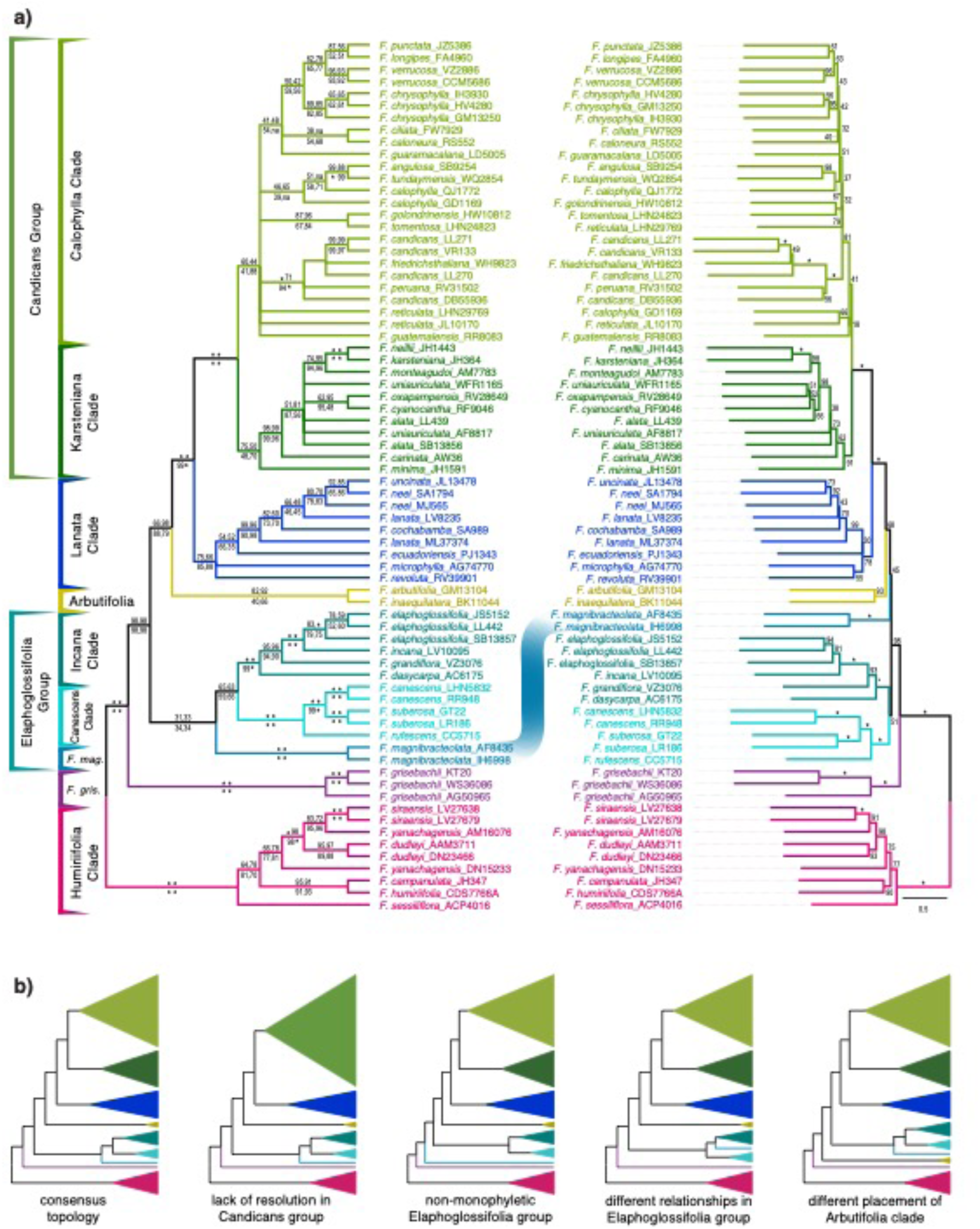
Names of clades consistently recovered and species tree topologies for a) The consensus tree of the four datasets in which the proportion of artifactual orthologs was reduced (*bipartition*, *by.eye*, *HybPhaser*, *orthologs.HybPiper.no.warnings*; left) versus the orthologs.unfiltered species tree (right). The placement of *F. magnibracteolata–* the major topological conflict between the two trees– is highlighted with the blue line connecting the tip labels in each. Branches with full support (LPP from ASTRAL = 1) are indicated by an asterisk; elsewhere, the LPP * 100 is provided. Support for each of the four datasets in the consensus tree are given clockwise from the top left: *bipartition*, *by.eye*, *HybPhaser*, *orthologs.HybPiper.no.warnings*. b) Cartoon trees displaying the most frequent differences in relationships between clades recovered by different datasets and the consensus tree of all datasets. Cartoons depict one topological scenario but not the only topological outcome.

### Phylogenetic relationships and introgression in Freziera

The majority-rule consensus tree of all 11 species trees resulting from data filtering and the consensus tree of the four datasets with the lowest proportions of artifactual orthologs were congruent in the major clades recovered and the relationships between those clades. Seven clades that were frequently inferred across analyses were identified—the Humiriifolia, Canescens, Incana, Arbutifolia, Lanata, Karsteniana, and Calophylla clades—along with *F. grisebachii* and *F. magnibracteolata*, which formed monotypic clades (Fig. 3a). The Elaphoglossifolia group, comprising the Canescens and Incana clades and F. magnibracteolata, was additionally recovered as monophyletic in both consensus trees. Five of the 11 datasets produced species trees consistent with monophyly of the nine clades and their branching order in the consensus tree (Table 1; Supplementary Fig. S2). Among species trees consistent with the consensus topology, some branches at deep nodes were inferred with high support (ppl≥0.97) across all analyses: the common ancestor of all *Freziera*, the common ancestor of core *Freziera* (all *Freziera* excluding the Humiriifolia clade), and the successive node within core *Freziera*, the Humiriifolia, Canescens, and Incana clades, and the Candicans group, which comprises the Karsteniana, and Calophylla clades (Fig. 3). Species-level relationships within most subclades were consistent across analyses except for the species-rich Candicans group (Supplementary Fig. S4).

Regions of lowest support in the consensus tree tended to be in conflict between species trees and the consensus topology. These regions often involved two notable sets of taxa: the Elaphoglossifolia group, comprising the Incana and Canescens clades and *F. magnibracteolata*, and the Candicans group, which comprises the Calophylla and Karsteniana clades (Table 1, Fig. 3; Supplementary Fig. S4). While the Elaphoglossifolia group was monophyletic in most datasets, including all in which the proportion of artifactual orthologs was reduced (Table 1), the unstable placement of *F. magnibracteolata* rendered it non-monophyletic three datasets: *orthologs.unfiltered*, *proportion.PI*, and *tree.length* (Table 1; Fig. 3b; Supplementary Fig. S2a,g,j). Contrasting with the Elaphoglossifolia group, the monophyly of the Candicans group was identified in all analyses, generally with high support despite a high degree of gene tree heterogeneity; however, the resolution of its constituent Calophylla and Karsteniana subclades varied across analyses (Table 1). While all datasets in which the proportion of artifactual orthologs was reduced resolved the Calophylla and Karsteniana subclades as monophyletic, species-level relationships within these groups varied (Supplementary Fig. S4).

For the *bipartition*, *by.eye*, and *orthologs.HybPiper2.no.warnings* datasets, p-values for the estimated D statistics could not be calculated for 75.9%, 85.5%, and 93.6%, respectively, of all evaluated trios due to lack of ABBA-BABA variants (Supplementary Fig. S5). Estimates of Patterson’s D >0 were significant (corrected p-value ≤ 0.05, Z-score >3) for 104 (3.27%), 162 (7.38%), and 46 (5.08%) trios out of the remaining evaluated trios. The number of statistically significant D statistics was probably underestimated in our study given the large proportion of trios for which p-values could not be calculated due to a lack of ABBA-BABA variants (∼76%– 94%). Resulting f4-ratio statistics across the three datasets also show evidence for multiple instances of introgression in *Freziera* (Supplementary Fig. S5). These results were further supported by the results from HybPhaser, which identified several samples with >80% locus heterozygosity (LH) and >1% allele divergence (AD) across all major groups recovered (online Appendix 2; Supplementary Fig. S6). This is indicative of prevalence of introgression and polyploidy (Nauheimer et al. 2021) in *Freziera* species included in this study.

## Discussion

Relying on DNA extracted primarily from herbarium specimens, we inferred the first phylogeny of *Freziera* (Pentaphylacaceae), an understudied tropical plant lineage. Our museomic dataset, representing hybrid-enriched target sequence capture of Angiosperms353 loci, highlights the many challenges of working with understudied clades even in the genomic era: no a priori phylogenetic hypothesis, a universal bait set, poor-quality DNA, and high proportion of paralogs, many of which we identified as artifactual orthologs. Despite phylogenomic complexity including a high degree of gene tree heterogeneity, we resolve *Freziera* into nine clades whose histories have been shaped by myriad evolutionary processes, including incomplete lineage sorting, introgression, and gene or genome duplication. In the face of these complexities, we identify that the biggest improvements to phylogenetic inference in *Freziera* did not come from filtering gene trees to maximize phylogenetic informativeness. Instead, they came from reducing the noise from artifactual orthologs, which was accomplished in a variety of ways: observing data by-eye, implementing automated pipelines, and identifying gene tree filtering mechanisms that are consistent with reducing this artifact. We offer recommendations on strategies for removal of similar data processing artifacts for phylogenetic inference of groups where multi-copy loci are expected to be prevalent.

### Identification of artifactual orthologs using automated pipelines

We observed a widespread pattern of artifactual orthology in our dataset, in which multi-copy genes were recovered as single copy due to errors in the assembly process. These loci had multiple identifiable motifs in their alignments, and their resulting gene trees typically exhibited polyphyly of species and deep divergences between clades, including outgroup taxa and members of those spuriously polyphyletic species (Fig. 2b). As troubling as the prospect of unfiltered paralogs may be for empiricists, artifactual orthologs may be easier to detect than biological pseudo-orthologs (Smith and Hahn 2021b) because of these striking, easily identifiable patterns.

Using our visual inspection as a baseline, we were able to assess the ability of automated paralogy detection pipelines to identify artifactual orthologs. There was substantial overlap in the loci that we identified by-eye and those removed through various automated paralog detection mechanisms (Fig. 2a). While artifactual orthologs result from the assembly of one contig for a truly multi-copy locus, they were often identifiable if more than one short contig covered >75% of the target length (the strategy used in *orthologs.HybPiper2.no.warnings*). Similarly, but to a lesser extent, artifactual orthologs were removed by filtering out loci with high proportion of SNPs (*HybPhaser*) and with the assembly of multiple contigs at at least 75% of sequence length (*orthologs.HybPiper2.long*). We recommend by-eye identification as an initial strategy to explore whether artifactual orthologs are present in an individual empirical dataset, and emphasize the utility of sampling multiple individuals per species.

Our dataset relied heavily on degraded DNA extracted from herbarium tissue, which we believe contributes to the assembly of artifactual orthologs. Though high-molecular weight DNA is fragmented during library preparation, the desired fragment size for Illumina libraries is 200- 500 base pairs (bp; Bronner and Quail 2019). As is common when working with specimens from the wet tropics (Bakker et al. 2015), many of our samples had a high proportion of fragments <200 bp (unpublished). These short fragments restrict data collection in non-coding regions, as fragments are less likely to span the regions flanking targeted exons. Using HybPiper v2.0.1, we identified a high proportion of stitched contigs—separate contigs concatenated into a single sequence. The assembly of multiple contigs belonging to different paralog copies into a single sequence (i.e., chimeric sequences) is an artifact that is possible in the presence of stitched contigs. The preferential selection of the longest contig with the highest coverage for a given gene region during assembly further contributes to decreased paralog detection and increased chimeric assembly. Despite the high proportion of stitched contigs, no genes were flagged by HybPiper v2.0.1 with a chimera warning for any sample (Supplementary Table S2). This is likely due to current limitations and stringency in chimeric sequence detection. Specifically, paired ends must completely map to separate contigs and reads must map entirely within an exon on the separate contigs to flag a chimera warning. Short fragments from degraded DNA reduce the likelihood of finding read pairs that will meet these criteria, and, therefore, the likelihood of detecting chimeric sequences, though they may be present in the dataset.

Effectively identifying chimeric sequences from single or stitched contigs (as even a single assembled contig for a given locus may be a chimera of different alleles) remains a challenge in the assembly of target capture data from short read sequences, especially as assembly pipelines cannot distinguish between reads derived from different paralogs or alleles. Detection of chimeric sequences and its impact on phylogenetic inference remains a fundamental problem in phylogenomics, particularly in groups where polyploidization and hybridization are suspected to be prevalent (Morales-Briones et al. 2018). While not possible with degraded DNA from herbarium specimens, a potential solution includes generating long-read sequence data at a sufficient depth for accurate phasing of copies.

We believe the lack of data spanning introns, resulting in a high proportion of stitched contigs, and the potential for greater disparity between capture of both copies from herbarium DNA, is a primary source of artifactual orthologs. Low sequencing depth resulting in poor coverage of non-coding regions is another factor that can increase the proportion of stitched contigs and/or differential success capturing copies. However, sequencing depth is unlikely to be the main source of artifactual orthologs in our dataset as the number of reads mapped to target loci (Supplementary Table S1) in our dataset is higher per sample than studies that have successfully examined paralogs in Angiosperms353 datasets (Johnson et al. 2019; Gardner et al. 2020). Taken together, this highlights caveats that remain with museomic data: long-read sequence data is not an option in many cases and deeper sequencing cannot recover missing data. Herbarium specimens are an invaluable resource for improving sampling and filling gaps in the tree of life, and we do not advocate for the exclusion of herbarium specimens in phylogenomic datasets. Rather, we recommend careful assessment of datasets considering these artifactual complications in assembly.

### Impact of standard gene tree filtering strategies in the presence of artifactual orthologs

Gene tree filtering is a commonly applied strategy to minimize gene tree estimation error (GTEE) and its impact on species tree estimation (Molloy and Warnow 2018). Without an explicit method to estimate GTEE in empirical data, multiple criteria including alignment length, the number/proportion of variable or parsimony informative sites, total tree length, the proportion of internal branch lengths, and average bootstrap support of gene trees have all been used as implicit proxies to account for GTEE (Leaché et al. 2014; Liu et al. 2015; Shen et al. 2016; Blom et al. 2017). It is argued that these metrics select for “higher quality” gene trees, however, this assumption is violated in the presence of artifactual orthologs, which are associated with higher values of many standard metrics (Fig. 2c).

Only one filtering criterion successfully reduced the proportion of artifactual orthologs in the *Freziera* dataset: bipartition support. The high efficiency of bipartition support (Fig. 2e) is likely due to the major topological differences between the consensus species tree and gene trees of artifactual orthologs (Fig. 2). However, a significant proportion of putative orthologs were also removed through this filtering mechanism. This curation strategy should be used with caution and may require additional assessment of loci falling below the threshold if investigating biological processes such as ILS or introgression, as some highly discordant, single-copy loci will also be filtered. Given that the most successful methods of automated paralog detection require assembly with HybPiper, bipartition support could provide a useful metric by which users can remove artifactual orthologs in datasets assembled using other pipelines.

Both artifactual orthologs and putative orthologs were at removed similar levels from datasets by the remaining four gene tree filtering criteria, in particular *tree.length* and *proportion.PI.* This is consistent with our predictions about characteristics of artifactual orthologs (Fig. 2d,e) and demonstrates that filtering by alignment or gene tree characteristics using standard techniques may actually result in a relative increase of data artifacts in complex phylogenomic datasets.

While gene tree filtering metrics have the potential to significantly improve phylogenetic support (Doyle et al. 2015) and clarify relationships, it may be an inappropriate strategy for some studies (Molloy and Warnow 2018). This is true in this case for *Freziera*, in which gene tree filtering resulted in lower branch support and spurious topological inference for all gene tree filtering criteria except bipartition support (Table 1, Supplementary Fig. S2). This is likely the result of removal of a large proportion of single-copy loci relative to artifactual orthologs (Fig. 2b). The impact of the presence of artifactual orthologs and gene tree filtering strategies on species tree inference will likely vary across datasets. We recommend careful observation of phylogenomic datasets before applying these criteria of data curation.

### Impact of data curation on phylogenetic performance in the face of artifactual orthologs

Species tree inference of *Freziera* with a two-step, coalescent-aware species tree inference algorithm (i.e., ASTRAL-III) was relatively robust to artifactual orthology. This is consistent with recent studies demonstrating that species tree methods are robust to the presence of paralogs (Yan et al. 2022), though undetected paralogs have also been documented to mislead species tree inference (Brown and Thomson 2017; Siu-Ting et al. 2019). Across analyses of our eleven datasets, backbone relationships were largely consistent and topological differences were primarily concentrated in a few portions of the phylogeny, at least one of which (the Candicans group) corresponds to a rapid radiation where levels of ILS and introgression are likely high. We found that the unfiltered dataset (i.e., *orthologs.unfiltered*) had a slightly higher proportion of well-supported nodes, though this number was close in absolute value to those from curated datasets with a reduced proportion of artifactual orthologs (Table 1). This may be simply a result of a larger dataset, as coalescent-based phylogenetic inference algorithms require a large number of independent loci to resolve challenging relationships (Leaché and Rannala 2011; Mirarab et al. 2014a).

Reducing artifactual orthologs had an overall positive impact on phylogenetic inference and support (Table 1; Fig. 3). Curated datasets where artifactual orthologs were removed (*orthologs.by.eye, orthologs.HybPiper2, HybPhaser, bipartition*; Table 1) had higher normalized quartet scores and average bootstrap support relative to the unfiltered dataset, and consistently recovered the monophyly of the Elaphoglossifolia group via the placement of *F. magnibracteolata*, a relationship that was not present in the unfiltered dataset (Table 1). Improvements in these datasets are likely the result of the reduction in gene tree heterogeneity with the removal of the noise introduced by artifactual orthologs, though gene tree heterogeneity persists in curated datasets and likely reflects biological processes that have shaped *Freziera*’s evolutionary history (Table 1). Ensuring that paralogs, including artifactual orthologs, are appropriately handled in phylogenetic analyses is not only essential for accurate phylogenetic estimation, it is also crucial to downstream analyses that require accurate branch lengths, including divergence date estimation (Siu-Ting et al. 2019).

Orthology inference using monophyletic outgroup (*MO*) was not associated with improved phylogenetic performance, despite being successfully applied in datasets where paralogs are represented as multiple copy loci in assembled datasets (Morales-Briones et al. 2022). The *MO* species tree had reduced gene tree discordance (i.e., higher normalized quartet sampling score) relative to the unfiltered dataset, likely from the removal of non-orthologous copies. However, it failed to recover some of the major clades identified from curated data where artifactual orthologs were removed. This may be due to information loss: the high number of multicopy loci in our dataset resulted in many trees being extensively pruned by MO, resulting in smaller subtrees with fewer taxa. MO is a valuable tool to identify orthologous clusters in complex phylogenomic datasets with multi-copy paralogs and can increase phylogenetic resolution in groups with a history of polyploidy (Morales-Briones et al. 2022); however, it cannot be used to identify artifactual orthologs and did not improve phylogenetic performance in *Freziera*, where paralogs were predominantly single-copy.

### Phylogenetic relationships and introgression in Freziera

One of the barriers to the study of Neotropical diversification is the difficulty resolving phylogenies of recent, rapid Andean radiations. Despite *Freziera*’s low species richness compared to many cloud forest plant clades, we find similar hallmarks of explosive radiation in our phylogenetic results. In the face of phylogenomic complexity—including the presence of multiple copy paralogs, artifactual orthologs, rapid radiations, and DNA extracted from herbarium specimens—relationships among species and subclades of *Freziera* were consistently inferred across curated datasets. Most topological differences across species trees in datasets with a reduced proportion of artifactual orthologs were within the Candicans group, a rapid radiation with short internal branch lengths. We resolve *Freziera* into nine subclades, most of which are moderately to well supported (Fig. 3). Despite their monophyly, these clades generally lack morphological synapomorphies and are not geographically structured. The Humiriifolia clade, which is sister to core *Freziera*, best represents the wide morphological diversity in each subclade: its species have among the largest (*F. humiriifolia*) and smallest (*F. yanachagensis*) leaves in the genus, despite occurring in close proximity in the Cordillera del Cóndor region of southern Ecuador. While our phylogenetic backbone is generally well-supported, there are three non-mutually exclusive biological sources that explain the very high levels of gene tree discordance in our dataset: introgression, ILS, and gene (or genome) duplication. Despite a genome-wide phylogenomic dataset, we are unable to pinpoint exactly when and where along the phylogeny each of these processes has occurred. However, our resulting phylogenetic framework provides a robust starting point from which to understand the non-bifurcating nature of diversification of this Andean shrub clade.

Incomplete lineage sorting is common in rapid Andean radiations (Morales-Briones et al. 2018, Murillo-A. et al. 2022). Within *Freziera*, the Candicans group, especially its substituent Calophylla clade, carries the hallmarks of ILS due to rapid radiation, including conflicting species relationships with short branch lengths between close relatives (Fig 3a; Supplementary Fig. S4). Not only is gene tree-species tree discordance higher in this clade compared to other regions of *Freziera*’s phylogeny, but species relationships also differ across analyses (Supplementary Fig. S4). Further indication of ILS in this clade is the fact that the Calophylla clade is the most widespread of *Freziera*’s subclades and includes species with some of the broadest distributions in the genus. These distributional patterns allow a greater possibility that ancestral populations were large and widespread, contributing to ILS, amplifying the effect of short times between speciation events in this group.

Even after applying data curation that may inflate support for bifurcating relationships, a strong signal of introgression was identifiable in *Freziera*. This was evidenced by multiple significant f4 and D statistics (Supplementary Fig. S5). Due to the limited number of unlinked SNPs in our target capture data and the correlation of D and f4 introgression statistics when trios share internal branches, we were not able to isolate the exact branches along which introgression has occurred— a challenge even among the most complete datasets (Tricou et al. 2022). However, our resulting phylogenetic framework provides a robust starting point from which to understand the non-bifurcating nature of *Freziera*’s diversification. Areas of conflict across species trees in our curated datasets are strong candidates for lineages that have been directly shaped by past introgression (Fig. 3, Supplementary Fig. S4). A species that is particularly promising for future investigation is *F. magnibracteolata*, whose unstable placement along the backbone of *Freziera* (Table 1. Fig. 3; Supplementary Figs. S2,S4) suggests a potential history of introgression (MacGuigan and Near 2019; Cai et al. 2021). Notably, the placement of this species was the only major relationship to be impacted by removing artifactual orthologs relative to the unfiltered dataset (Table 1, *Monophyletic Elaphoglossifolia group*). To further assess the extent of gene flow and identify the branches involved in introgression events in this rapid radiation, future research will target a much larger portion of the genome (Malinsky et al. 2021) and include deeper taxon with multiple individuals per species.

Finally, either gene or genome duplication has resulted in paralogs in *Freziera*. We identify both paralogs for which multiple copies are identifiable during assembly and artifactual orthologs that are represented by only a single copy in our dataset (Fig. 2b, Supplementary Table S3). While the processes that gave rise to these paralogs are yet to be examined in detail, it is likely that they are product of allopolyploidy, especially considering the extensive history of genome duplication via polyploidy in Ericales, the order to which *Freziera* belongs (Larson et al. 2020), as well as chromosome count variation within Pentaphylacaceae (Rice et al. 2015).

## Conclusion

A major current challenge in phylogenomics is the difficulty in teasing apart specific sources of gene tree discordance in empirical datasets, and to account for these in phylogenetic inference (Morales-Briones et al. 2022; Tricou et al. 2022). It is a significant challenge to accurately identify paralogs, pinpoint specific instances of introgression, disentangle incomplete lineage sorting from historical gene flow, and reduce the impact of gene tree estimation error in a single empirical phylogenomic dataset in which all of these sources of discordance are present. In addition to these challenges, phylogenomic data have the potential to be very complex, particularly for clades that are well-understood to be recalcitrant like Andean plant radiations (Pease et al. 2016; Vargas et al. 2017). Here, we showed that careful data curation allowed us to detect a high proportion of artifactual orthologs, which we were able to reduce with multiple, non-mutually exclusive methods: heeding paralog warnings, removing gene trees with a high proportion of heterozygous sites, and filtering gene trees using bipartition support. These data curation strategies were subsequently associated with higher support, lower gene tree conflict, and a more stable species tree— the first for an understudied tropical plant clade that previously lacked any phylogenetic information. We advocate for the observation of empirical phylogenomic data, including gene tree alignments and topologies, and that data curation be tailored to unique properties of individual datasets to better address the above-mentioned complexities in phylogenetic inference.

While commonly used filtering techniques, assembly parameters, and other automated aspects of phylogenomics are powerful tools to improving phylogenetic inference, we have shown that they can also increase the proportion of data artifacts (i.e., artifactual paralogs) and have negative impacts on phylogenetic support and inference. Automated filtering techniques are not a replacement for a deep understanding of a dataset. Selecting filtering strategies for individual datasets should be informed by the latter since the decision is likely to be a balance between minimizing the presence of data artifacts while maximizing the number of loci useful for phylogenetic inference. While targeted sequence capture of universal loci offers potential, especially for phylogenetic studies relying heavily on DNA from natural history collections, these datasets are not without limitations related to the nature of the data themselves and to the algorithms we use to process and analyze them. Combining exploration of datasets with deep knowledge of the organismal biology of targeted clades is crucial toward overcoming these limitations and inferring robust phylogenetic hypotheses.

## Data Availability

All data generated in this study are publicly accessible. Raw sequence reads are deposited on NCBI’s Sequence Read Archive: (*web address added upon acceptance*). Supplementary materials cited in this paper and processed data are available from the Dryad Digital Repository (https://datadryad.org/stash/share/X6HXB23fQmPqODdRRdXNJnGePrw29vUPbm1JgIijjFo).

## Acknowledgements

This research was funded by a Louisiana Board of Regents Research Competitiveness Subprogram grant, the LSU College of Science and Office of Research and Economic Development, and NSF Award DEB-2055525. We would like to thank the Missouri Botanical Garden (MO) for their access to their collections. We thank Brant Faircloth, Matthew Johnson, Carl Oliveros, and Jessie Salter for their guidance in library preparation, and Brant Faircloth for access to laboratory equipment. Computational analyses were performed on LSU High Performance Computing’s SuperMike cluster. We thank three anonymous reviewers for their comments and criticism, which greatly improved previous versions of this work. This manuscript benefited from feedback from Laymon Ball, Janet Mansaray, and Diego Paredes-Burneo. The taxonomic expertise of Daniel Santamaría-Aguilar was critical throughout the design and implementation of this research.

Online Appendix 1. Supplementary Materials and Methods

Online Appendix 2. Paralogs identified by HybPhaser for all samples and for each sample.

Supplementary Figure S1. Heat map showing gene recovery efficiency for all submitted samples (rows) across the 353 targeted genes (columns).

Supplementary Figure S2. ASTRAL-III species trees for the 11 datasets: a) *orthologs.unfiltered*, b) *orthologs.by.eye*, c) *orthologs.HybPiper.long*, d) *orthologs.HybPiper.no.warnings*, e) *HybPhaser*, f) *MO*, g) *proportion.PI*, h) *proportion.internal.branch.lengths*, i) *average.BS*, j) *tree.length*, k) *bipartition*

Supplementary Figure S3. Number of reads mapped to target regions in silica (blue) and herbarium (red) samples. There is a correlation between the number of reads mapped and the number of paralog warnings from HybPiper in a sample (R^2^ = 0.55, F(df=1) = 123.6, p <0.0001). Assembly and paralog detection are improved with increased sequencing effort.

Supplementary Figure S4. Tanglegram plots showing pairwise comparisons between ASTRAL-III species tree topologies for a) *consensus* vs. *orthologs.unfiltered*, b) *consensus* vs. *orthologs.by.eye*, c) *consensus* vs. *orthologs.HybPiper2.long*, d) *consensus* vs. *orthologs.HybPiper2.no.warnings*, e) *consensus* vs. *HybPhaser*, f) *consensus* vs. *MO*, g) *consensus* vs. *proportion.PI*, h) *consensus* vs. *proportion.internal.branch.lengths*, i) *consensus* vs. *average.BS*, j) *consensus* vs. *tree.length*, k) *consensus* vs. *bipartition*, l) *orthologs.by.eye* vs. *orthologs.unfiltered*, m) *orthologs.by.eye* vs. *orthologs.HybPiper2.long*, n) *orthologs.by.eye* vs. *orthologs.HybPiper2.no.warnings*, o) *orthologs.by.eye* vs. *HybPhaser*, p) *orthologs.by.eye* vs. *MO*, q) *orthologs.by.eye* vs. *proportion.PI*, r) *orthologs.by.eye* vs. *proportion.internal.branch.lengths*, s) *orthologs.by.eye* vs. *average.BS*, t) *orthologs.by.eye* vs. *tree.length*, u) *orthologs.by.eye* vs. *bipartition*, v) *orthologs.HybPiper2.no.warnings* vs. *orthologs.unfiltered*, w) *orthologs.HybPiper2.no.warnings* vs. *orthologs.HybPiper2.long*, x) *orthologs.HybPiper2.no.warnings* vs. *HybPhaser*, y) *orthologs.HybPiper2.no.warnings* vs. MO, z) *orthologs.HybPiper2.no.warnings* vs. *proportion.PI*, aa) *orthologs.HybPiper2.no.warnings* vs. *proportion.internal.branch.lengths*, bb) *orthologs.HybPiper2.no.warnings* vs. *average.BS*, cc) *orthologs.HybPiper2.no.warnings* vs. *tree.length*, dd) *orthologs.HybPiper2.no.warnings* vs. *bipartition*, ee) *HybPhaser* vs. *orthologs.unfiltered*, ff) *HybPhaser* vs. *orthologs.HybPiper2.long*, gg) *HybPhaser* vs. MO, hh) *HybPhaser* vs. *proportion.PI*, ii) *HybPhaser* vs. *proportion.internal.branch.lengths*, jj) *HybPhaser* vs. *average.BS*, kk) *HybPhaser* vs. *tree.length*, ll) *HybPhaser* vs. *bipartition*, mm) *bipartition* vs. *orthologs.unfiltered*, nn) *bipartition* vs. *orthologs.HybPiper2.long*, oo) *bipartition* vs. *MO*, pp) *bipartition* vs. *proportion.PI*, qq) *bipartition* vs. *proportion.internal.branch.lengths*, rr) *bipartition* vs. *average.BS*, ss) *bipartition* vs. *tree.length*, tt) *orthologs.HybPiper2.long* vs. *orthologs.unfiltered*, uu) *MO* vs. *orthologs.unfiltered*, vv) *proportion.PI* vs. *orthologs.unfiltered*, xx) *proportion.internal.branch.lengths* vs. *orthologs.unfiltered*, yy) *average.BS* vs. *orthologs.unfiltered*, zz) *tree.length* vs. *orthologs.unfiltered*.

Supplementary Figure S5. Results of D and f4 statistics calculated with Dsuite on the “bipartition support”, “by eye”, and “HybPiper2 no warnings" datasets. The columns indicate 1) values of Patterson’s D-statistic for evaluated trios, 2) highly significant p-values associated with the D-statistics after Benjamini-Hochberg correction, and 3) f4 ratio statistics. Introgression is detected across all datasets even after assessing family-wise error rate.

Supplementary Figure S6. Scatterplot showing locus heterozygosity (proportion of loci with SNPs) versus allele divergence calculated with HybPhaser for all included accessions.

Supplementary Table S1. Per sample data--species name, voucher information and herbarium code for tissue gathered from specimens (codes follow Index Herbariorum: http://sweetgum.nybg.org/science/ih/), collection date, provenance, and sample ID used in phylogenetic analyses--and summary statistics generated with HybPiper v.1.3.1.

Supplementary Table S2. HybPiper v.2.0.1 assembly statistics for all samples included in this study. Paralog warnings per individual are shown.

Supplementary Table S3. Paralog identification results from the by-eye and automated strategies per locus. The proportion of stitched contigs, paralog warning by depth, and long paralog warnings from HybPiper v.2.0.1, paralog warnings from HybPiper v.1.3.1, removed loci with HybPhaser as well as per locus statistics for alignments and gene trees used to explore how standard gene tree filtering affects the presence of artifactual orthologs in our dataset are shown. Details of by-eye identification of paralogs, and the handling of reports from HybPiper v,2.0.1 are in Appendix 1.

Supplementary Table S4. HybPiper v.2.0.1 paralog report from assembly using a taxon-specific target file generated with Easy353. Reference sequences were extracted from transcriptome data of Ternstroemia gymnanthera (Pentaphylacaceae). Numbers on the table correspond to the number of contigs assembled per locus.

Supplementary Table S5. Per sample summary statistics from HybPhaser, including the recovered sequence length (bp), the recovered sequence length as the proportion of the target sequences from HybPiper (bpoftarget), number of paralogs found in all samples, number of additional paralogs found for each sample, number of loci recovered, allele divergence, locus heterozygosity, loci with >0.5% SNPs, loci with >1% SNPs, loci with >2% SNPs

