## Supplementary material for "Strong phylogenetic signal despite high phylogenomic complexity in an Andean plant radiation (*Freziera,* Pentaphylacaceae)": Online Appendix 1

Online Appendix 1. Supplementary Materials and Methods

*DNA extraction, library prep, target enrichment, and sequencing*

Five hundred mg of dried leaf tissue was homogenized using a FastPrep-24TM 5G bead beating and lysis system (MP Biomedicals, Solon, Ohio, United States). DNA extraction followed a modified sorbitol extraction protocol (Štorchová et al. 2000). To reduce the risk of cross-contamination between samples, tissue from each sample was weighed on disposable weigh paper, and tissue sampling and DNA extractions were performed in small batches of 12 samples at a time. Forceps used to break off pieces of leaf tissue for extraction were cleaned with 70% ETOH between samples, and filter tips were used during all laboratory work. Double-stranded DNA concentration was quantified using a Qubit 4 Fluorometer (Invitrogen, Waltham, Massachusetts, United States) and fragment size was assessed on a 1% agarose gel. For samples with a high concentration of large fragments (>800 bp), DNA was sheared using a Bioruptor Pico (Diagenode Inc., Denville, New Jersey, United States) until most fragments were less than 500 bp in length.

Library preparation with KAPA Hyper Prep followed the manufacturer’s protocol (KR0961 – v8.20) with the following modifications: reaction volumes were halved (i.e., 25 μL starting reaction) and bead-based clean-ups were performed at 3X volume rather than 1X volume to preserve more small fragments from degraded samples that are characteristic of herbarium specimens. As the 3X volume bead-based clean-up retains adapter dimers as well as short fragments, samples were visualized again using a 1% agarose gel to identify samples with an abundance of fragments shorter than 100 bp. Those samples were processed with a GeneRead Size Selection kit to remove fragments shorter than 150 bp (Qiagen, Germantown, Maryland, United States). Library amplification reactions were performed at 50 μL. Library preparation and amplification was, again, performed in small batches using filter tips to prevent large-scale contamination of samples.

Target enrichment followed the modifications to the manufacturer’s protocol outlined in (Hale et al. 2020); i.e., pools of 20-24 samples and RNA baits diluted to ¼ concentration). Twenty nanograms of unenriched DNA library were added to the cleaned, target enriched pool to increase the amount of off-target, chloroplast fragments in the sequencing library.

*Raw data processing and locus extraction and alignment cleaning*

For version HybPiper v1.3.1, read mapping and contig assembly were performed using the reads_first.py script. The intronerate.py script was run to extract introns and intergenic sequences flanking targeted exons. Coding and non-coding regions were extracted using the retrieve_sequences.py script with “dna” and “supercontig” arguments, respectively. Supercontigs include both coding and non‐coding regions as a single concatenated sequence for each target gene. Individual genes were aligned using MAFFT v. 7.310 (Katoh and Standley 2013). Loci flagged as paralogous by the paralog_retriever.py script in HybPiper v. 1.3.1 were removed from downstream analyses.

*Paralog warnings and detection of artifactual orthologs*

HybPiper v.2.0.1 provides two new reports for users for each sample: (1) a file with “Yes” or “No” as to whether a stitched contig was produced and (2) a file with “True” or “False” as to whether a paralog warning by contig depth was issued for each gene with sequences. These two types of reports were collected across samples and compiled into a single document for each. The proportion of “Yes” results were calculated for each locus to determine the frequency of stitched contigs in our dataset, and the proportion of “True” results were calculated for each locus to identify additional putative paralogs.

*Phylogenetic analyses*

*Gene tree inference and filtering*— Preliminary gene trees were generated from aligned sequences for the 322 loci lacking paralog flags with RAxML v8.2.12 (Stamatakis 2014) under the GTR model with optimization of substitution rates and site-specific evolutionary rates (-m GTRCAT) and 200 rapid bootstrap replicates. The preliminary trees were then processed with TreeShrink v1.3.3 [(Mai and Mirarab 2018)](https://pa) on a “per-gene” and “all-gene” basis to identify long branches that are likely associated with spurious sequences. The “per-gene” test identifies exceedingly long branches from the distribution of signature values (i.e., the maximum reduction in tree diameter resulting from removal of a set of terminal branches) within each gene, whereas the “all-gene” creates one distribution based on all genes to which species signatures are compared (Mai and Mirarab 2018). The identified samples were removed from alignments, except in instances where the entire *Ternstroemia* clade in the outgroup was identified, as this was considered more likely to reflect clade-specific differences in mutation rate and/or time elapsed since MRCA than spurious alignments.
