## Supplementary figures and images for "Strong phylogenetic signal despite high phylogenomic complexity in an Andean plant radiation (*Freziera,* Pentaphylacaceae)"

### _a_consensus_vs_unfiltered.pdf

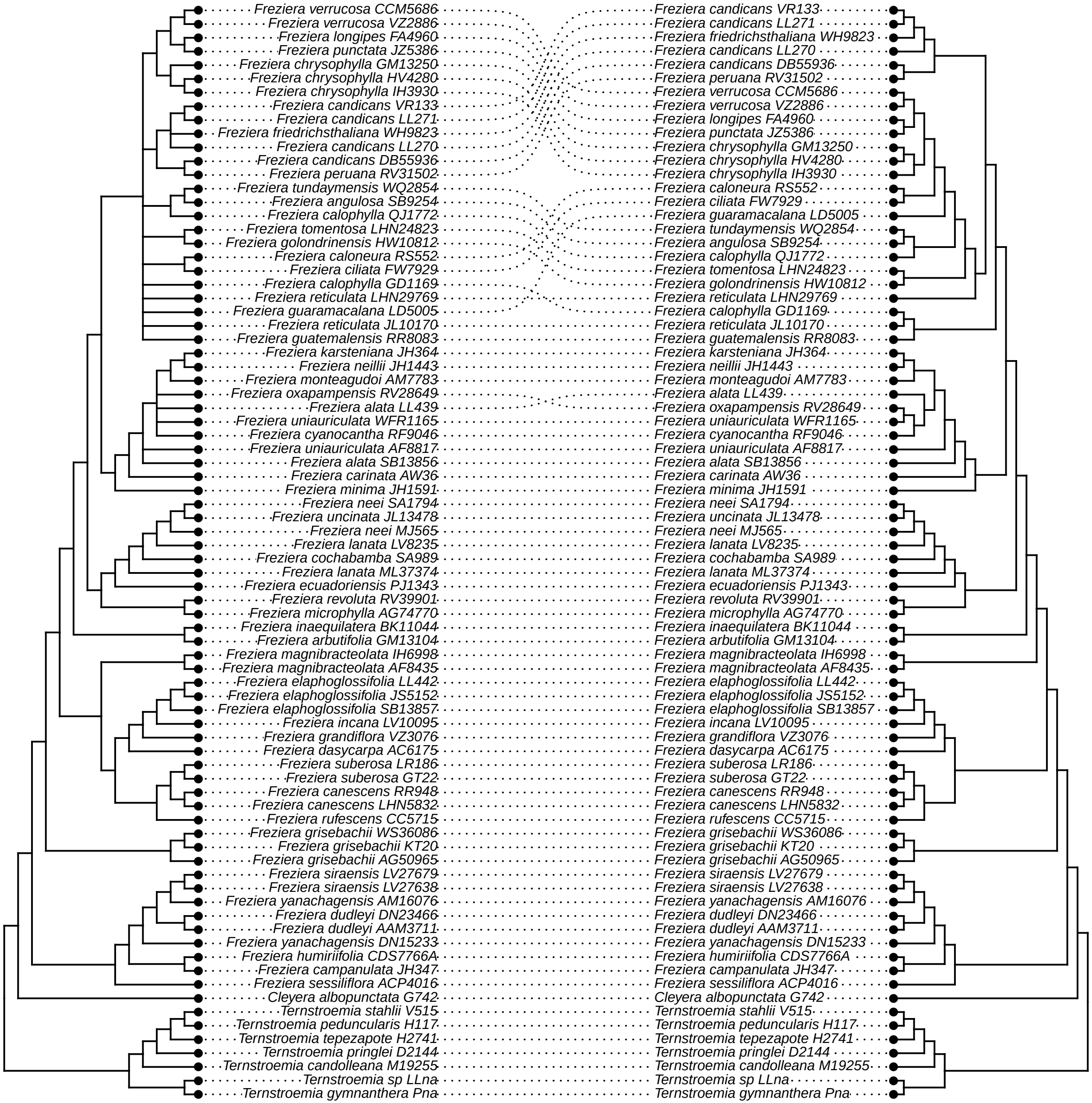

### _b_consensus_vs_by_eye.pdf

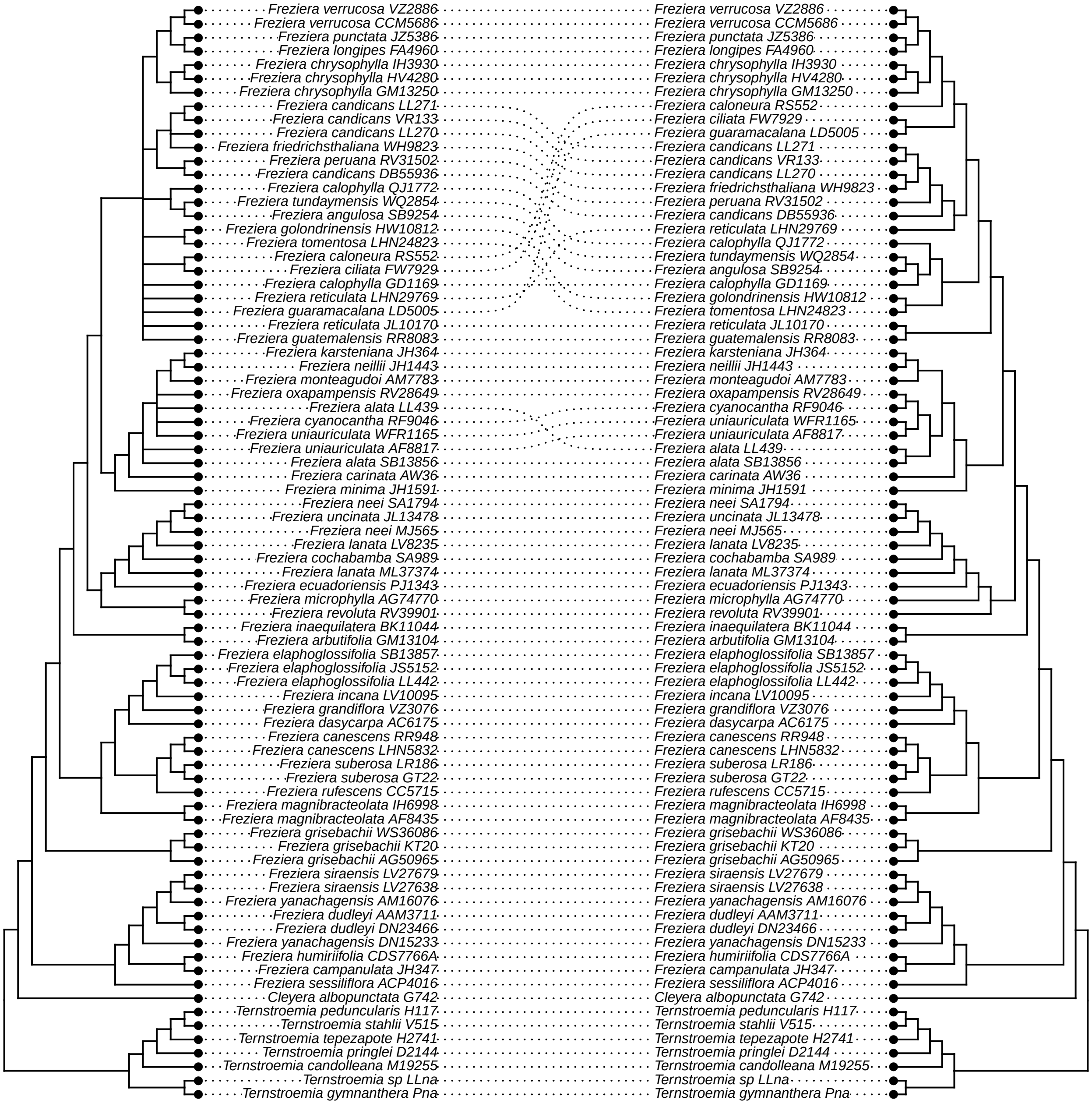

### _c_consensus_vs_hp2_long.pdf

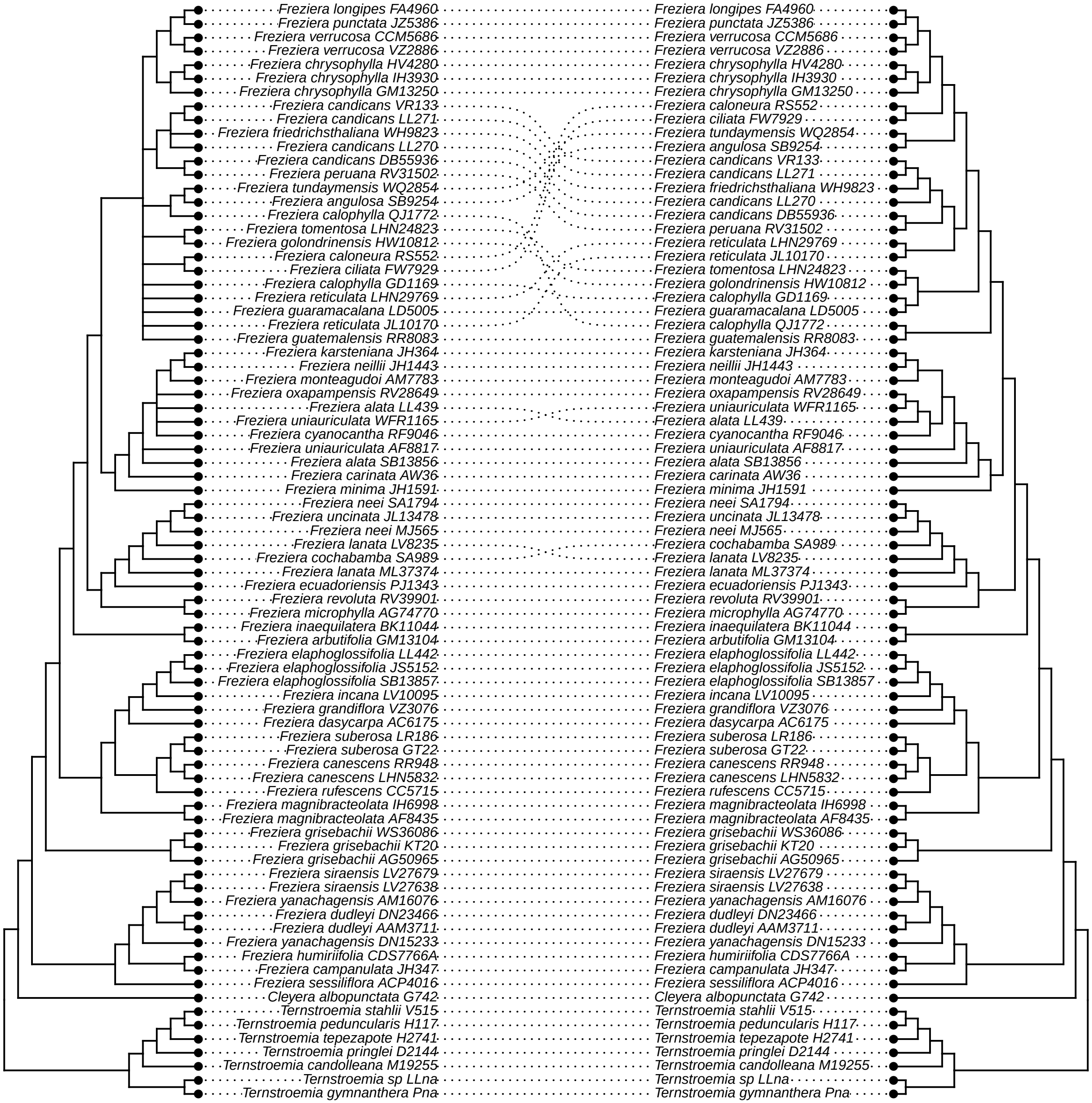

### _d_consensus_vs_hp2_no_warnings.pdf

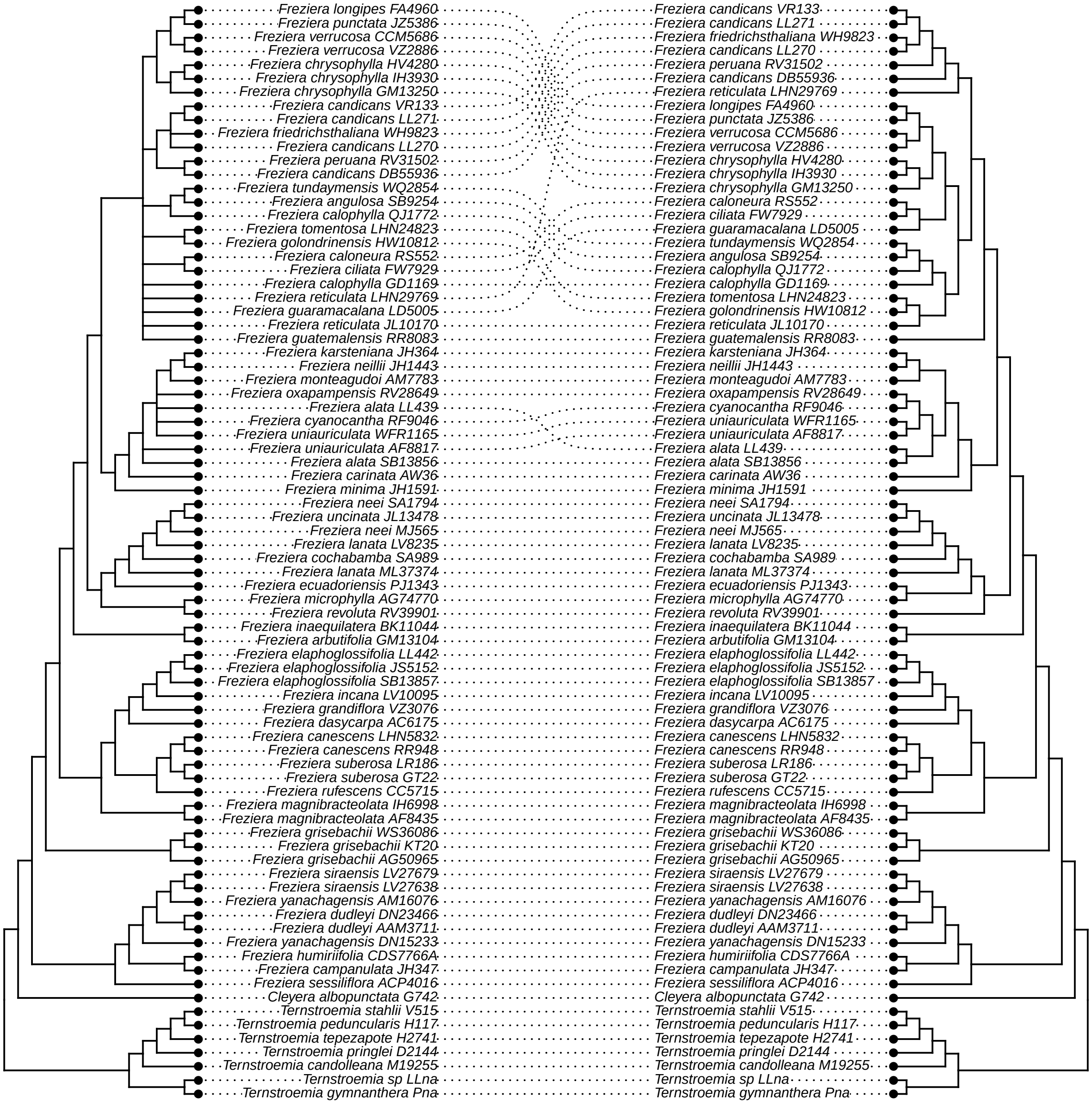

### _e_consensus_vs_HybPhaser.pdf

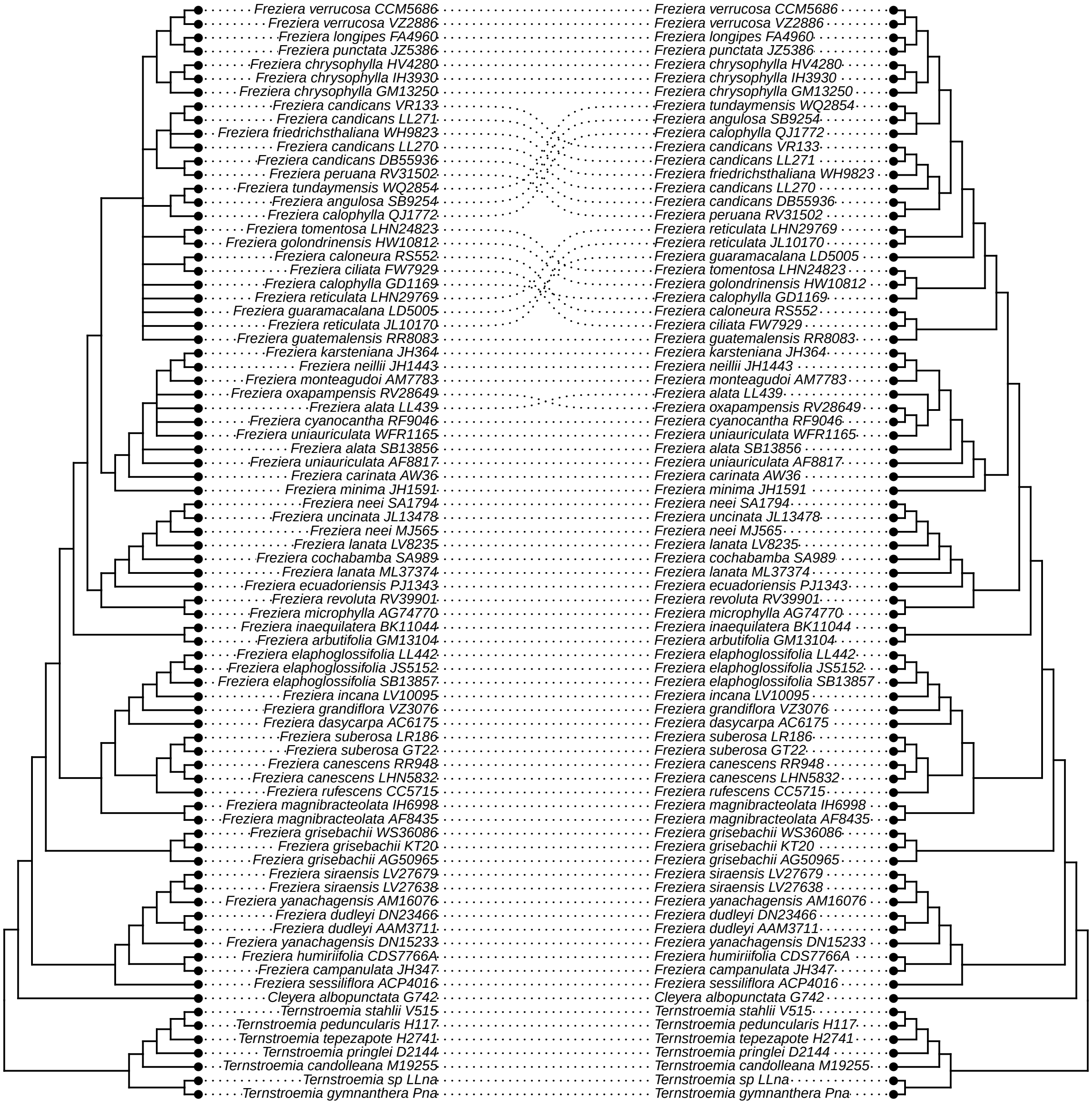

### _f_consensus_v_MO.pdf

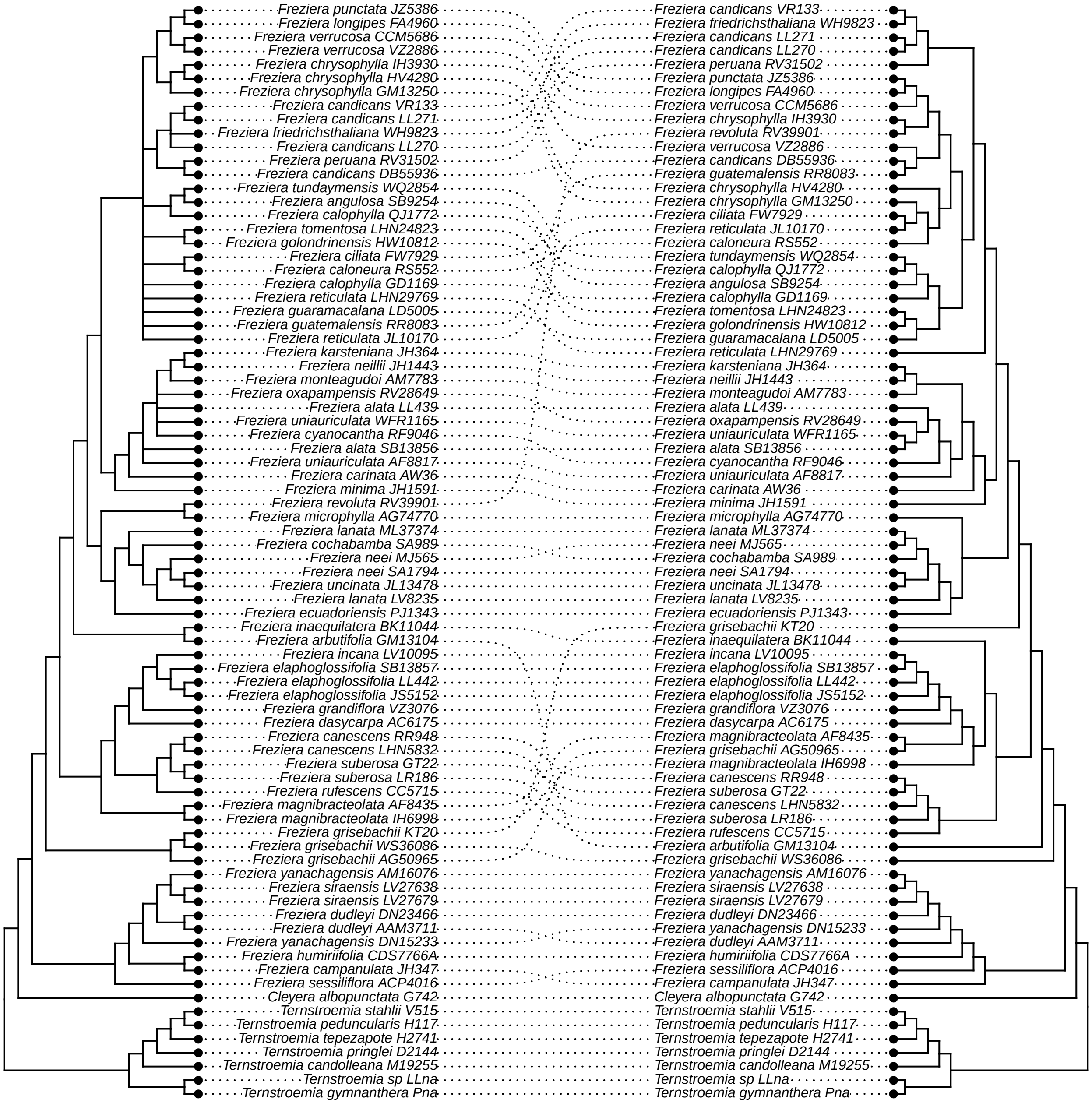

### _g_consensus_v_prop_PI.pdf

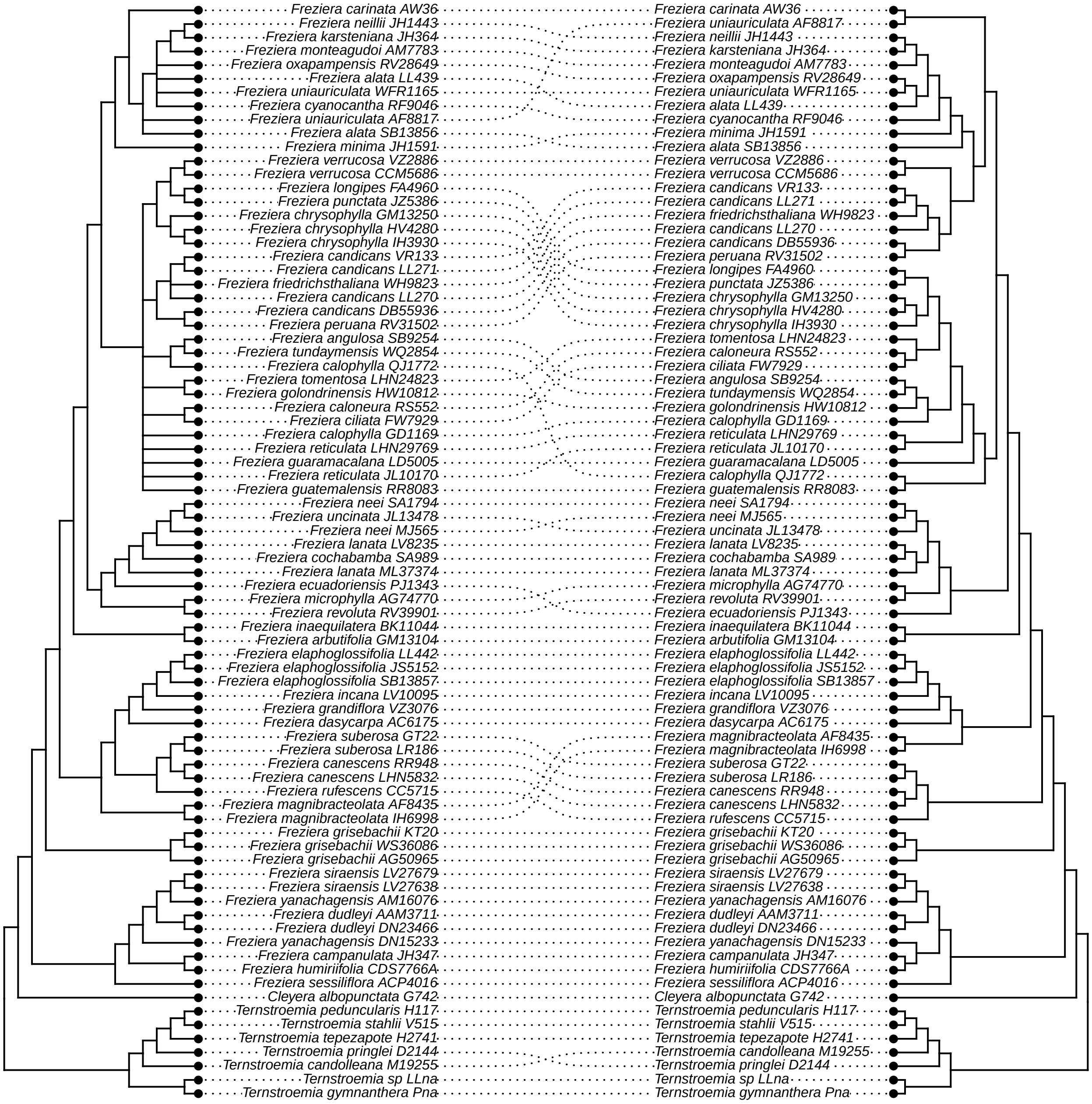

### _h_consensus_v_internal.pdf

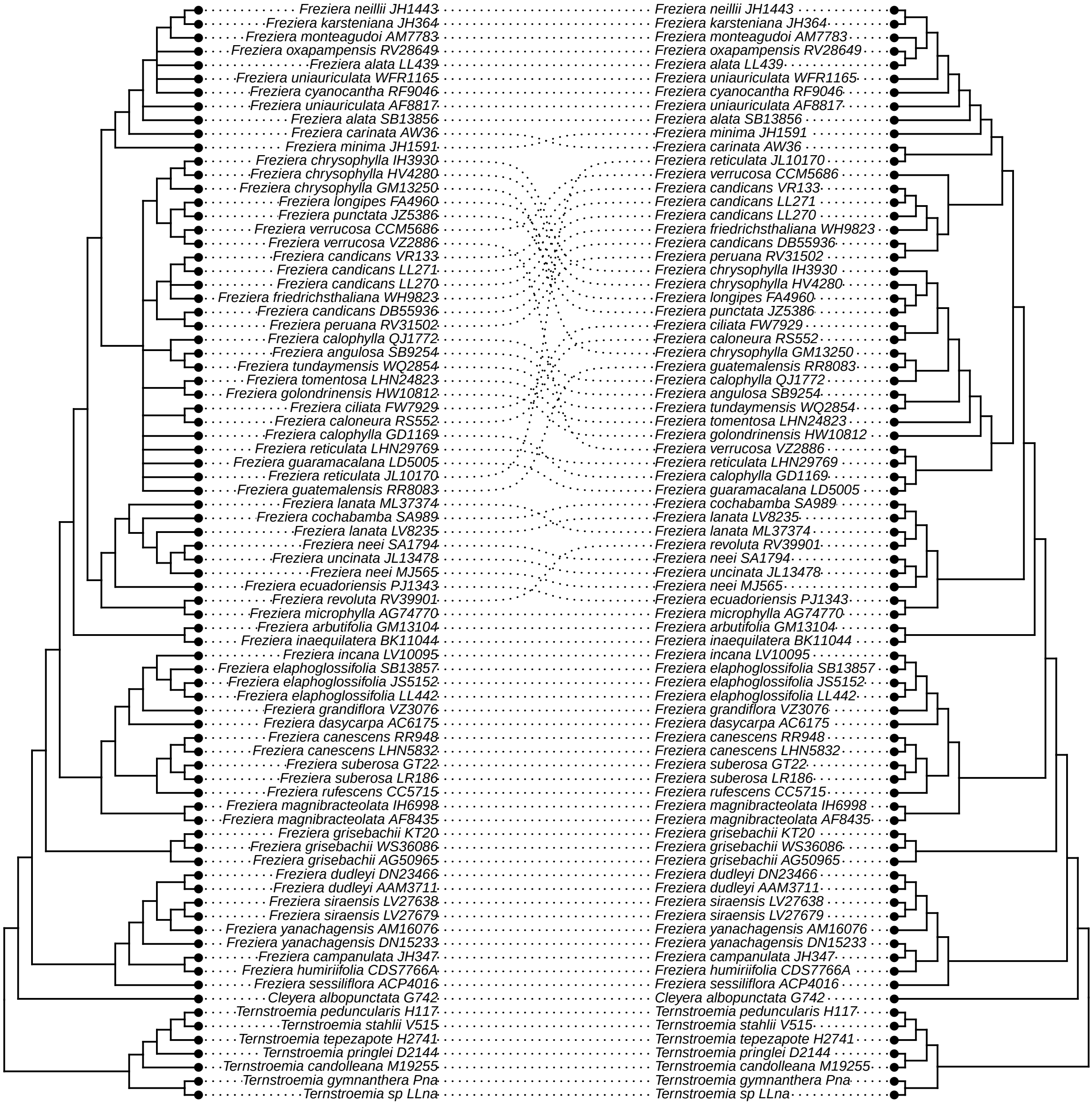

### _i_consensus_v_avg_BS.pdf

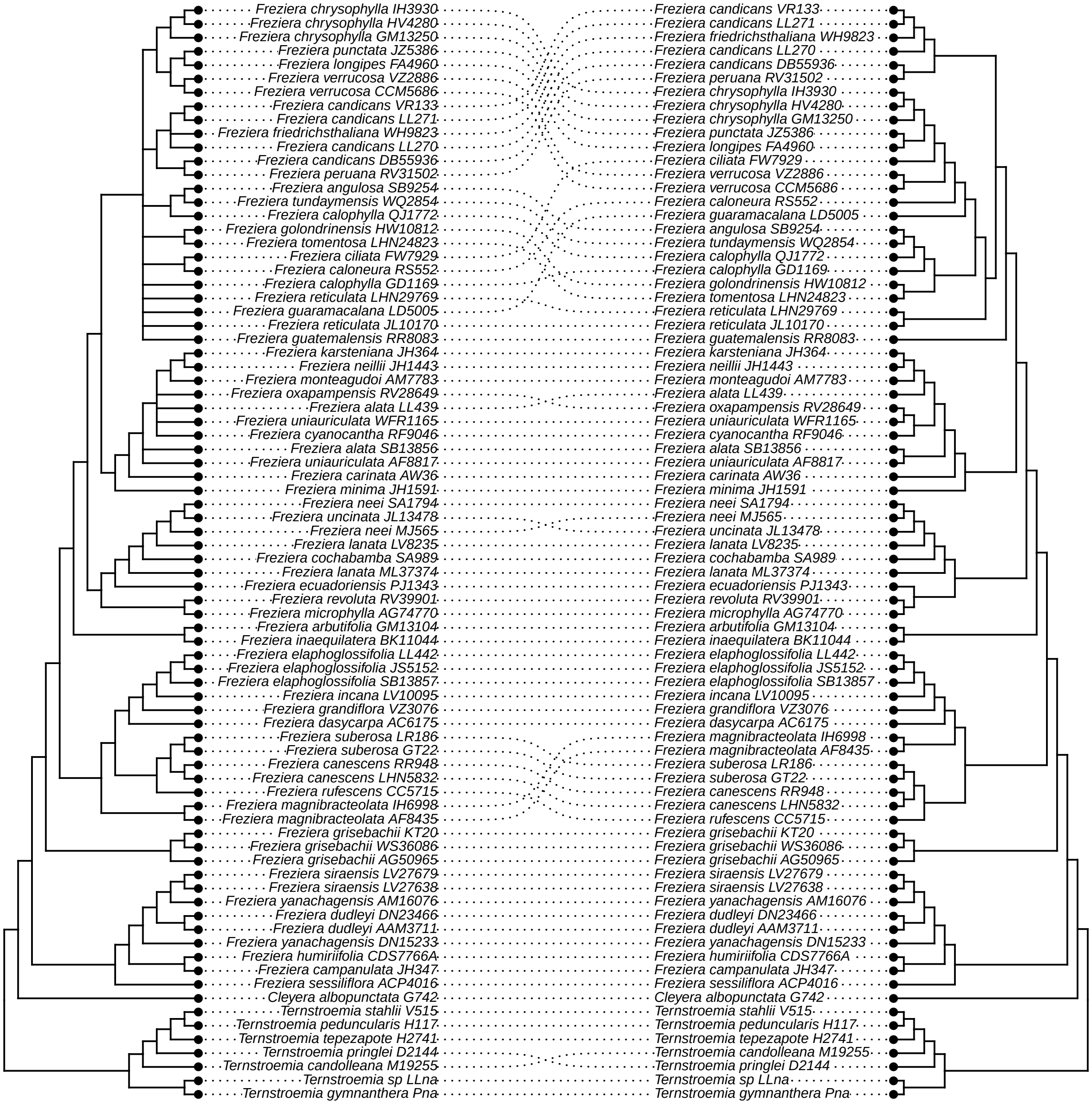

### _j_consensus_v_tree_length.pdf

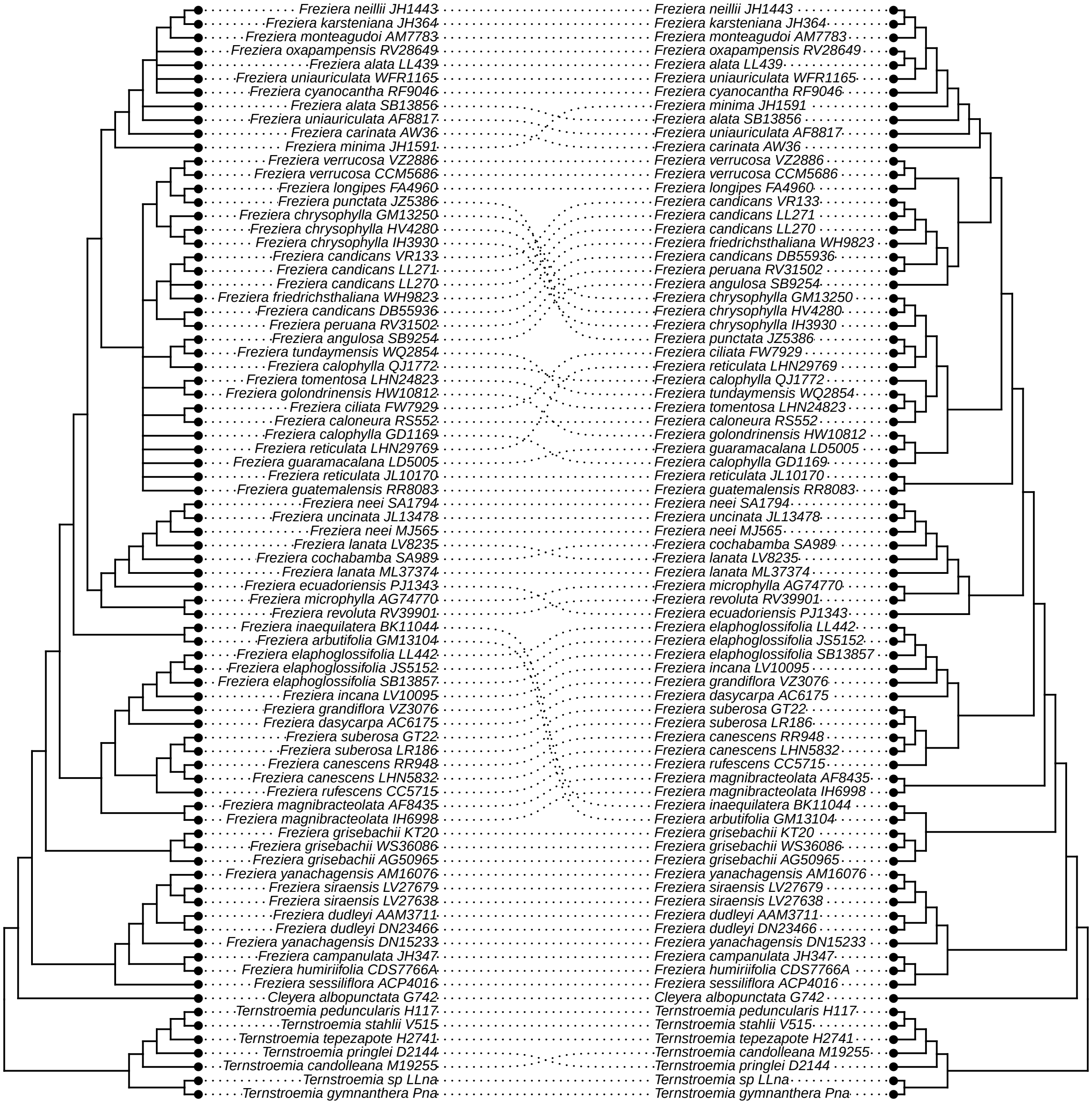

### _k_consensus_vs_bipartition.pdf

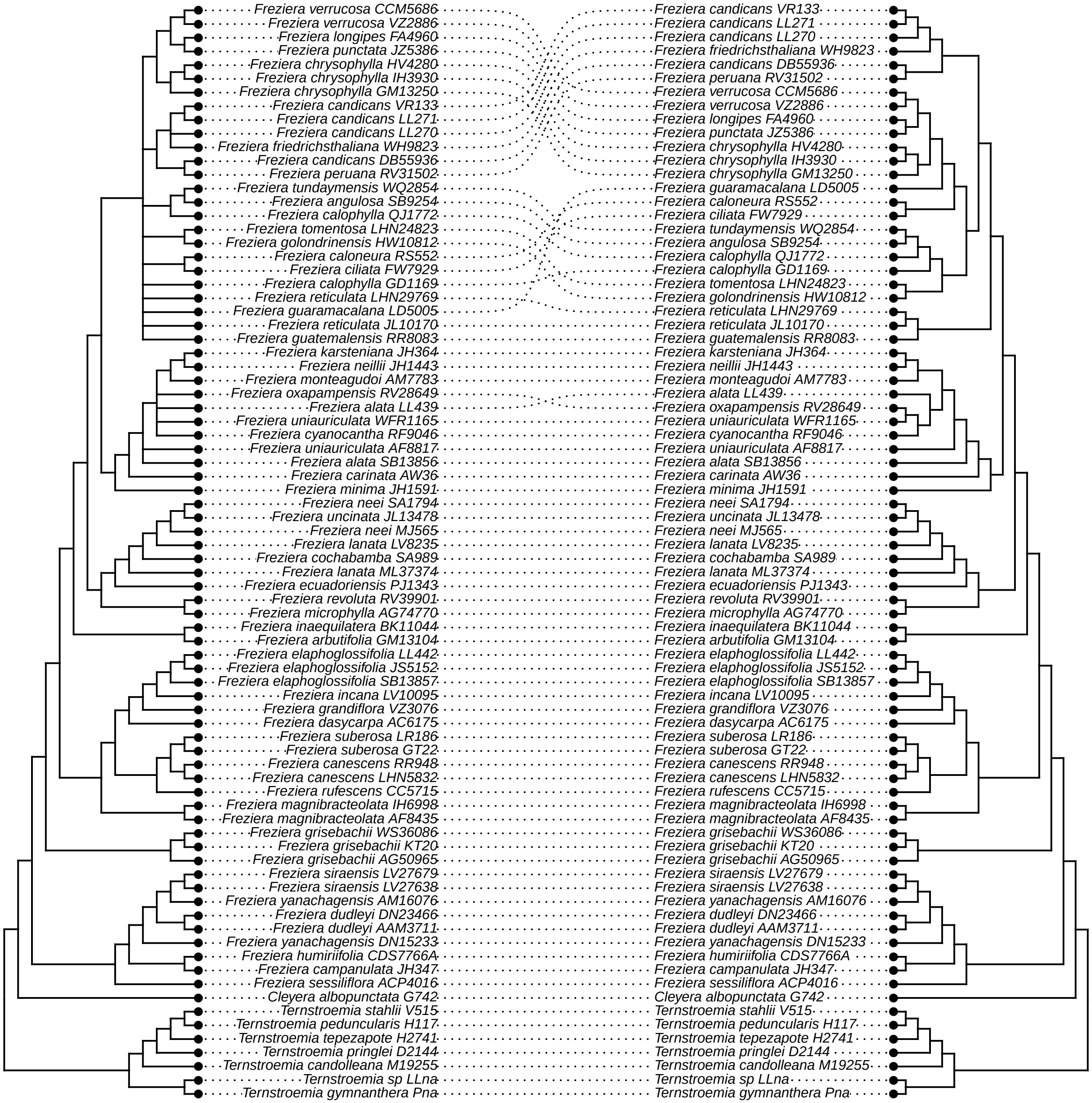

### _l_by_eye_vs_unfiltered.pdf

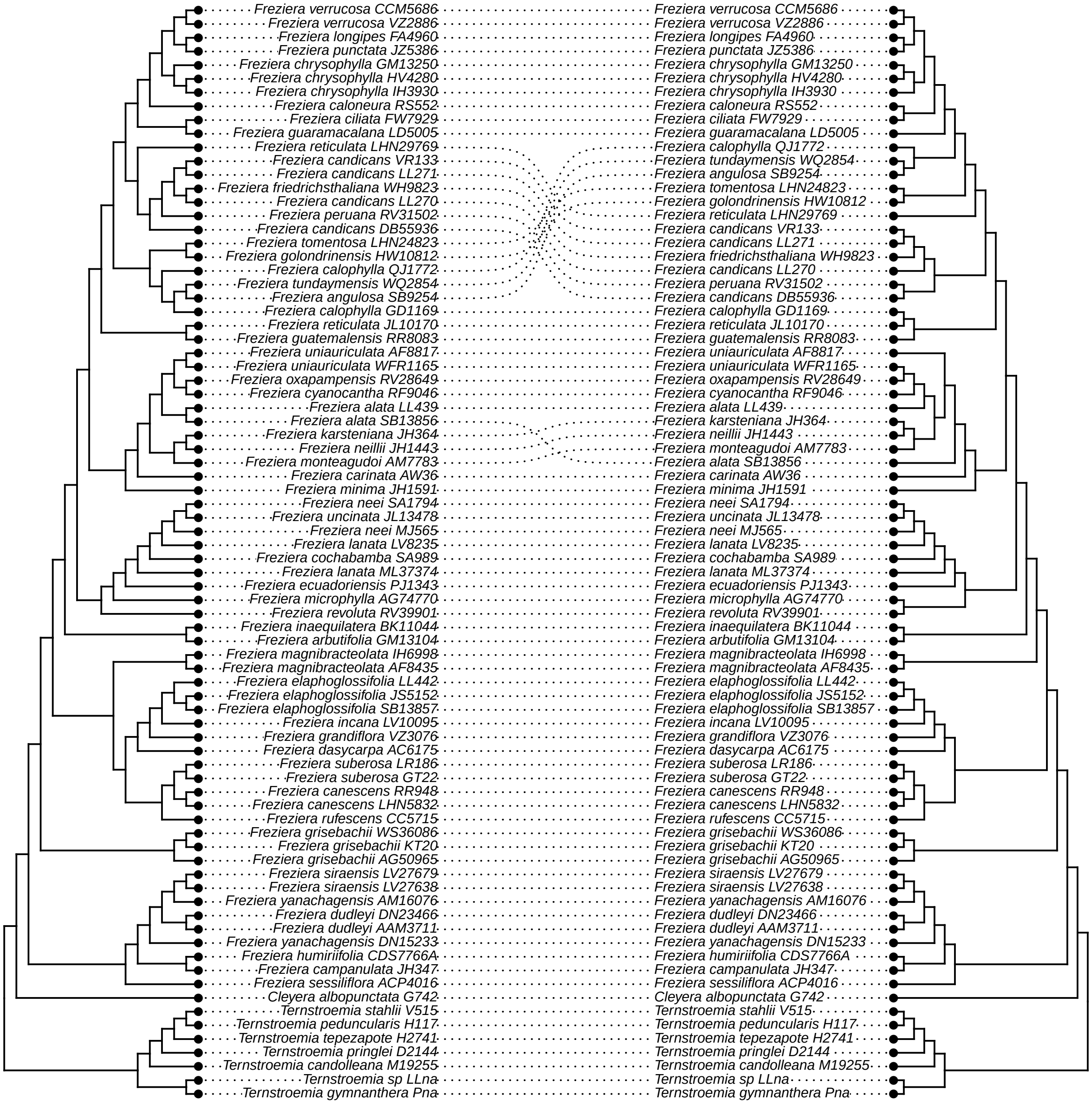

### _m_by_eye_vs_hp2_long.pdf

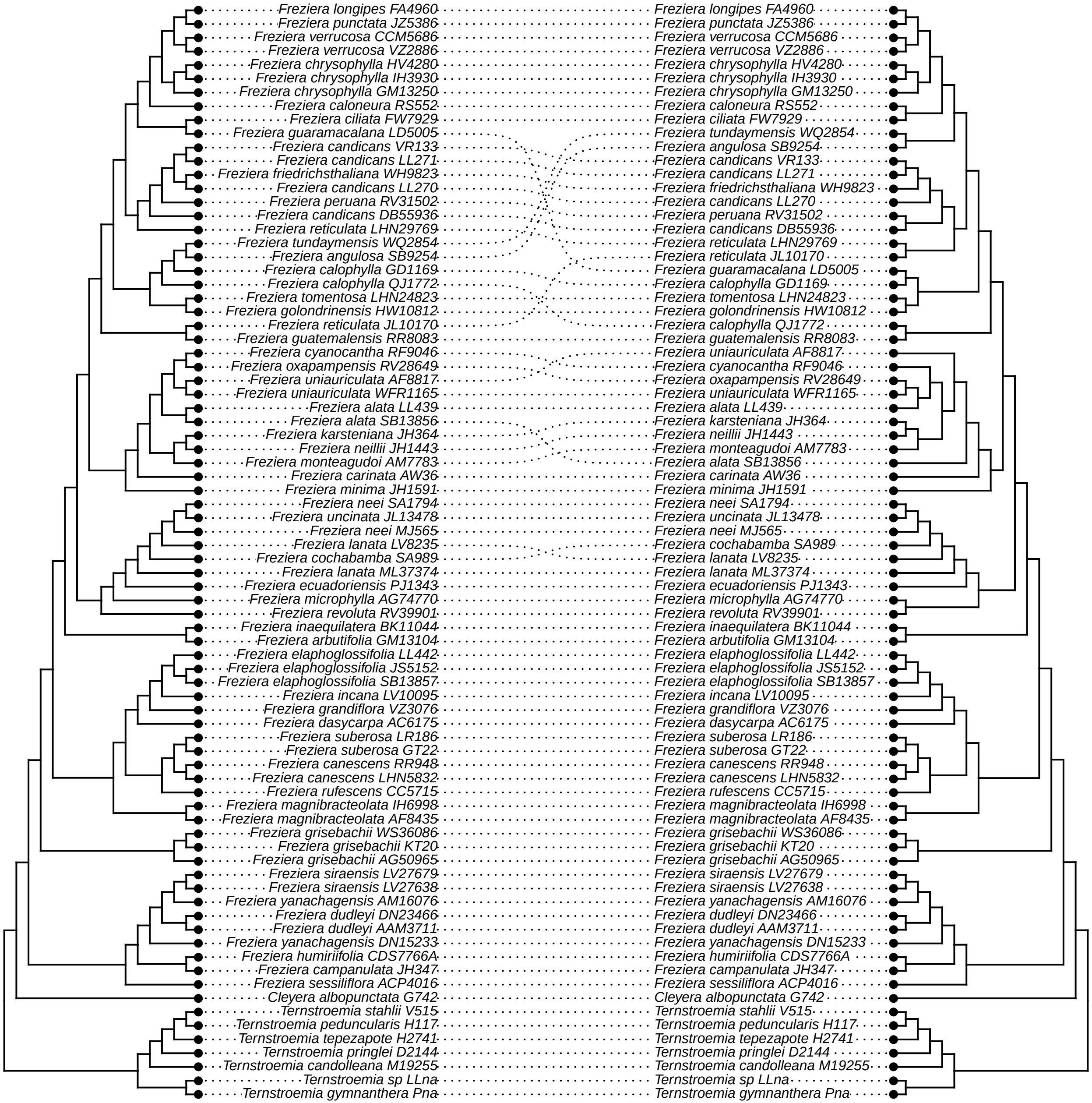

### _n_by_eye_vs_hp2_no_warnings.pdf

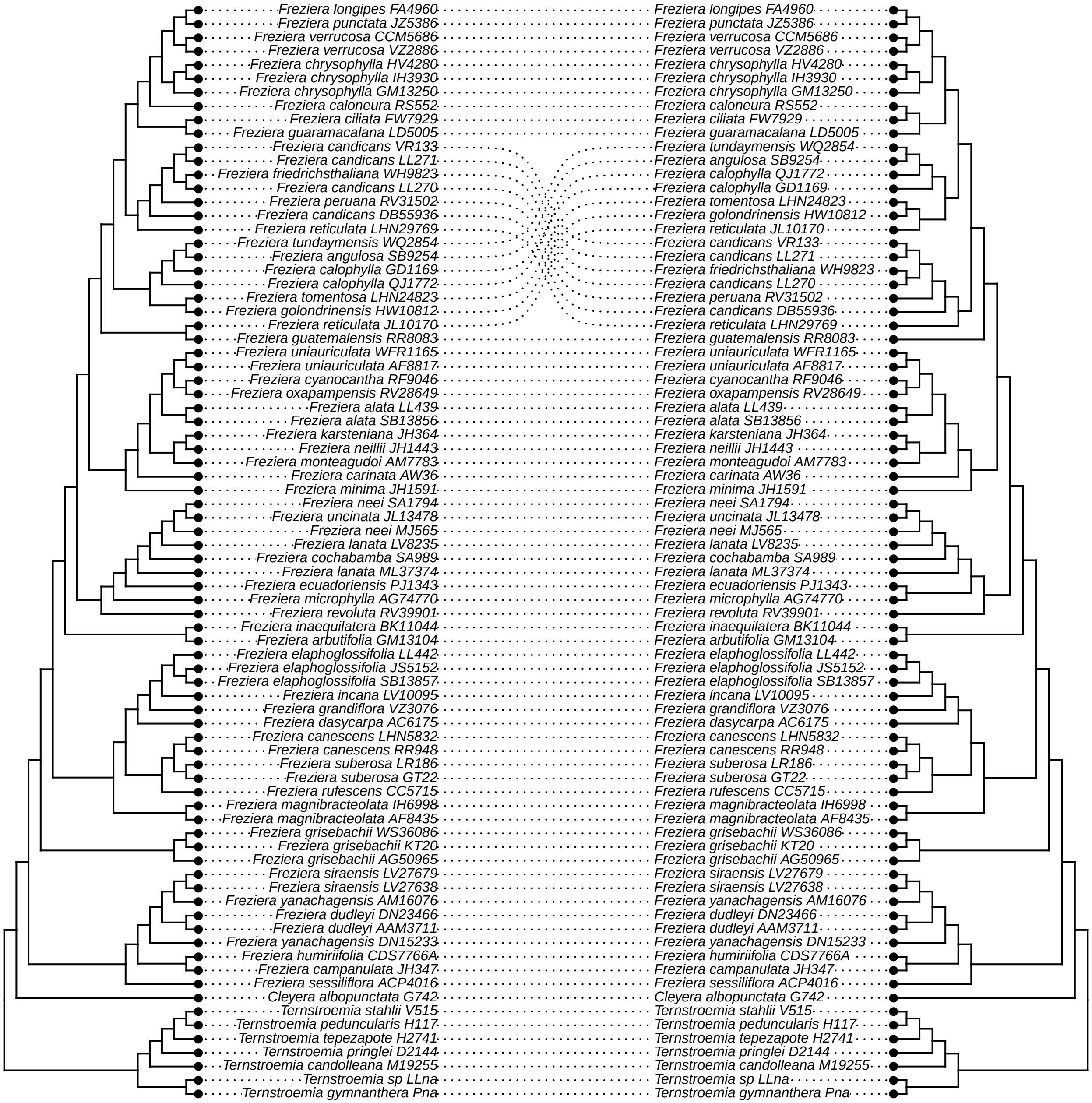

### _o_by_eye_vs_HybPhaser.pdf

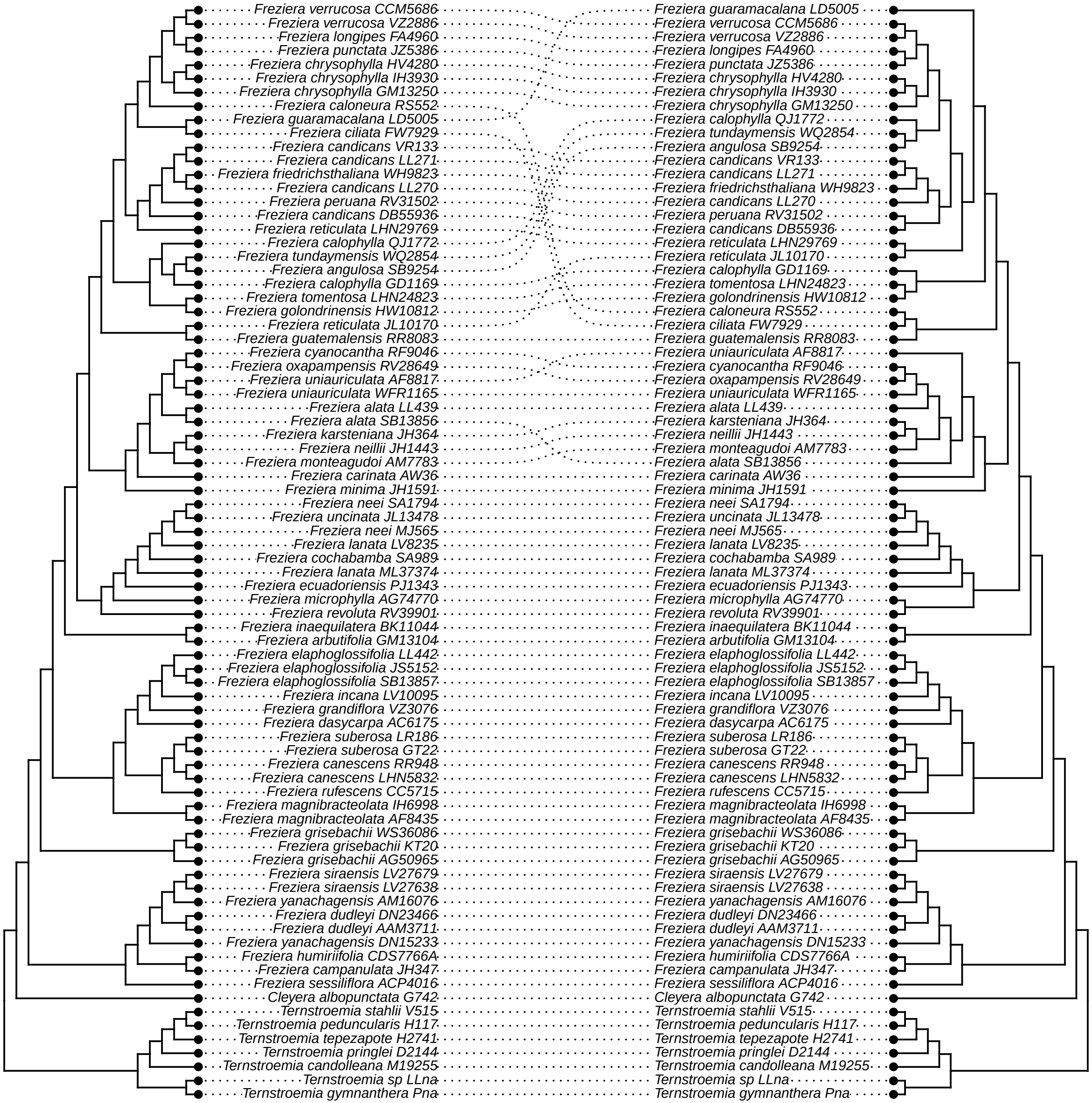

### _p_by_eye_v_MO.pdf

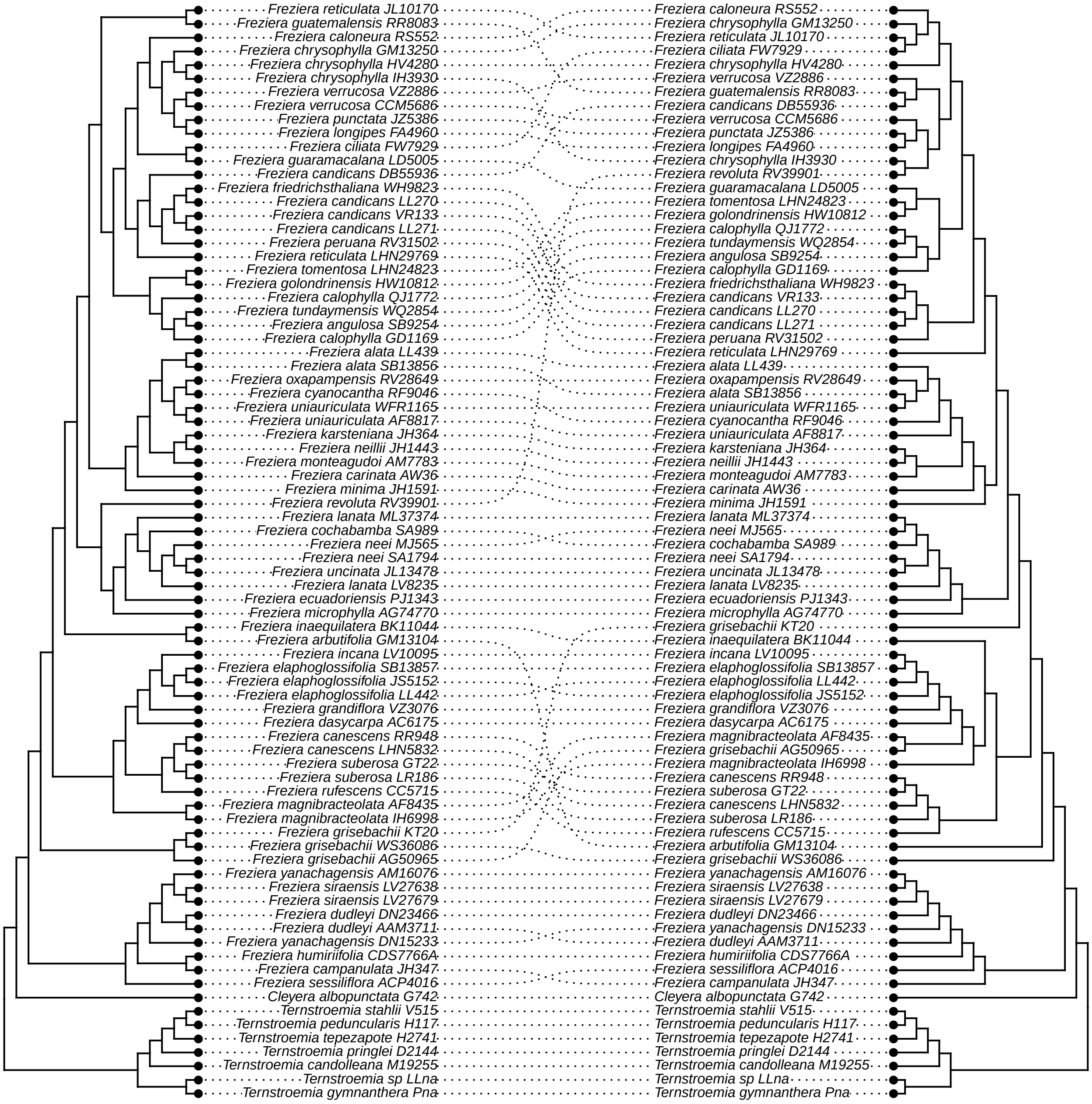

### _q_by_eye_v_prop_PI.pdf

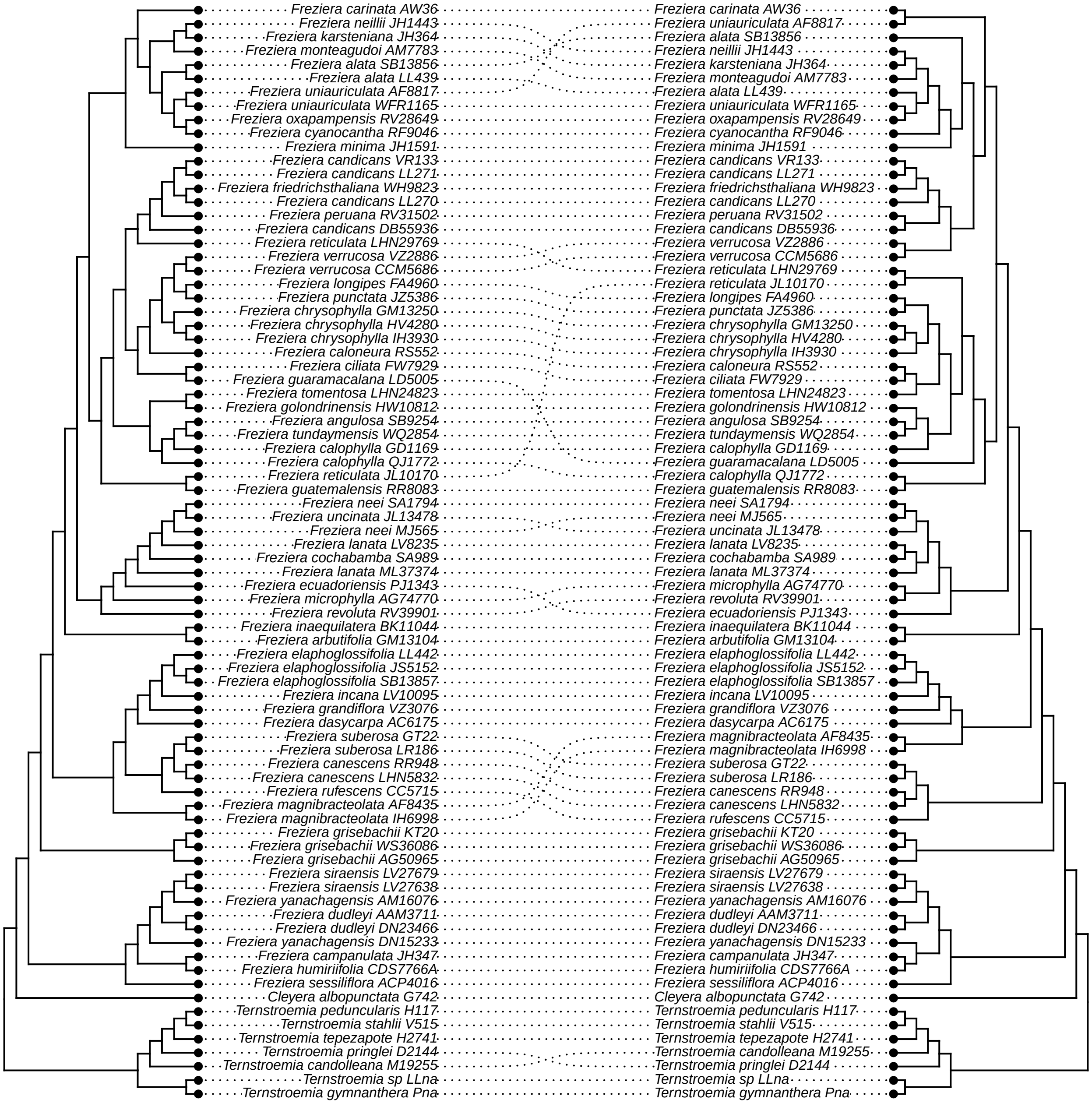

### _r_by_eye_v_internal.pdf

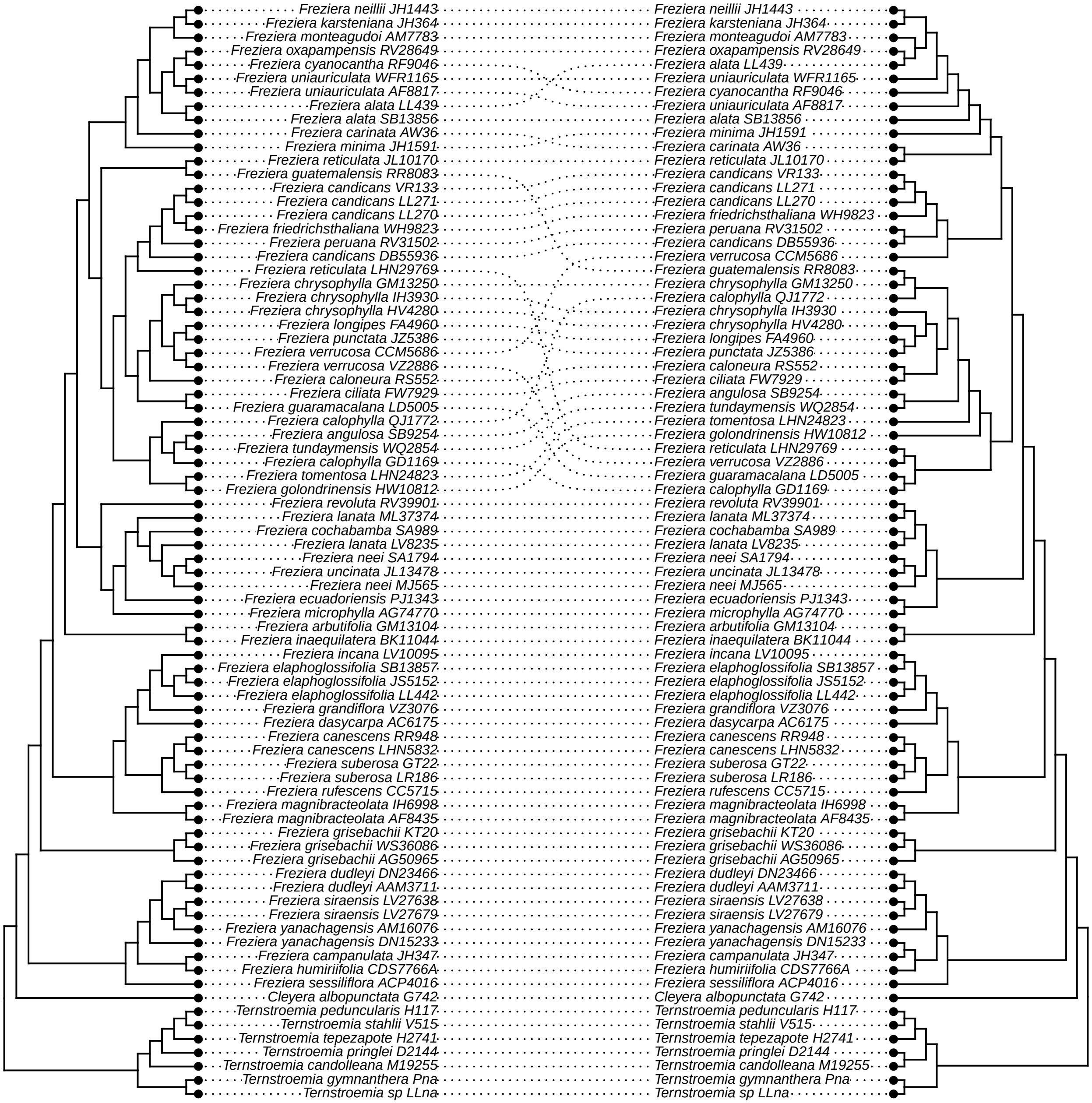

### _s_by_eye_v_avg_BS.pdf

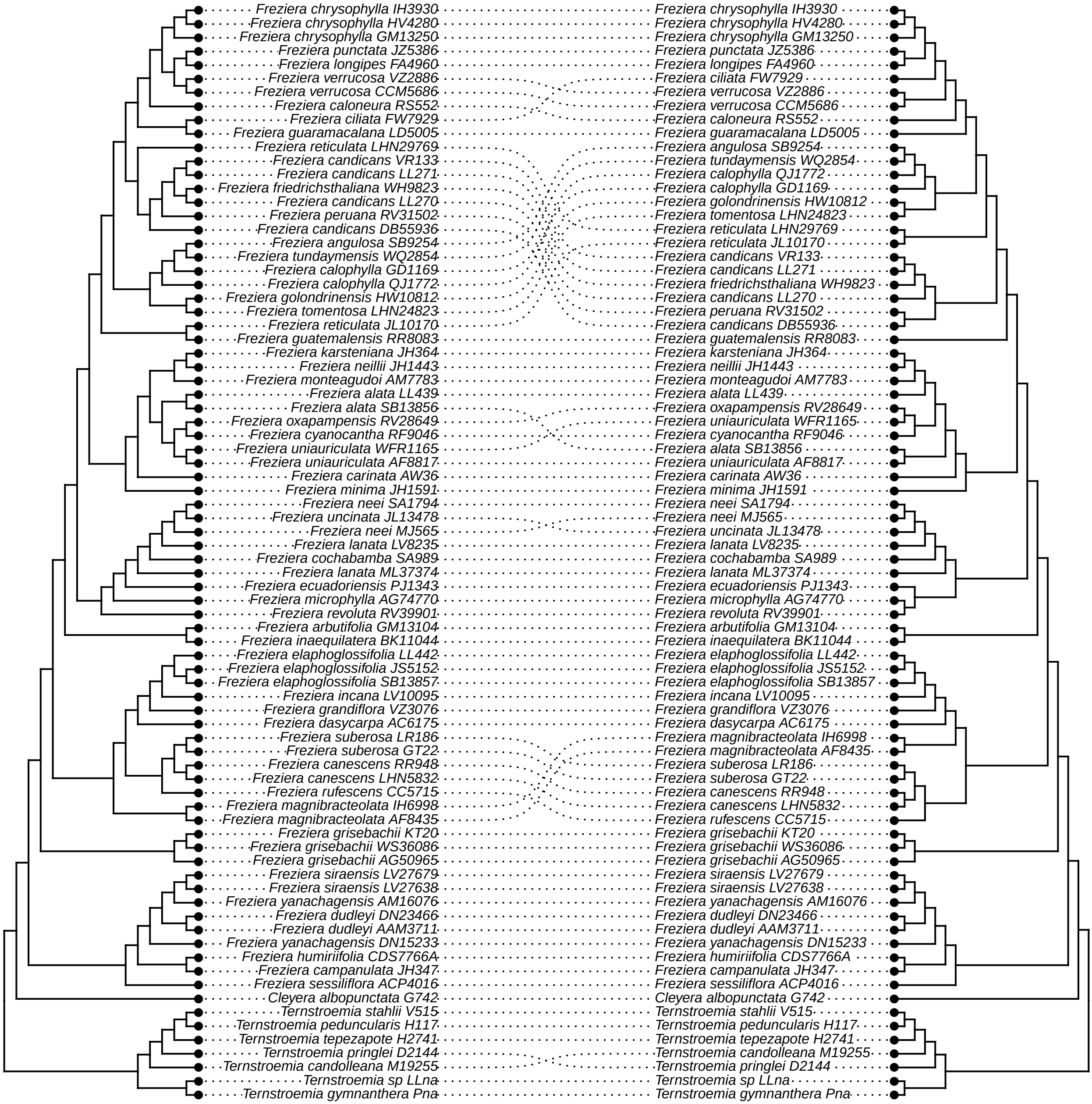

### _t_by_eye_v_tree_length.pdf

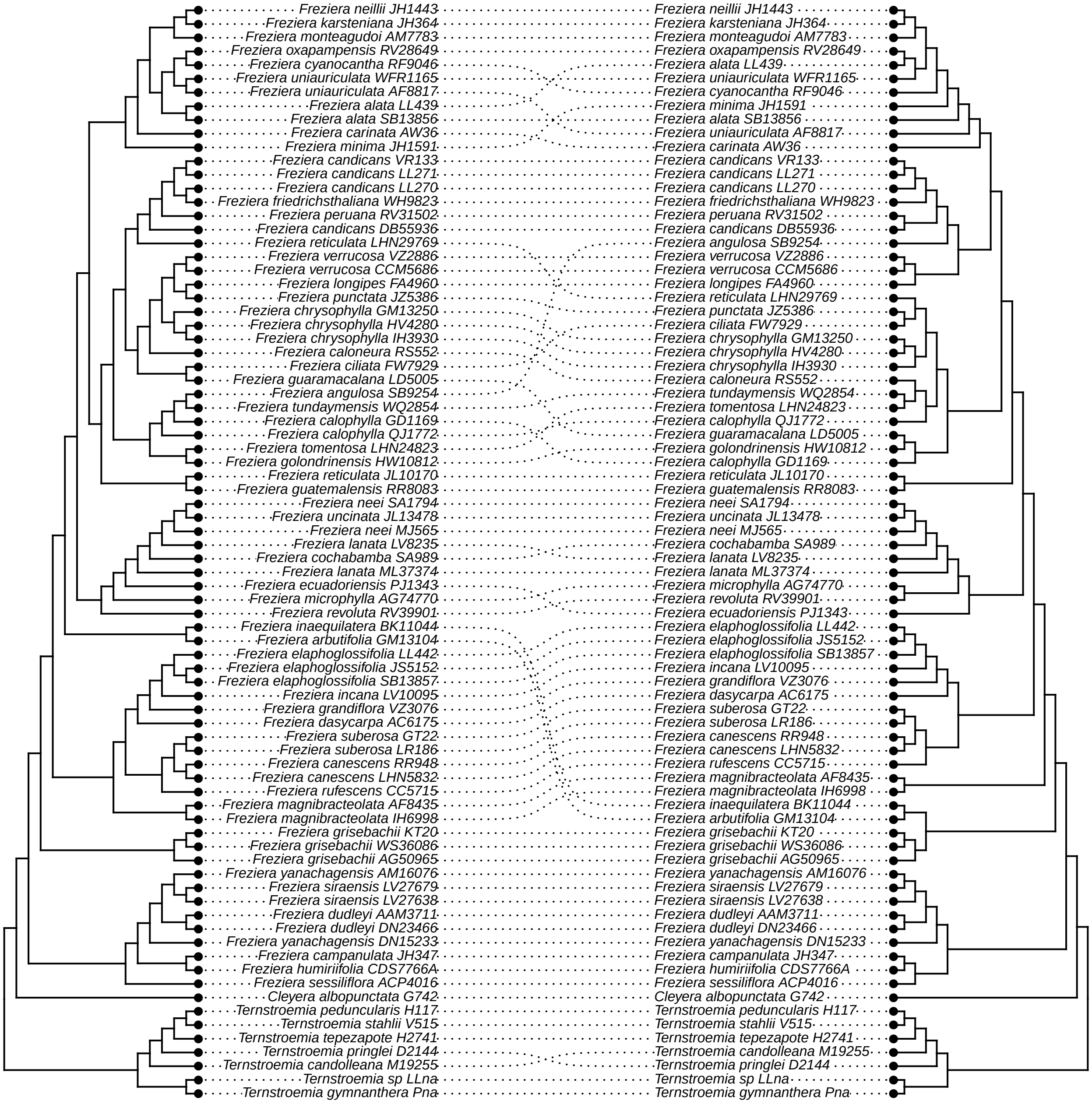

### _u_by_eye_vs_bipartition.pdf

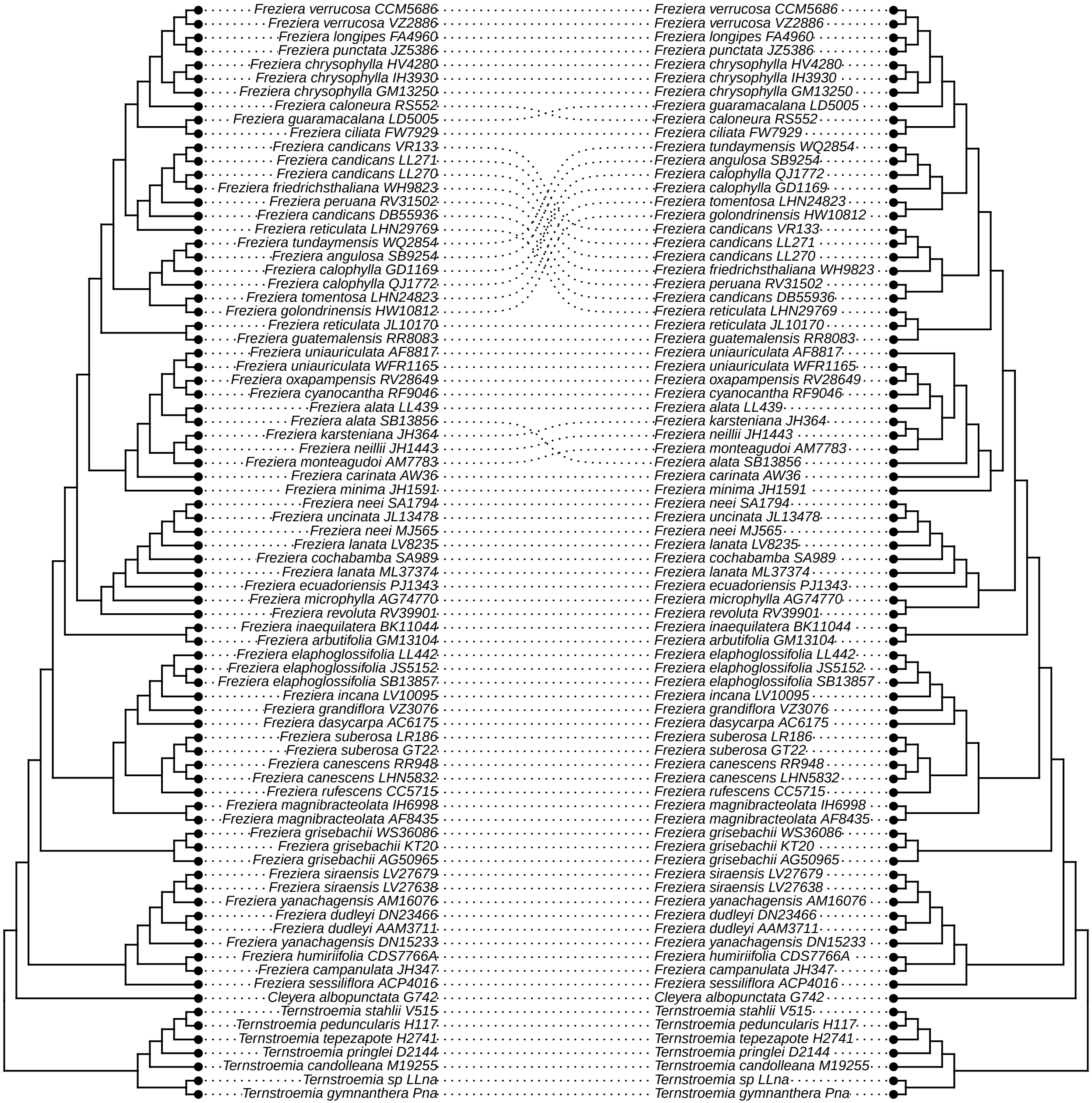

### _v_hp2_no_warnings_vs_unfiltered.pdf

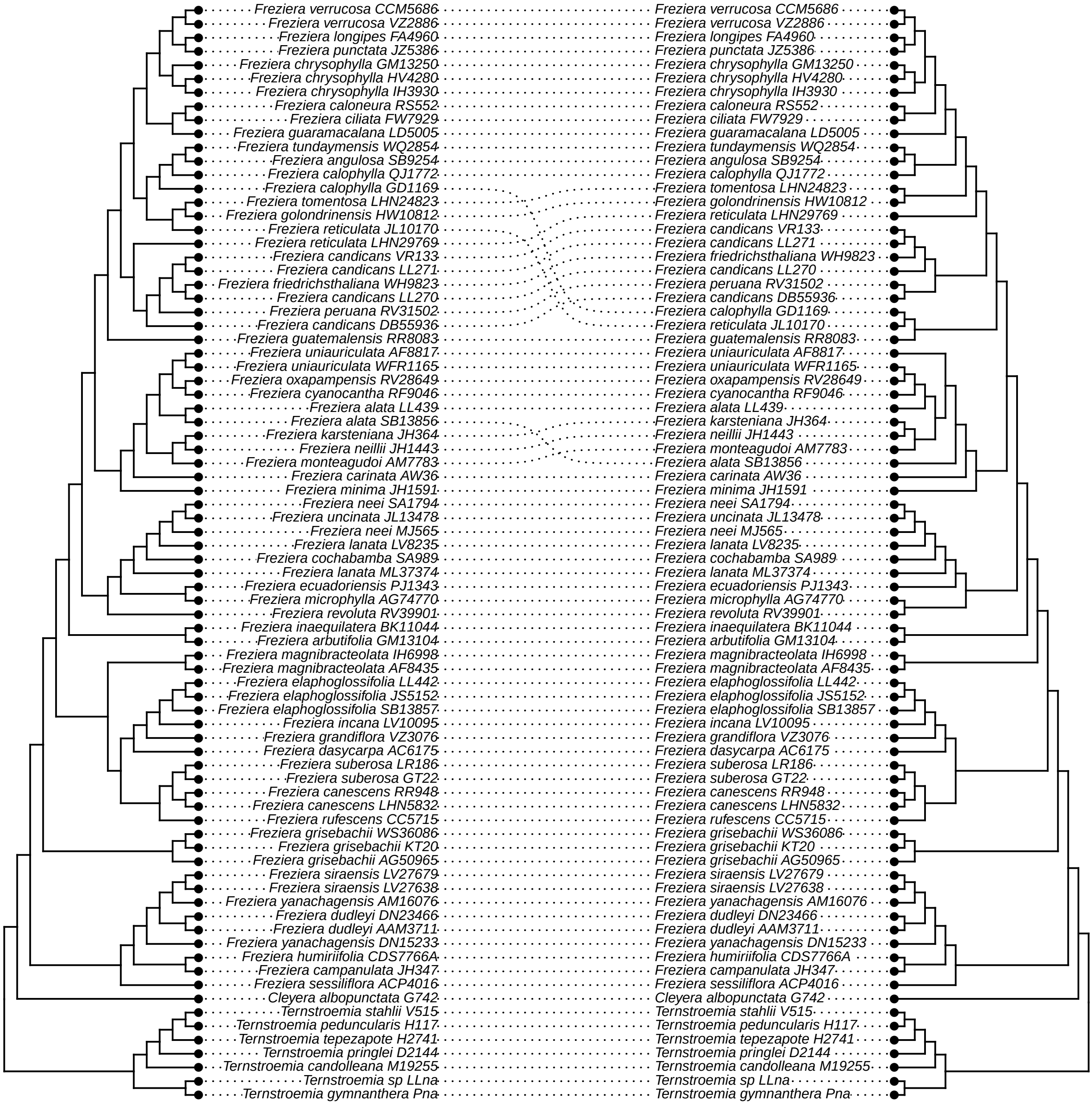

### _w_hp2_no_warnings_vs_hp2_long.pdf

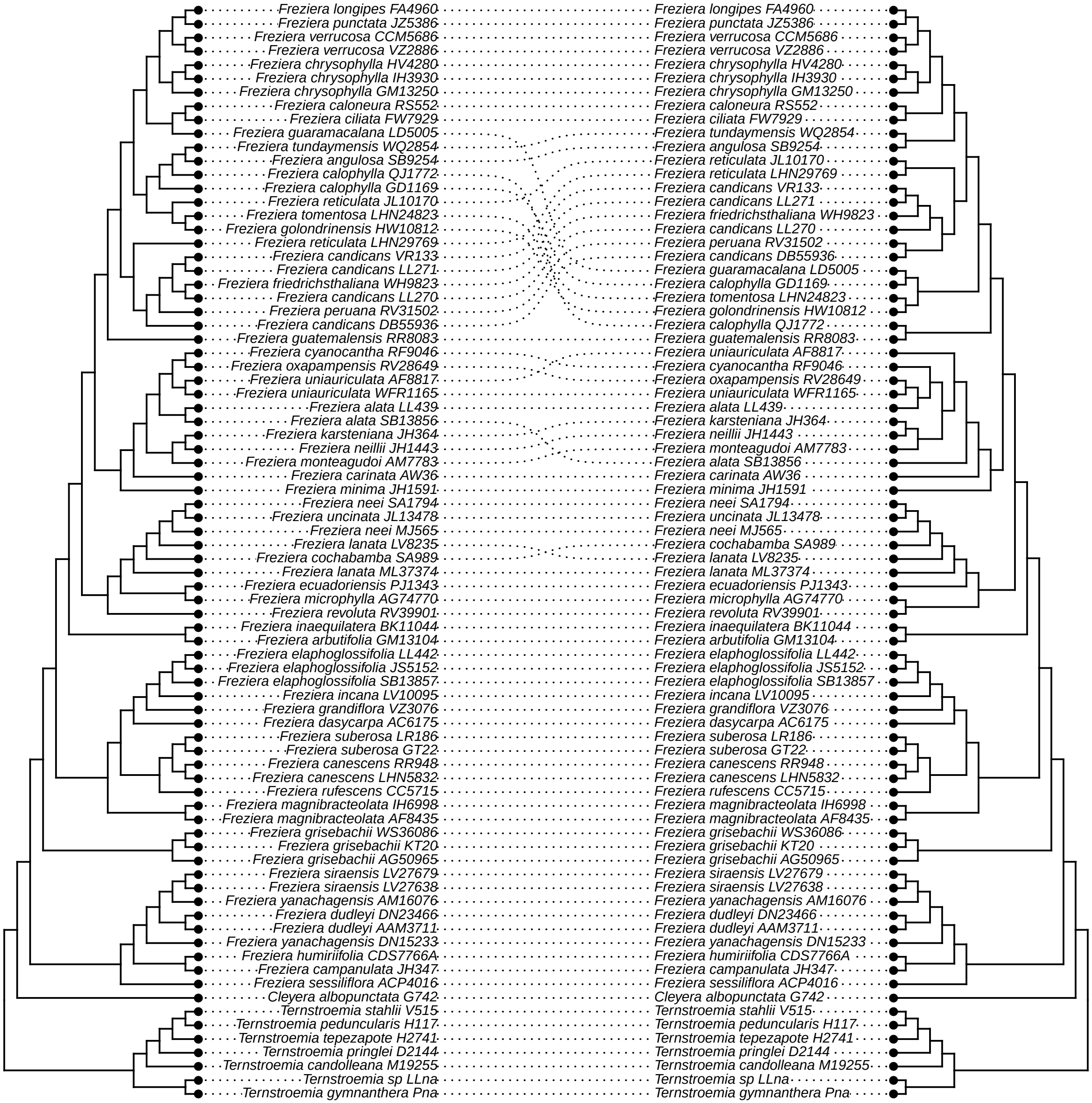

### _x_hp2_no_warnings_vs_HybPhaser.pdf

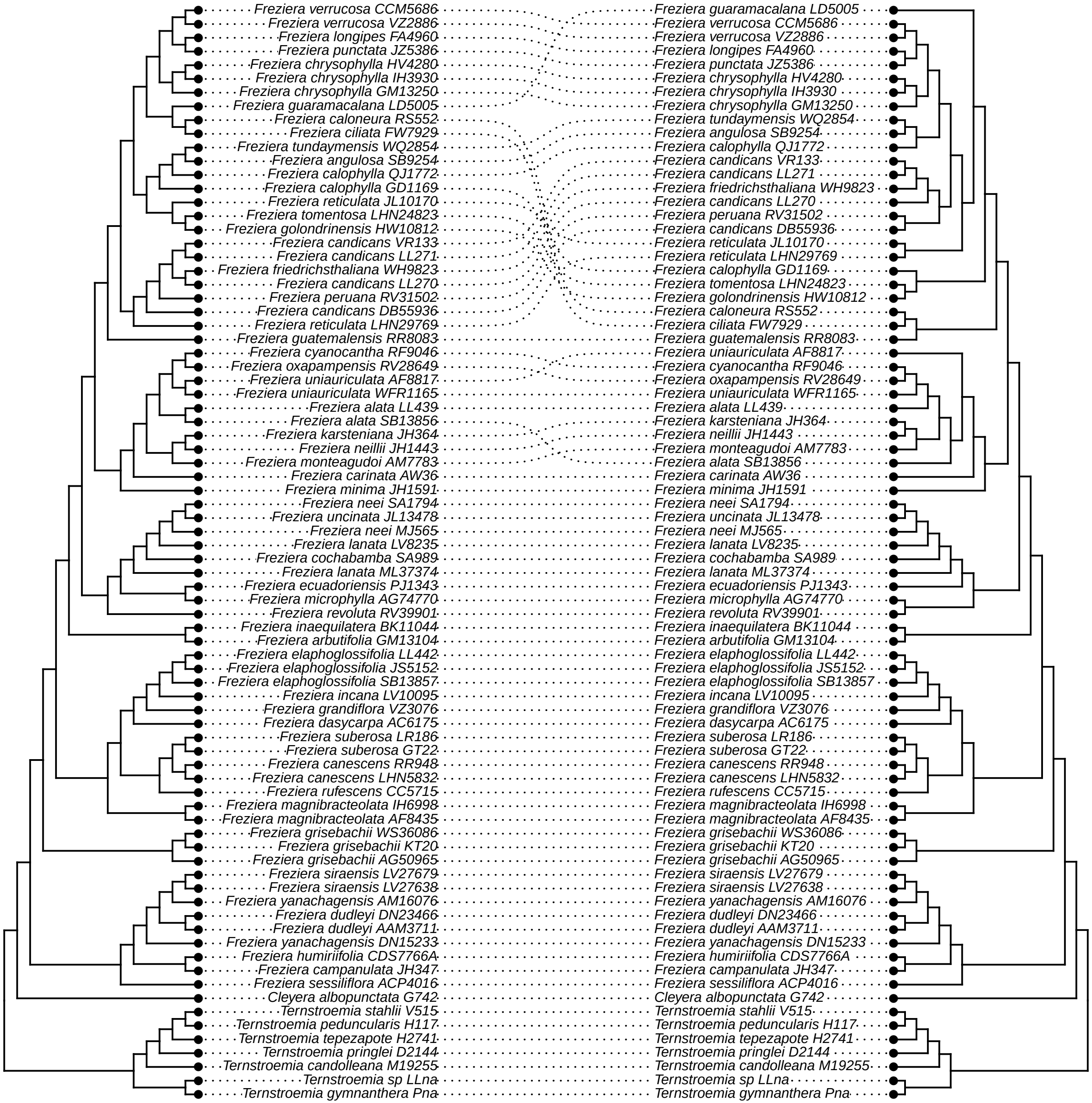

### _y_hp2_no_warnings_v_MO.pdf

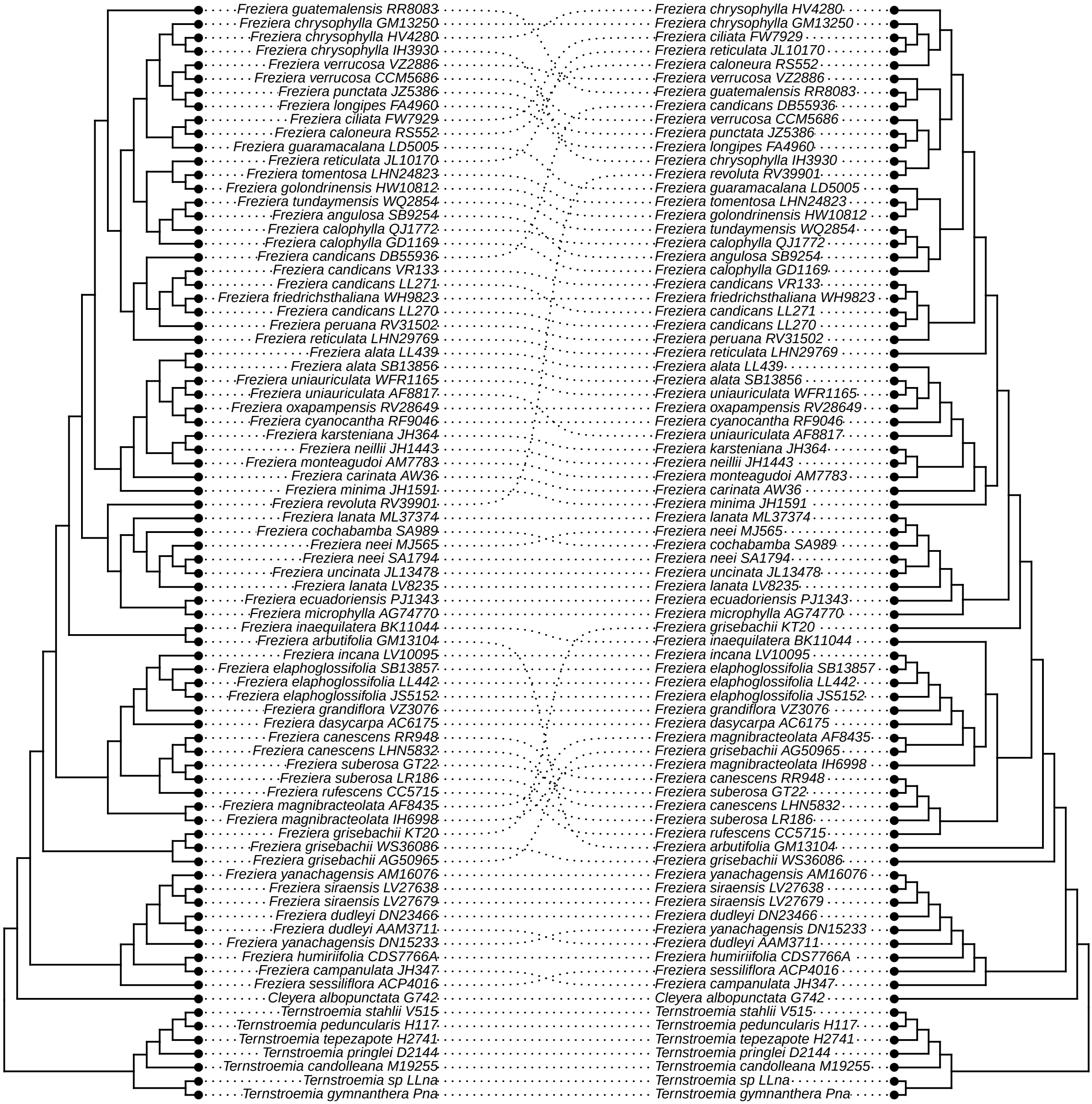

### _z_hp2_no_warnings_v_prop_PI.pdf

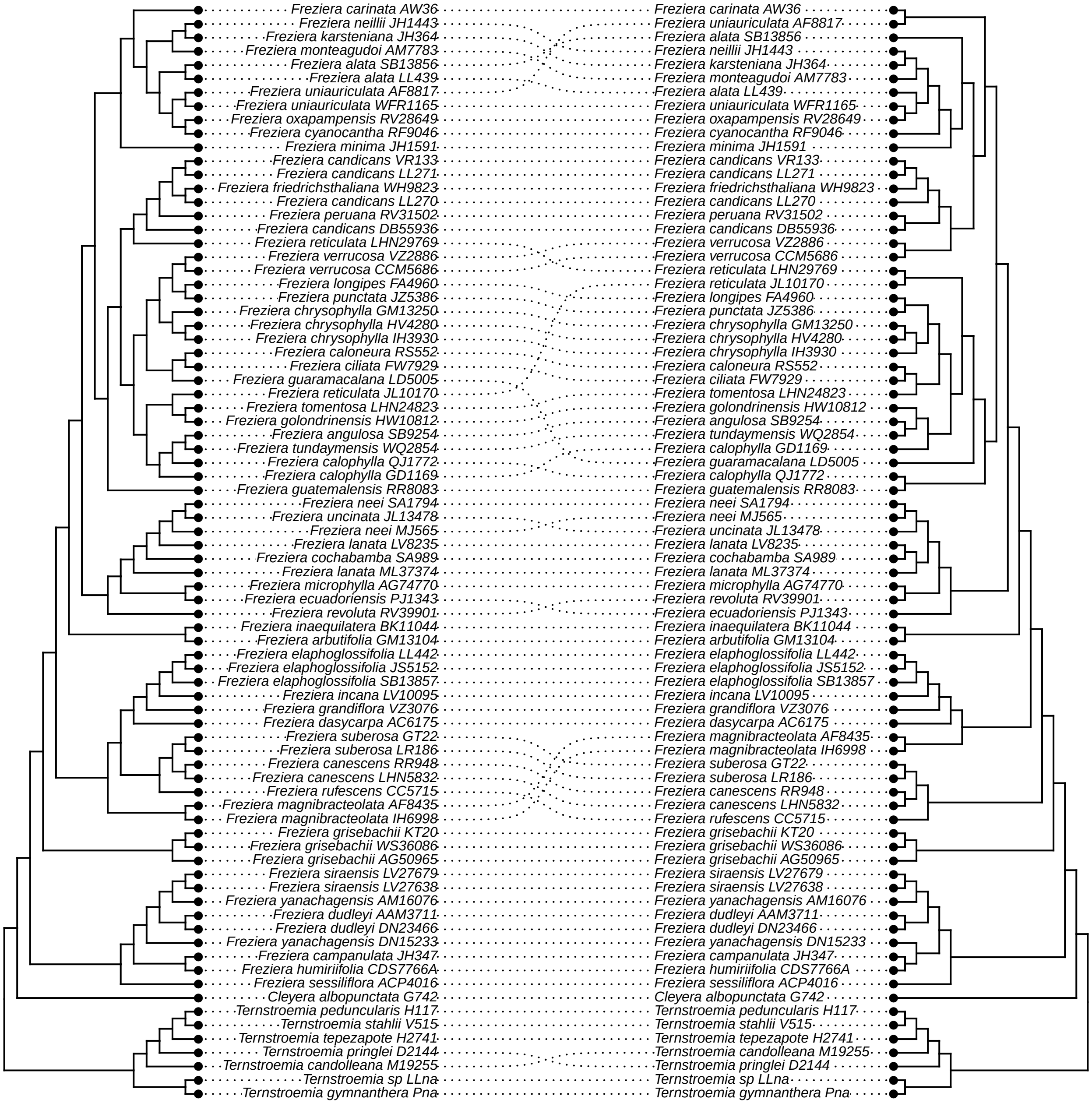

### aa_hp2_no_warnings_v_internal.pdf

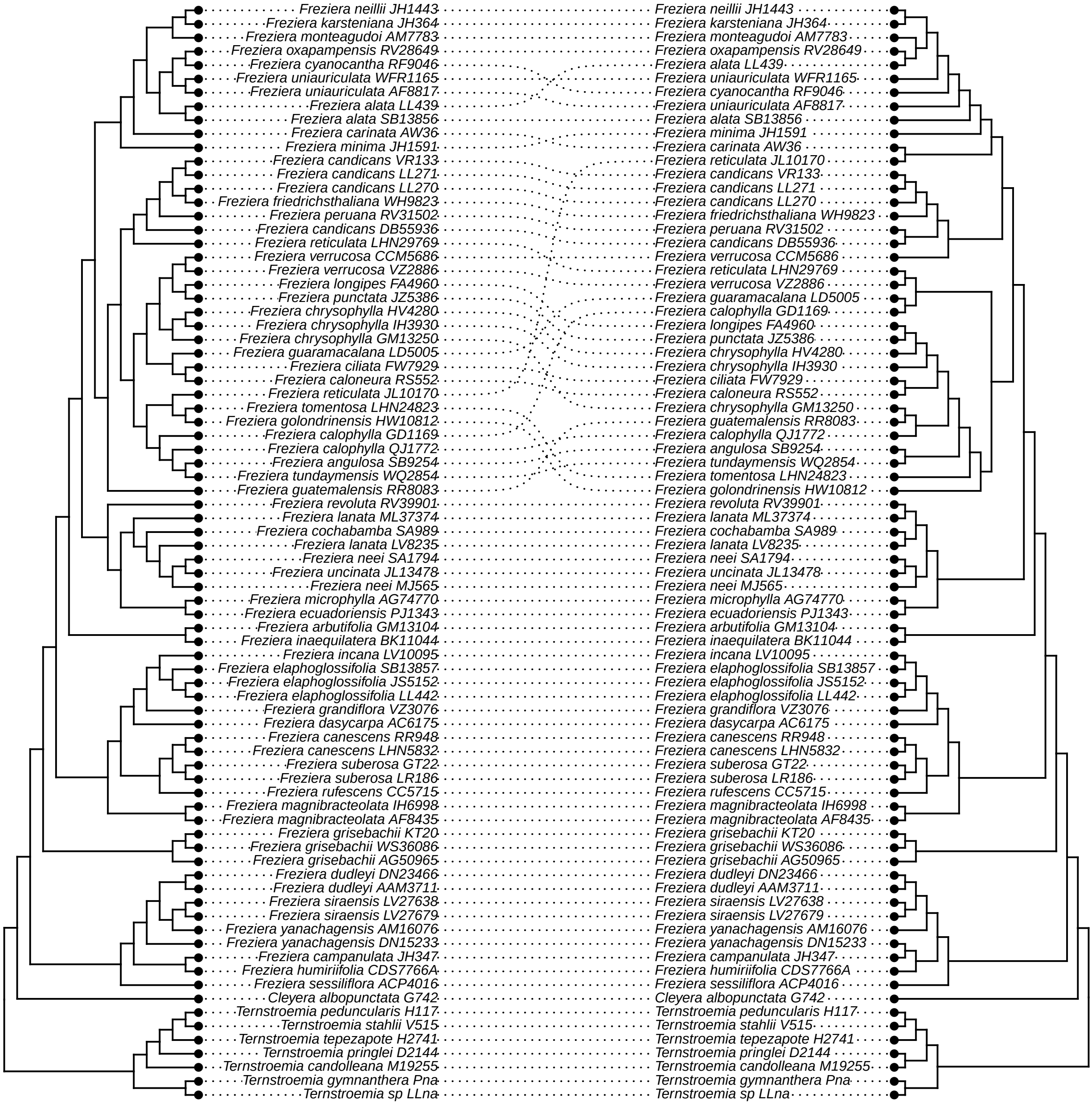

### bb_hp2_no_warnings_v_avg_BS.pdf

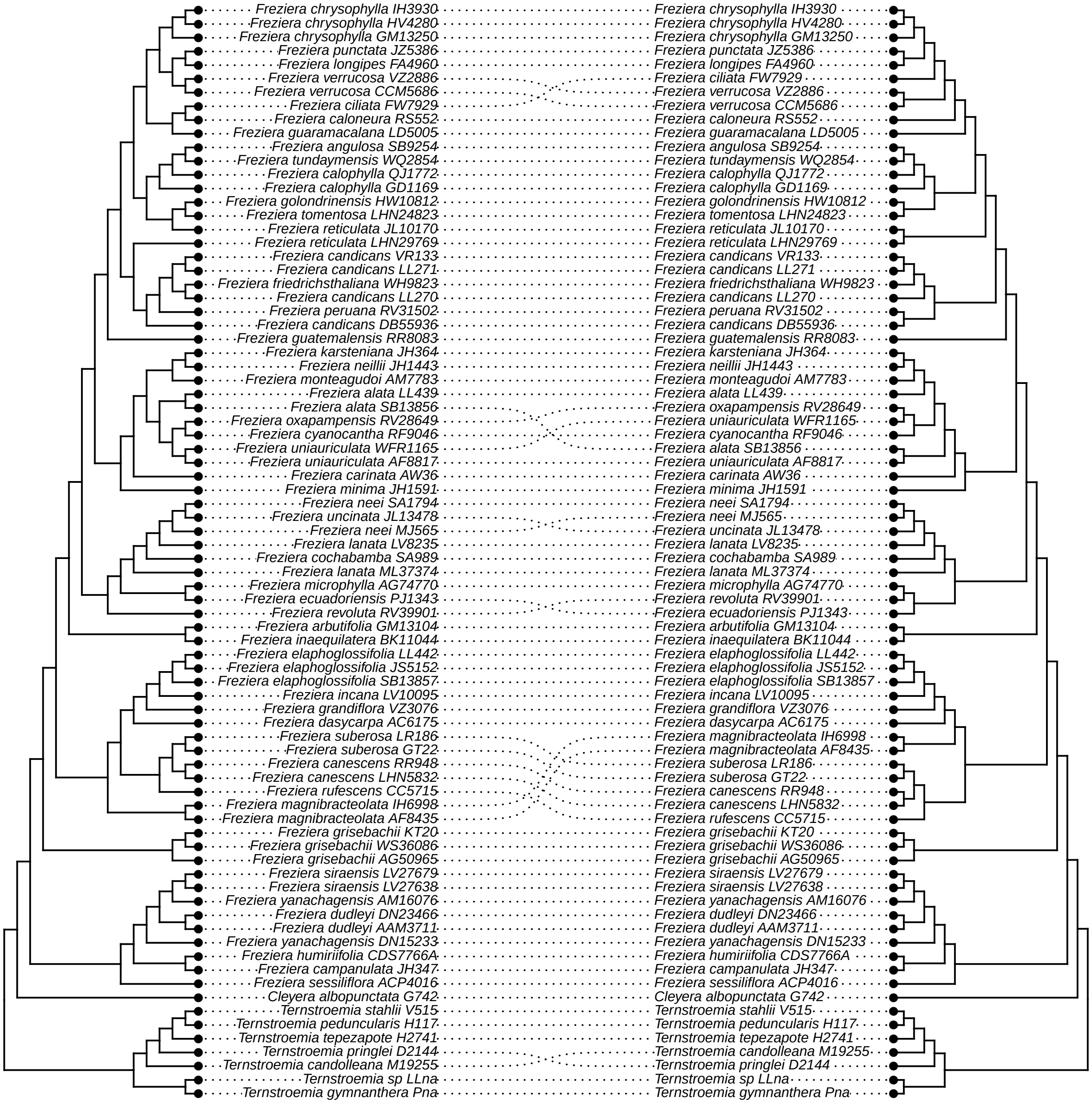

### cc_hp2_no_warnings_v_tree_length.pdf

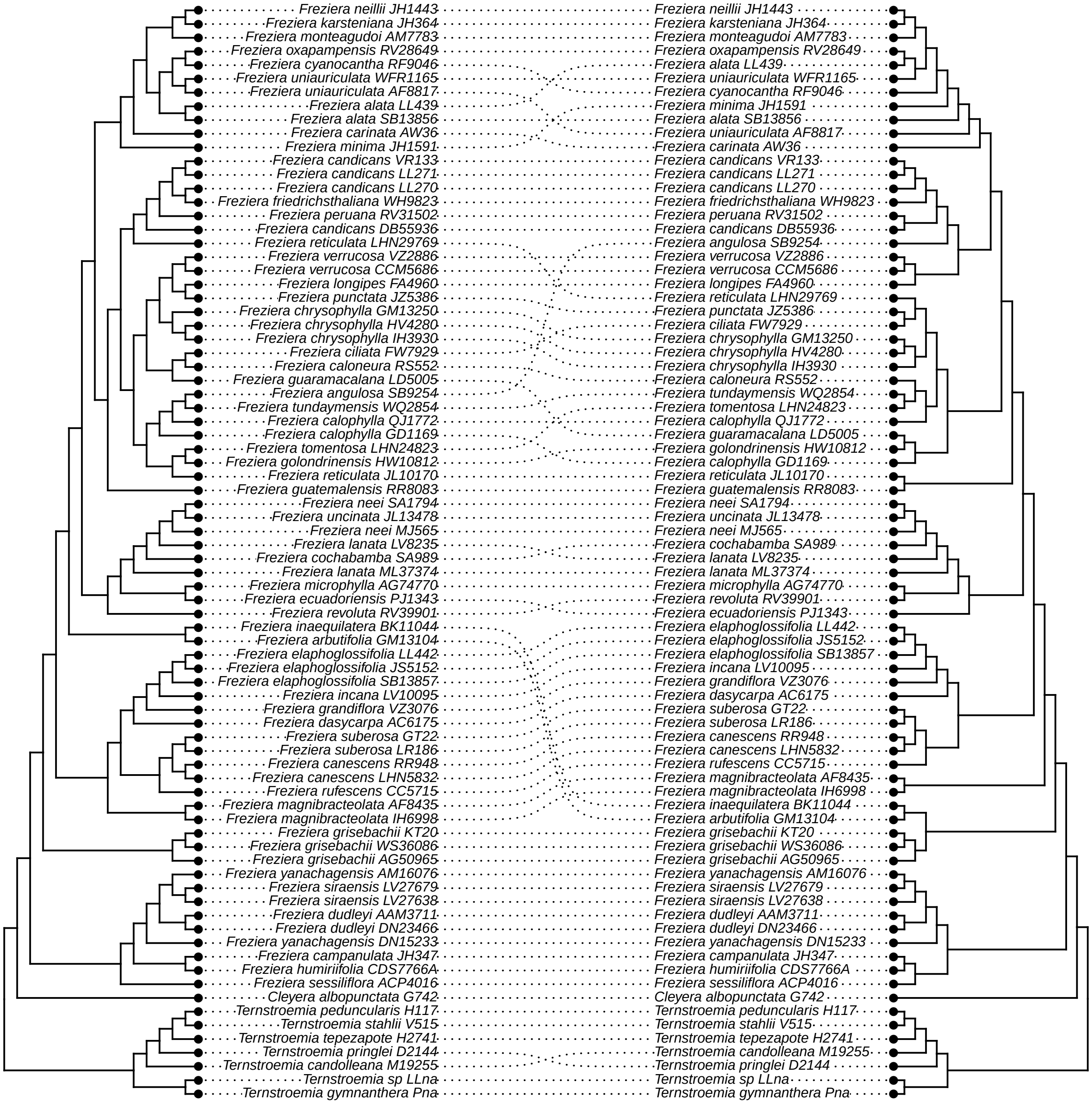

### dd_hp2_no_warnings_vs_bipartition.pdf

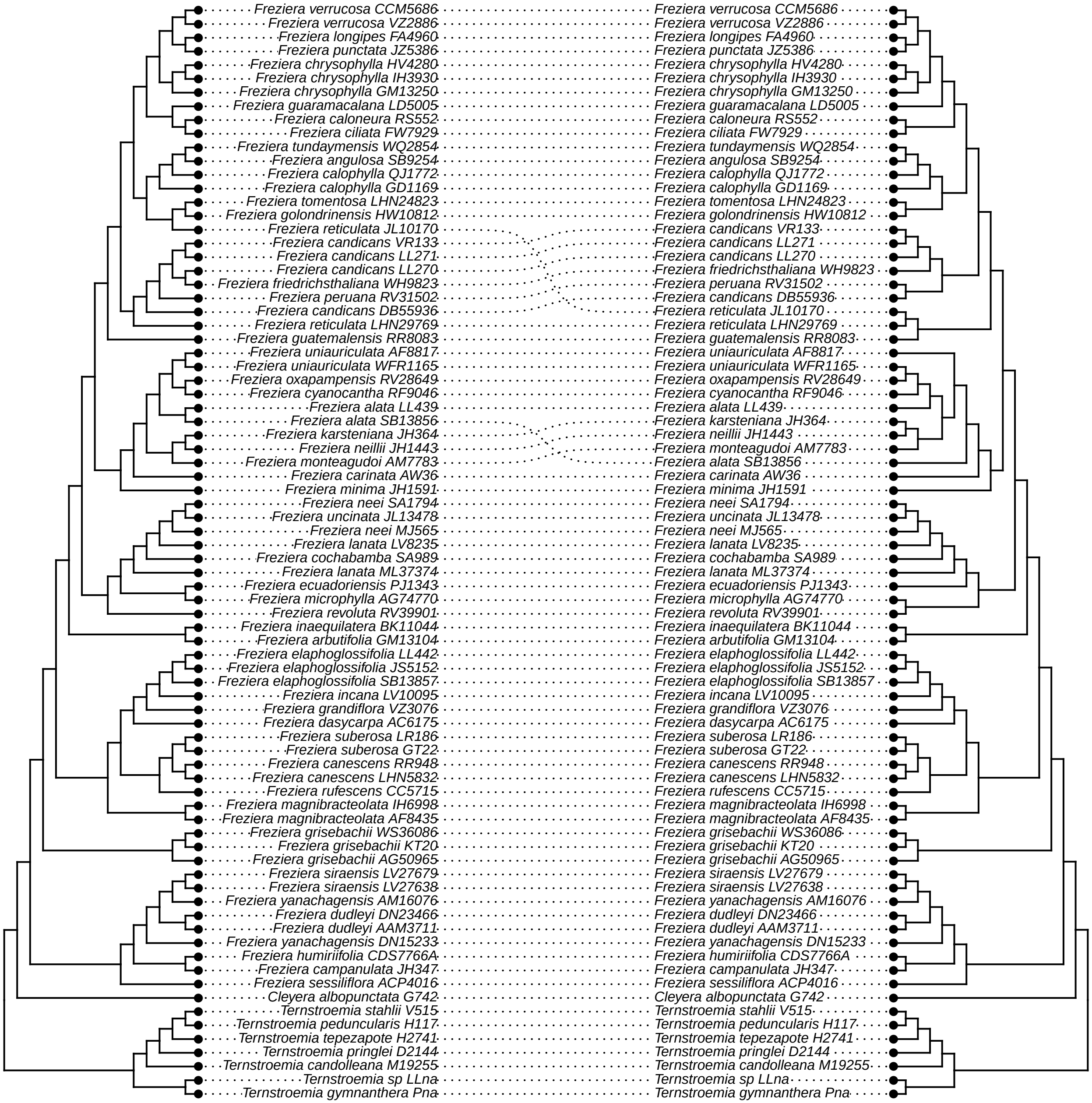

### ee_HybPhaser_vs_unfiltered.pdf

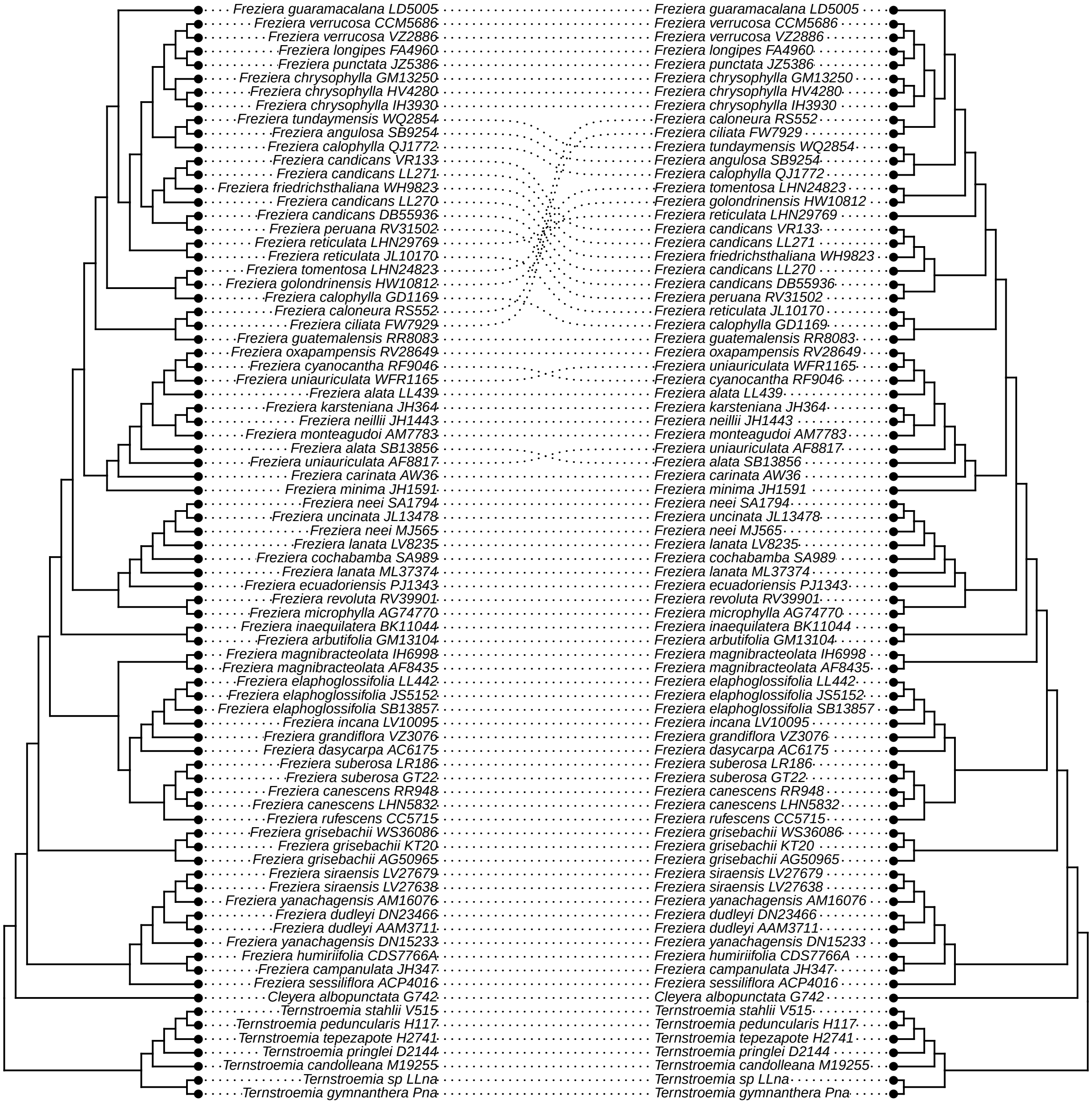

### ff_HybPhaser_v_hp2_long.pdf

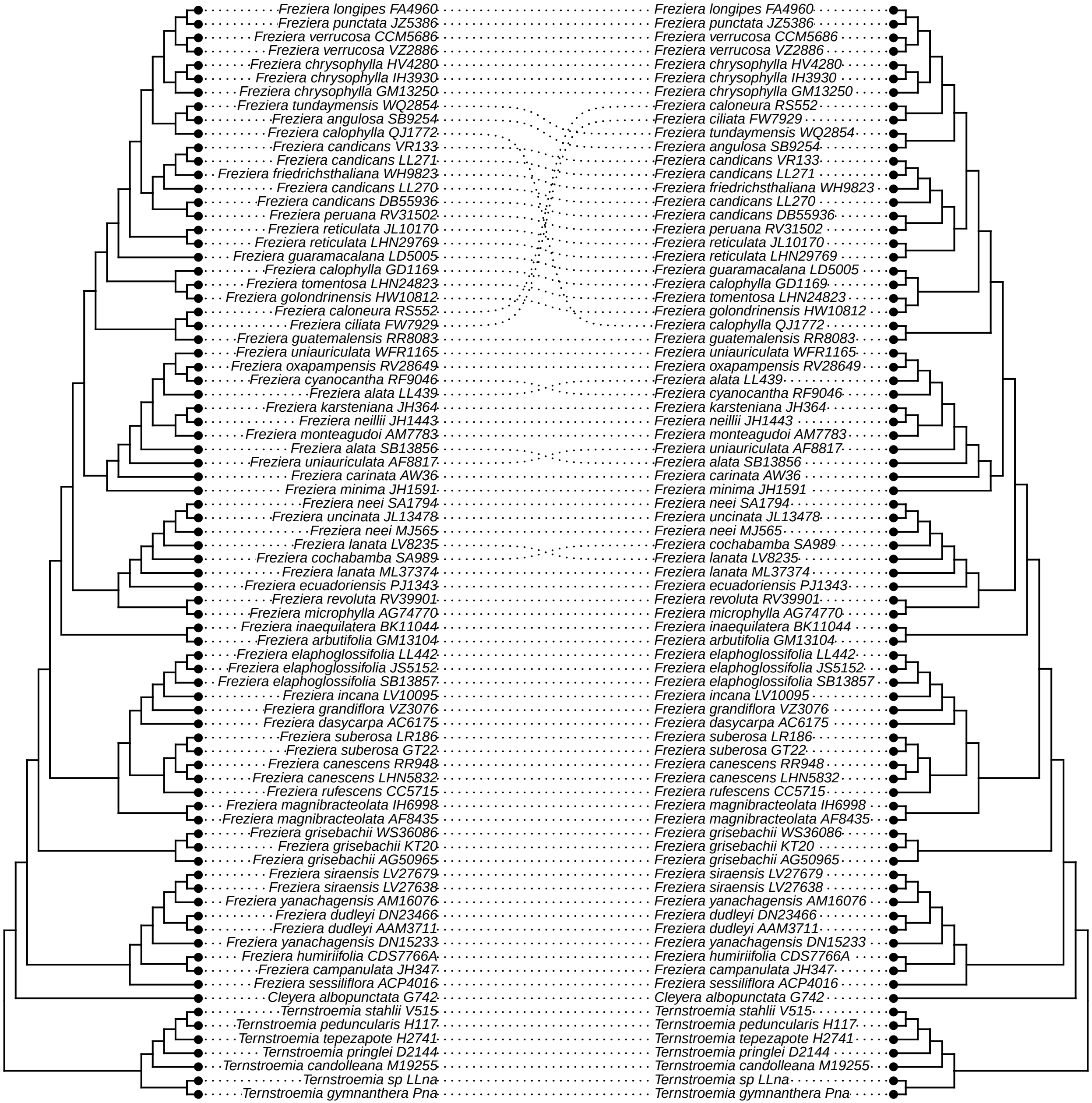

### Figure S1

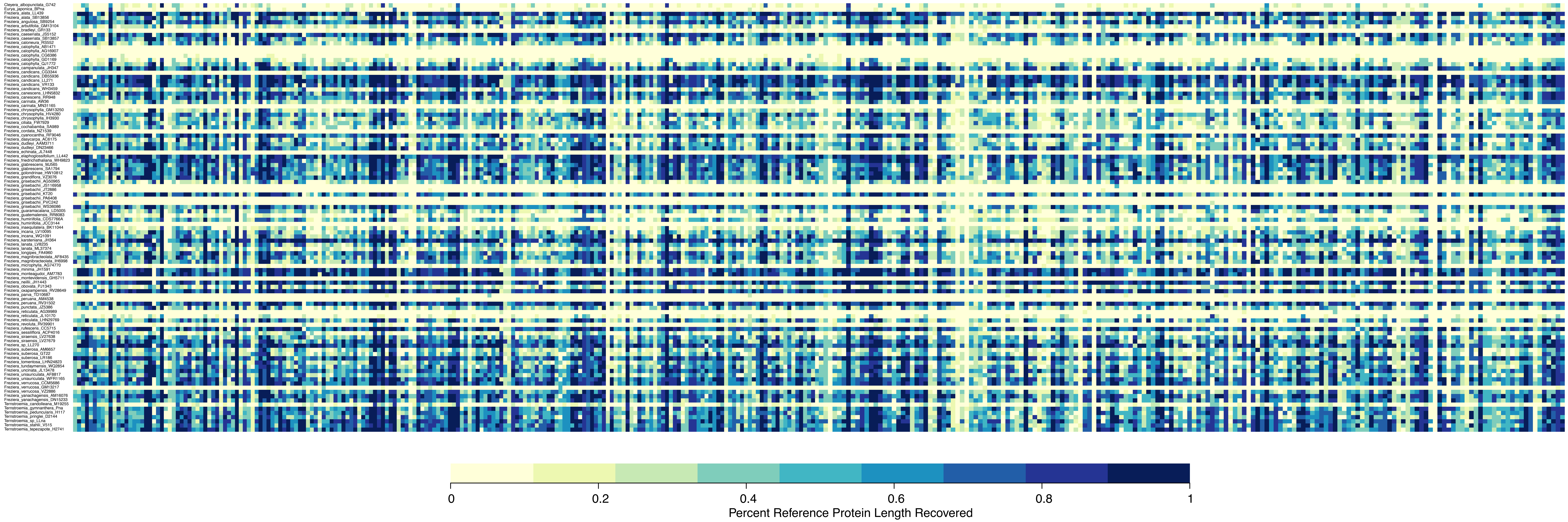

### Figure S3

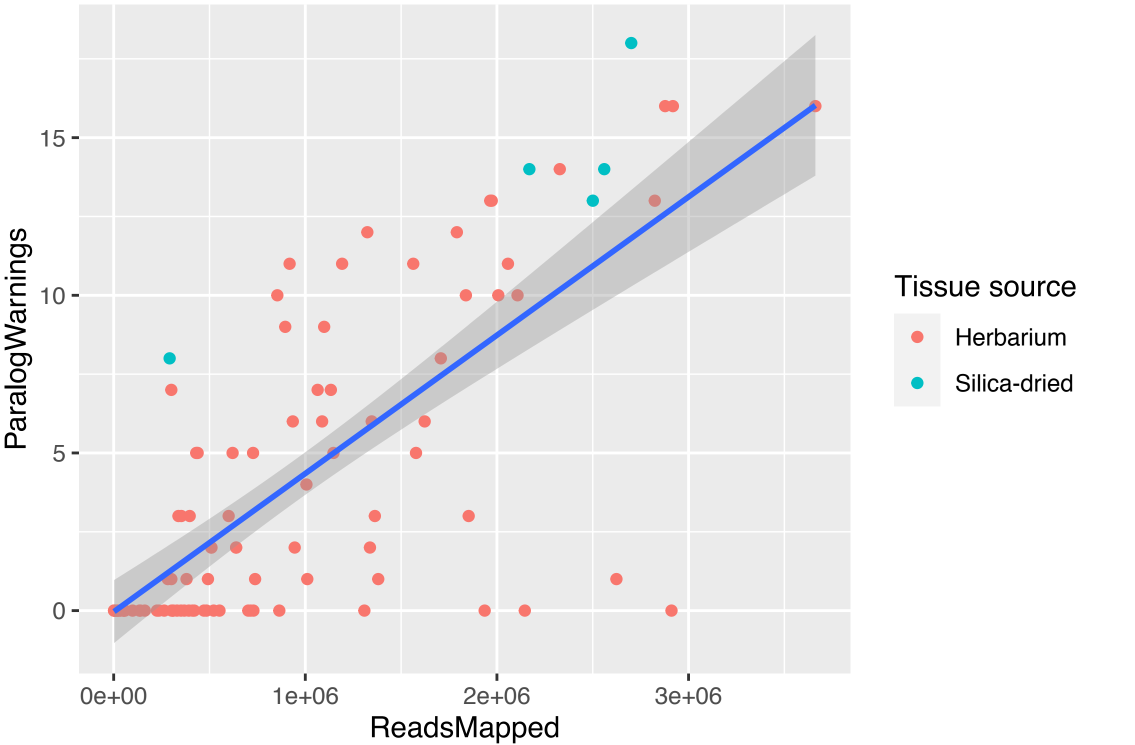

### Figure S5

#### Bipartition support (166)

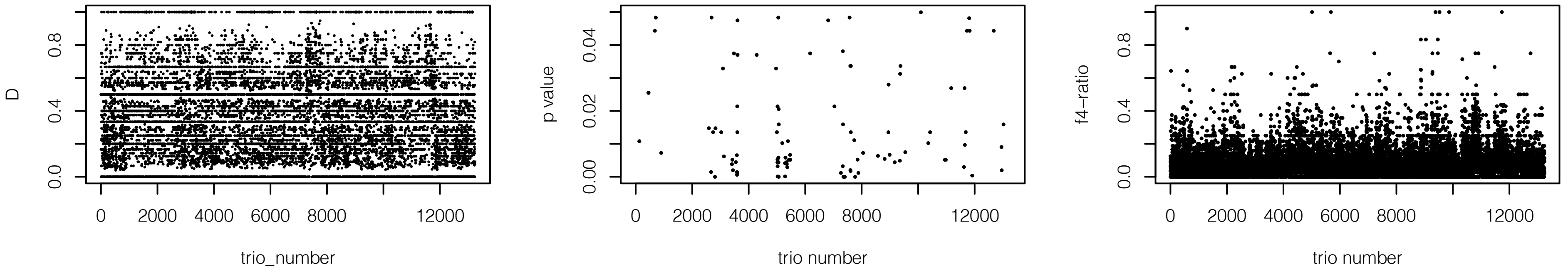


#### By eye (182)

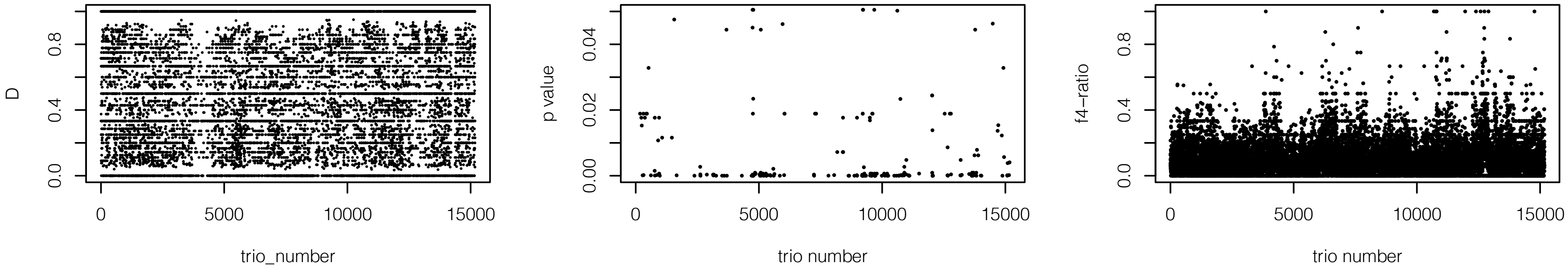


#### HybPiper2 no warnings (187)

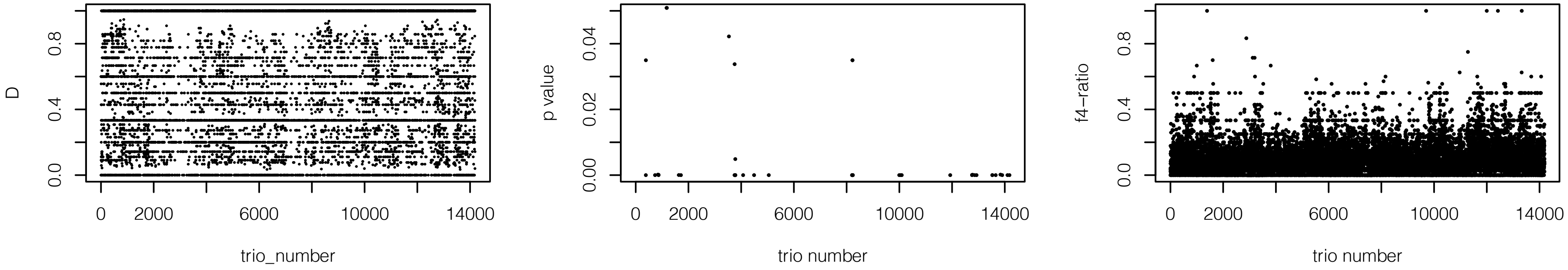

### Figure S6

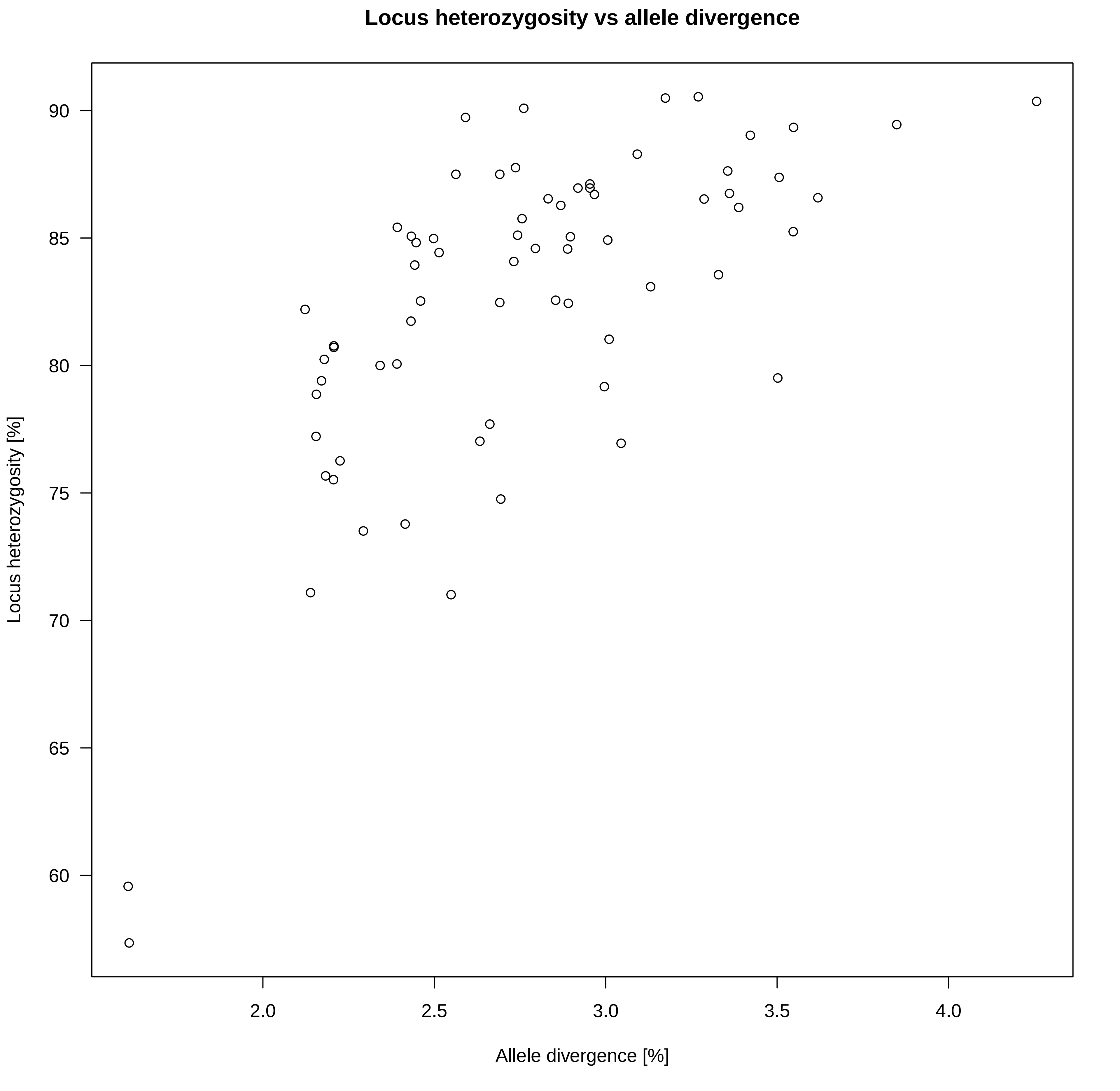

### gg_HybPhaser_v_MO.pdf

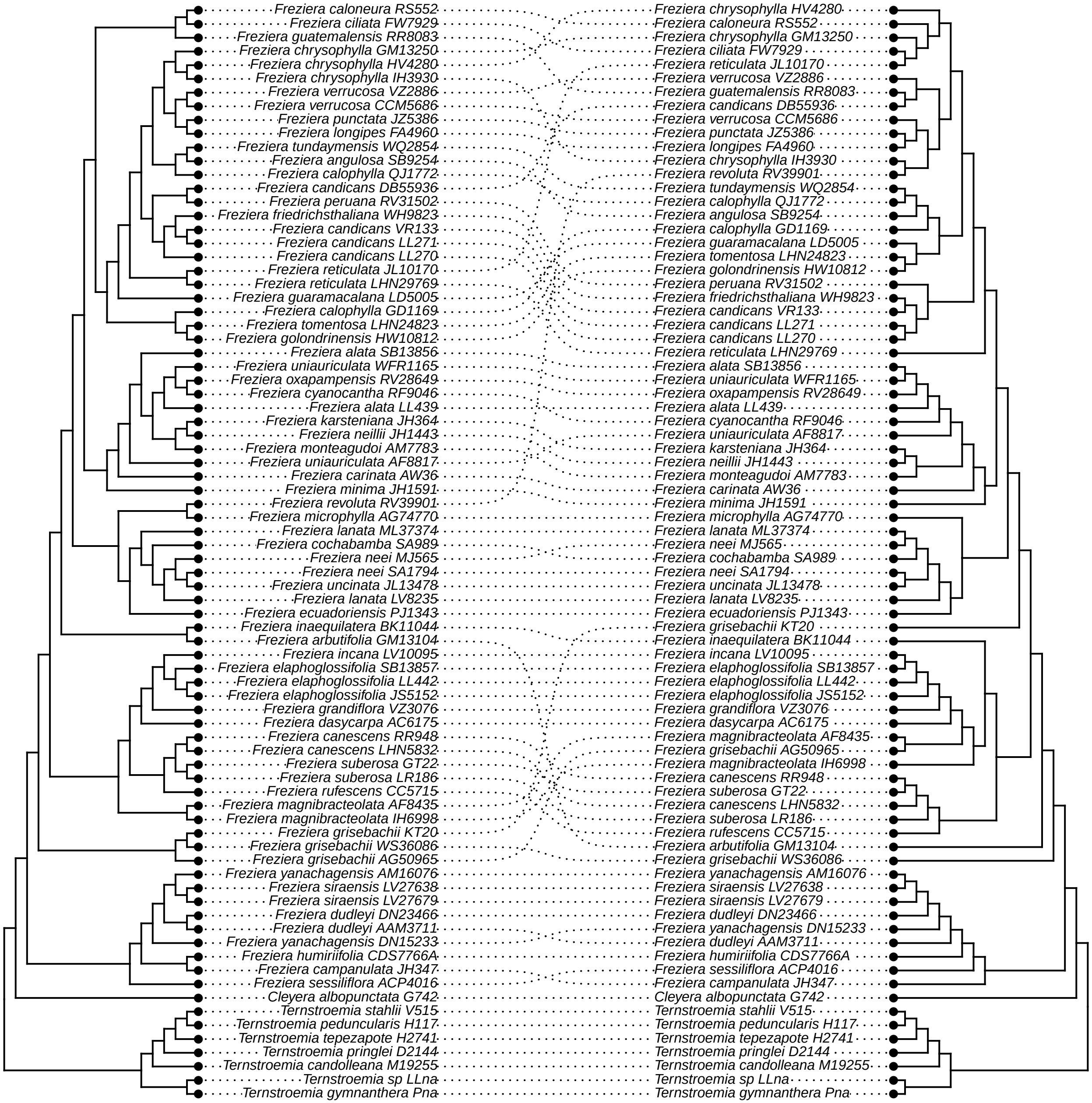

### hh_HybPhaser_v_prop_PI.pdf

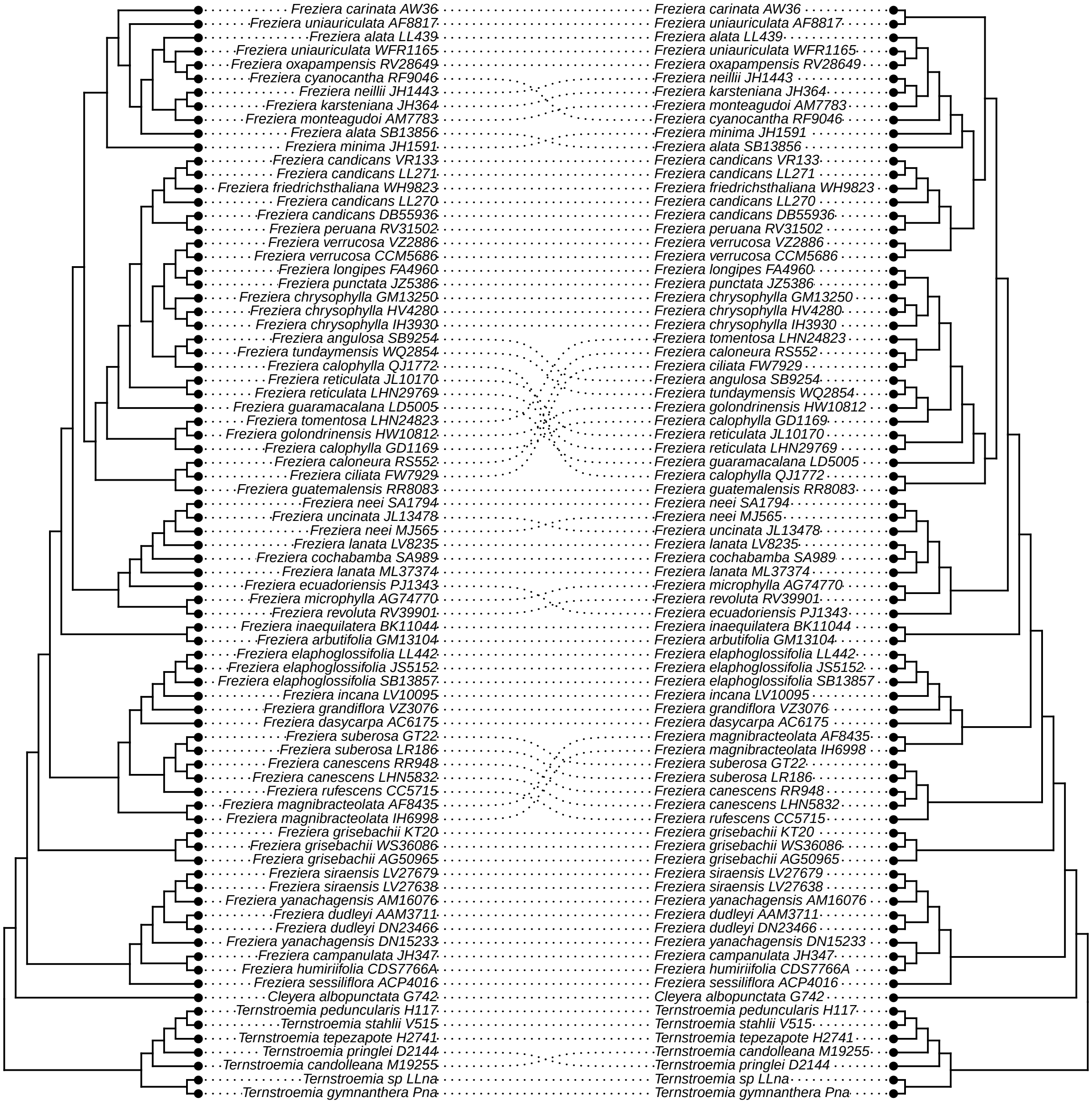

### ii_HybPhaser_v_internal.pdf

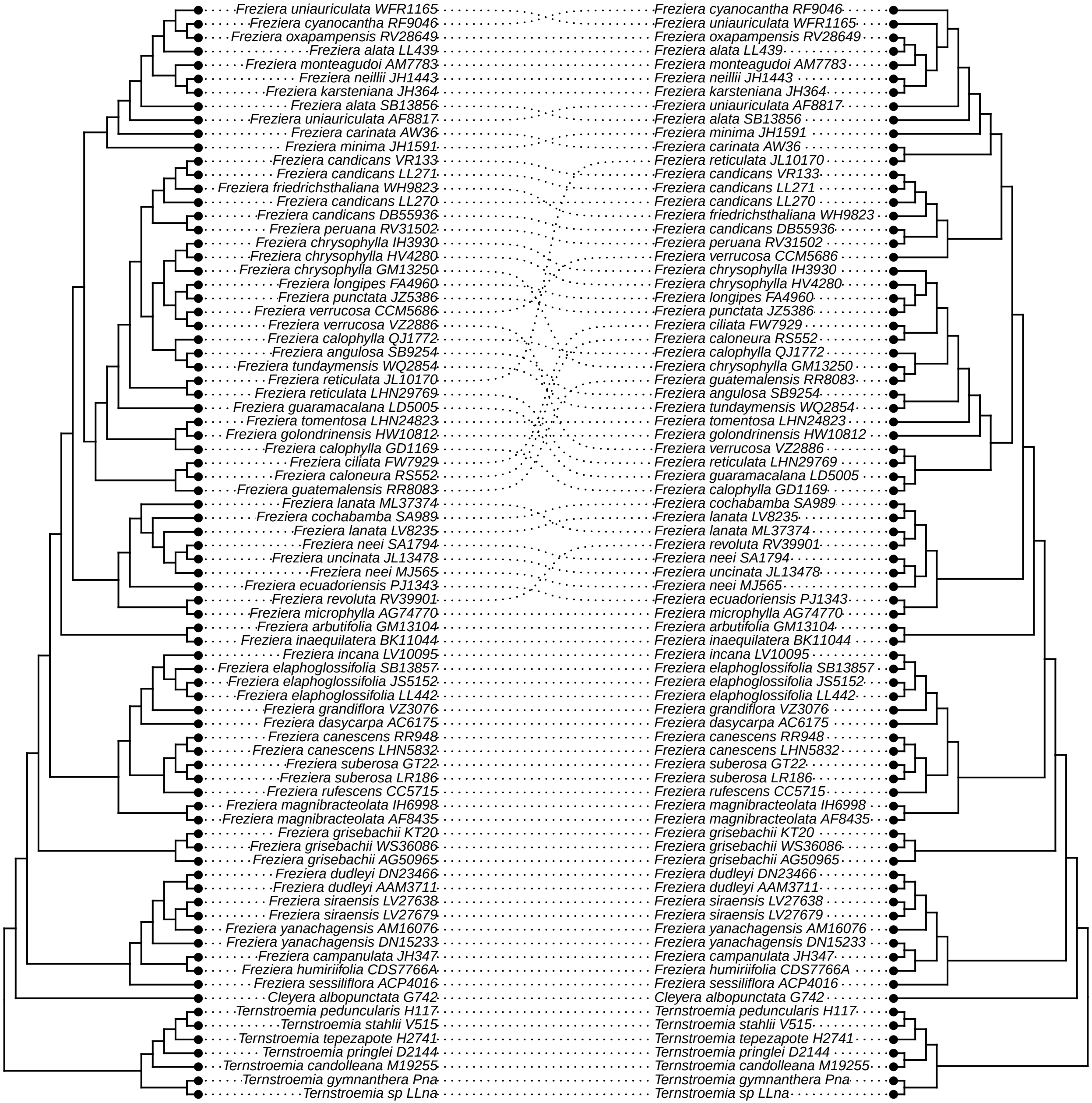

### jj_HybPhaser_v_avg_BS.pdf

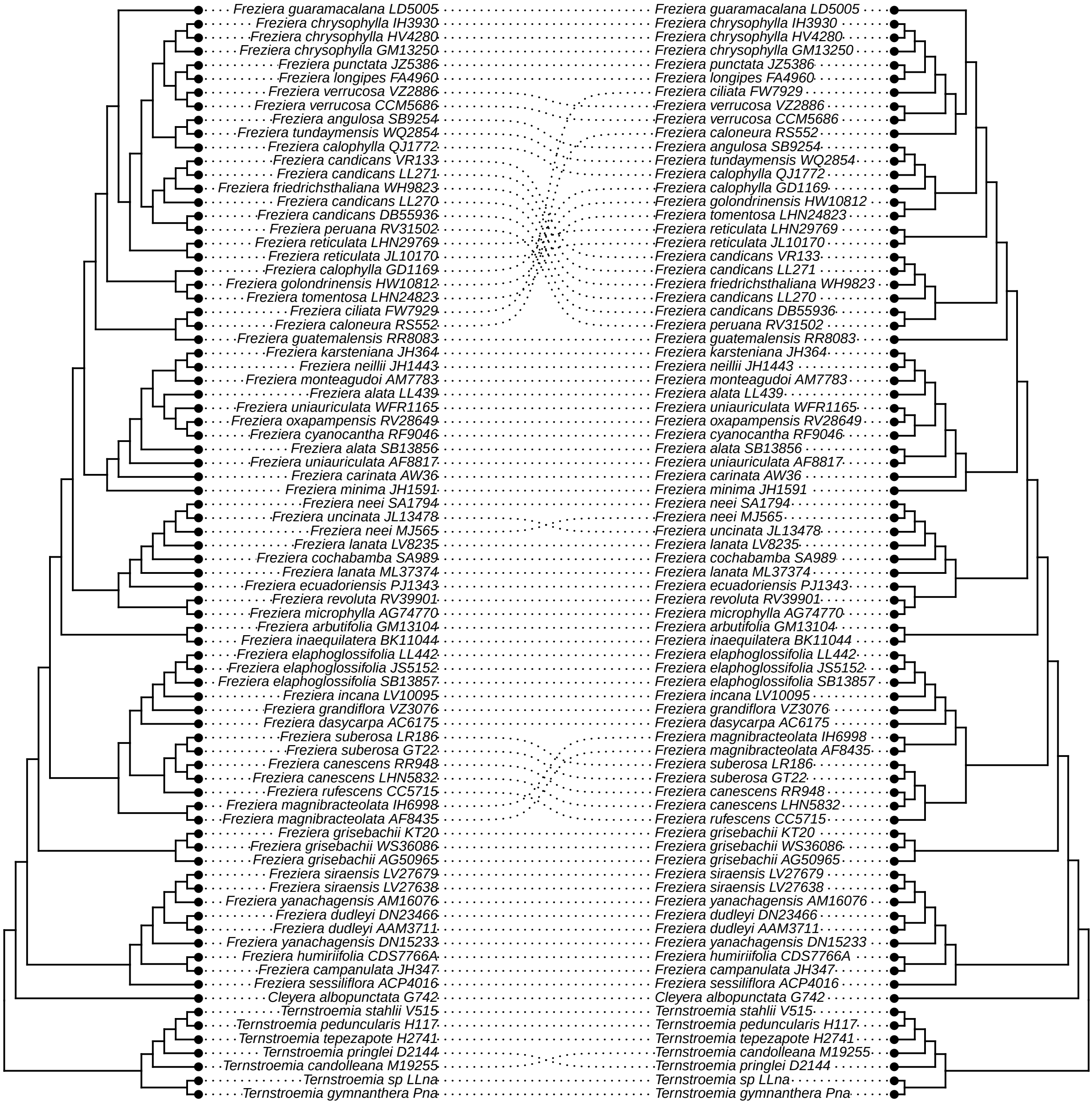

### kk_HybPhaser_v_tree_length.pdf

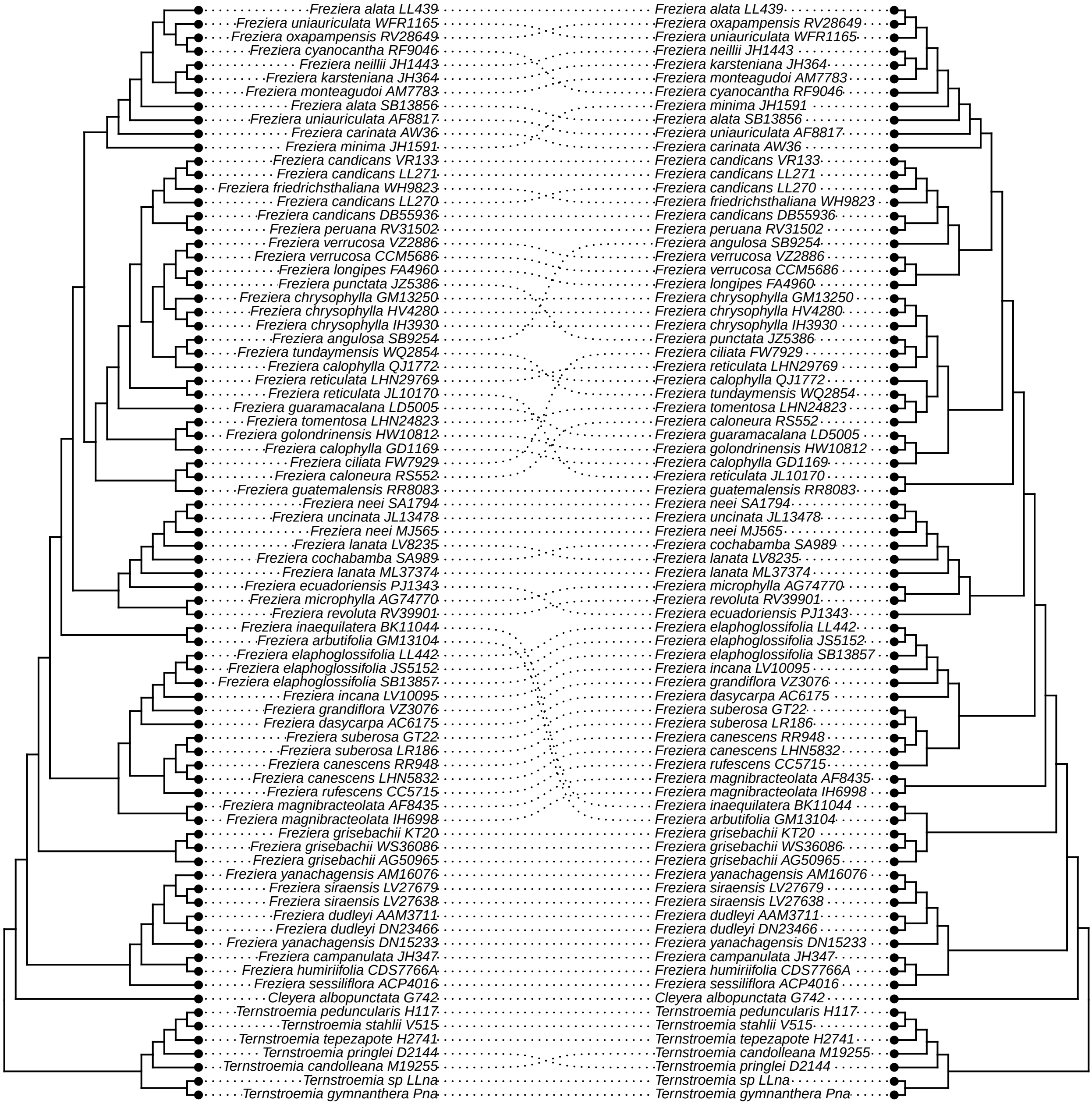

### ll_HybPhaser_vs_bipartition.pdf

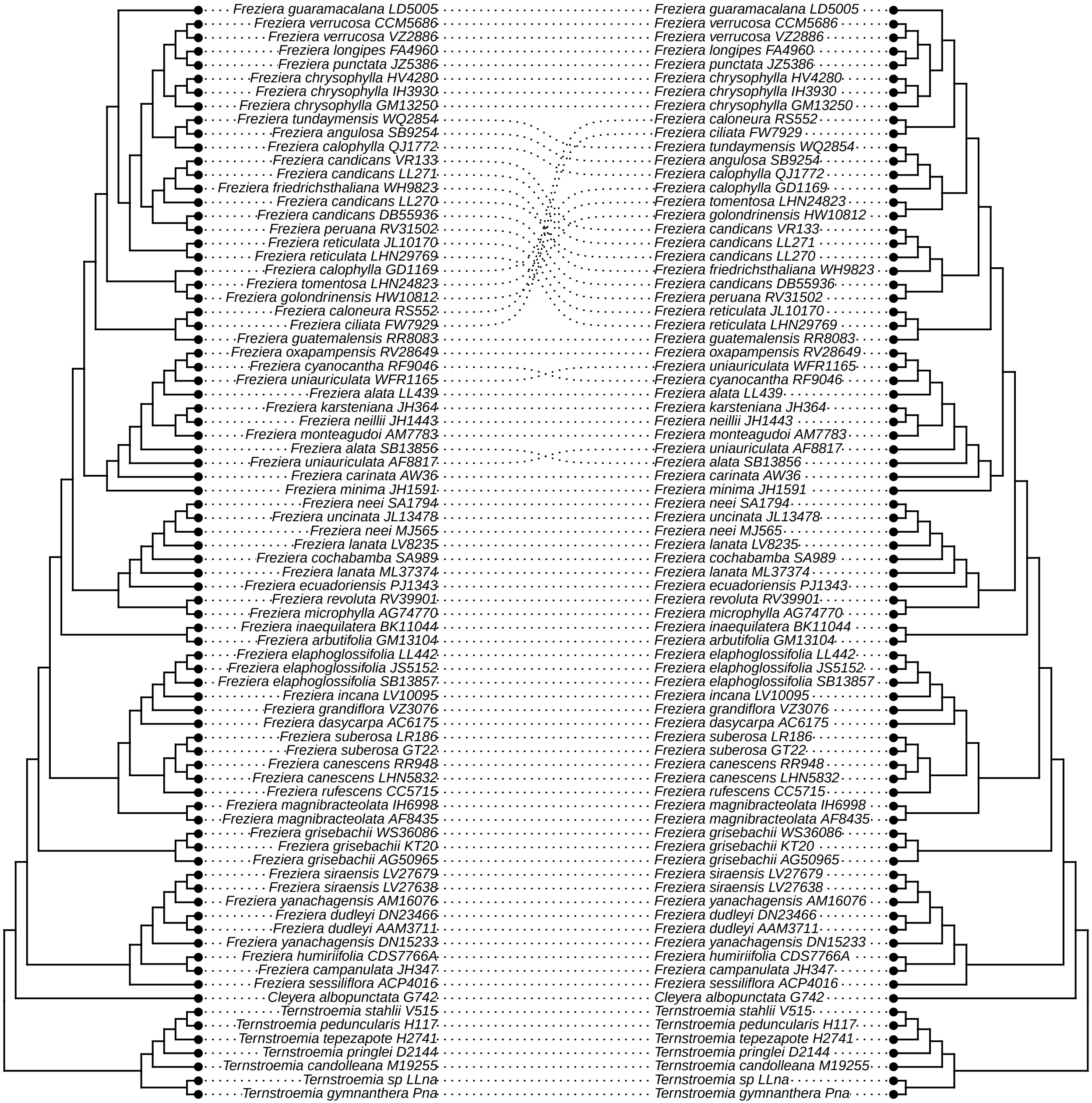

### mm_bipartition_vs_unfiltered.pdf

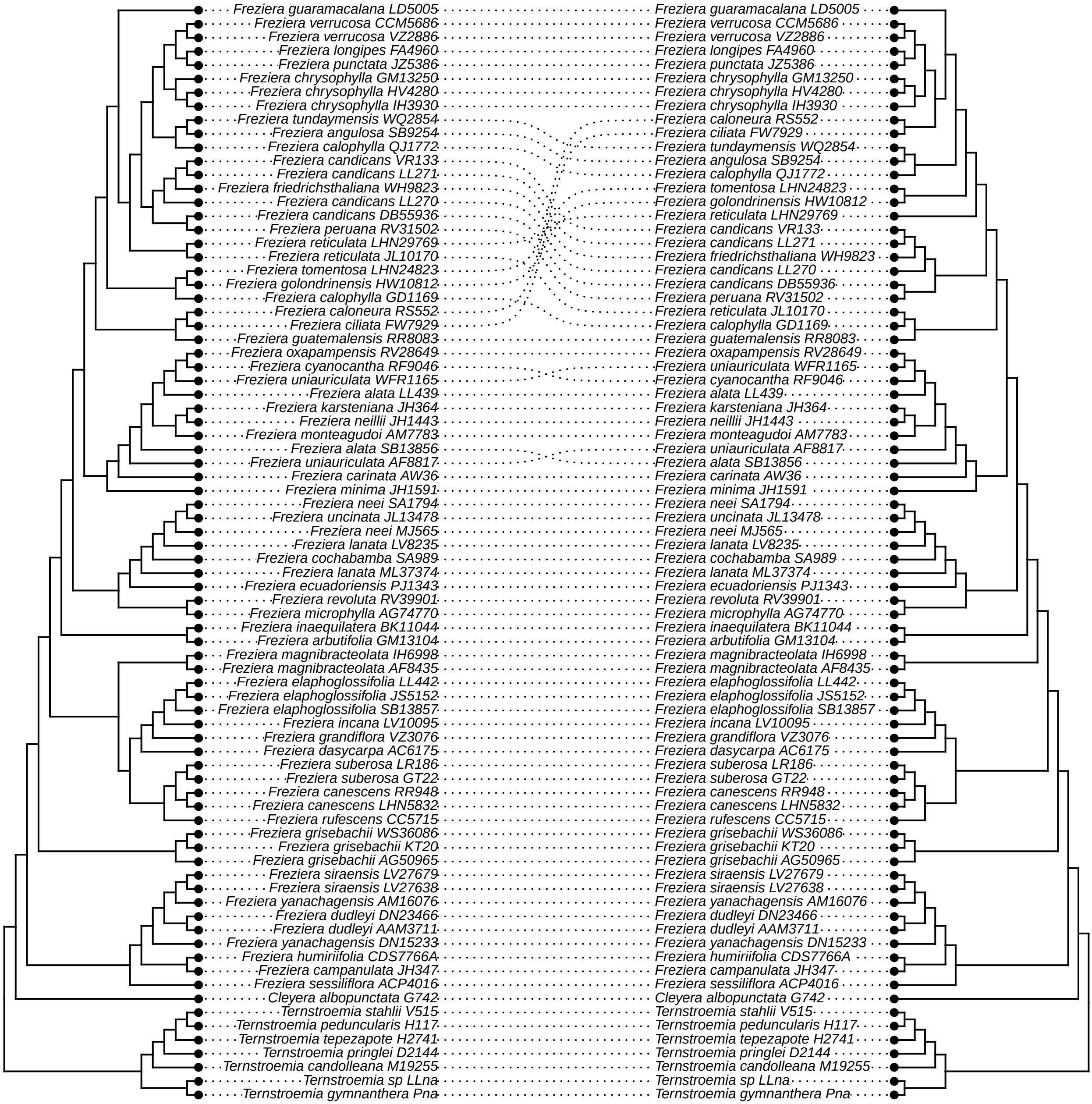

### nn_bipartition_v_hp2_long.pdf

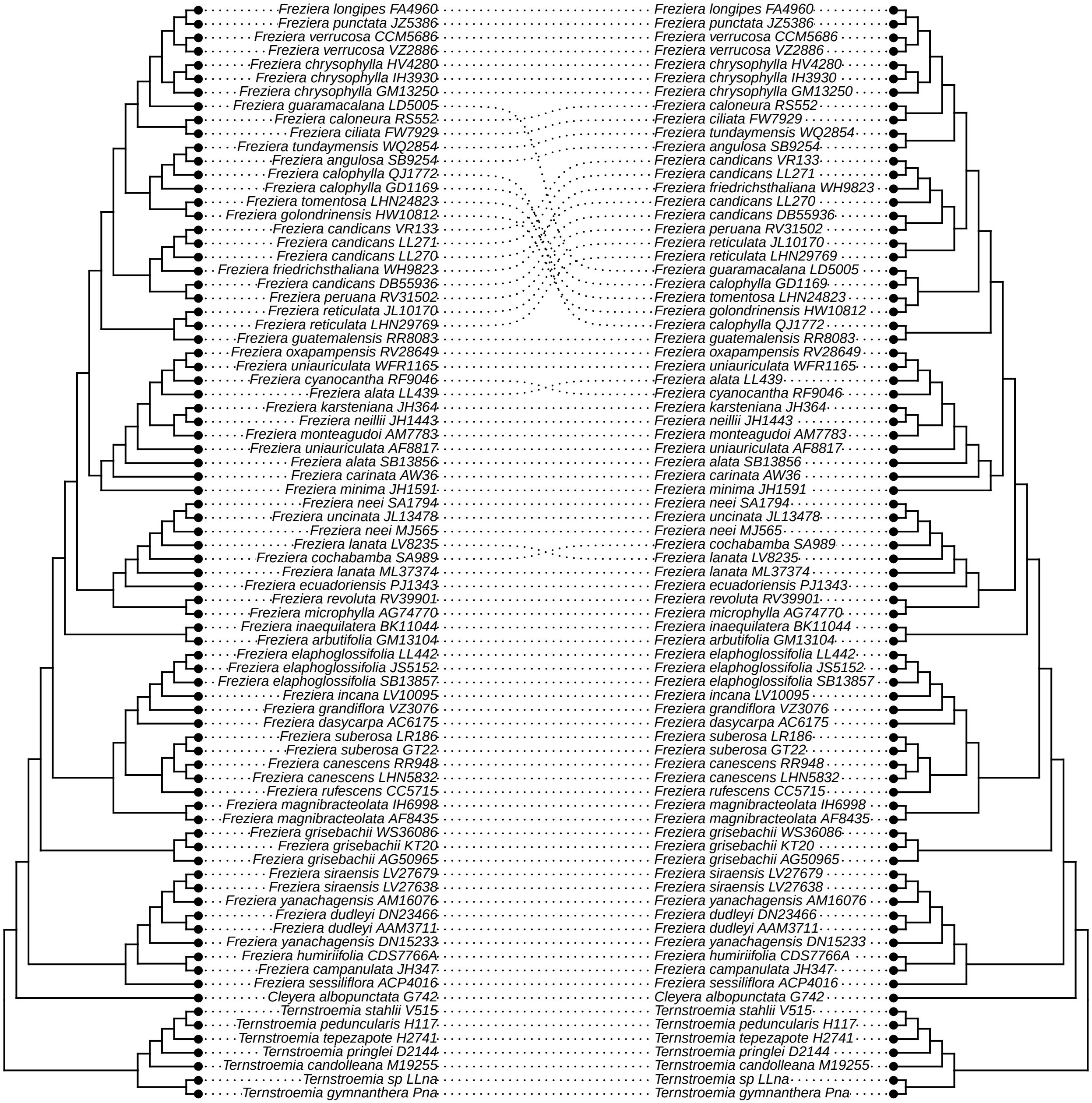

### oo_bipartition_v_MO.pdf

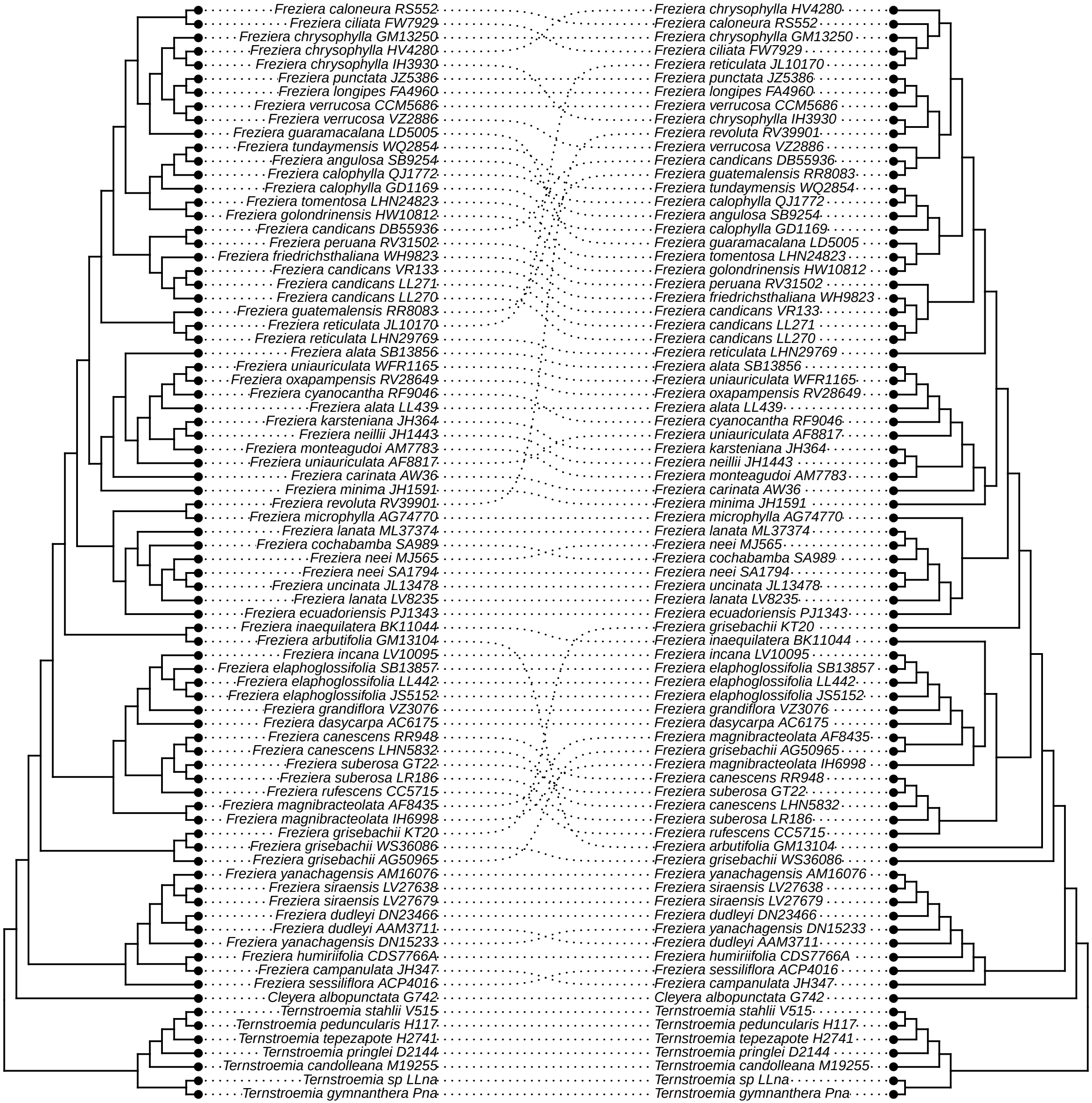

### pp_bipartition_v_prop_PI.pdf

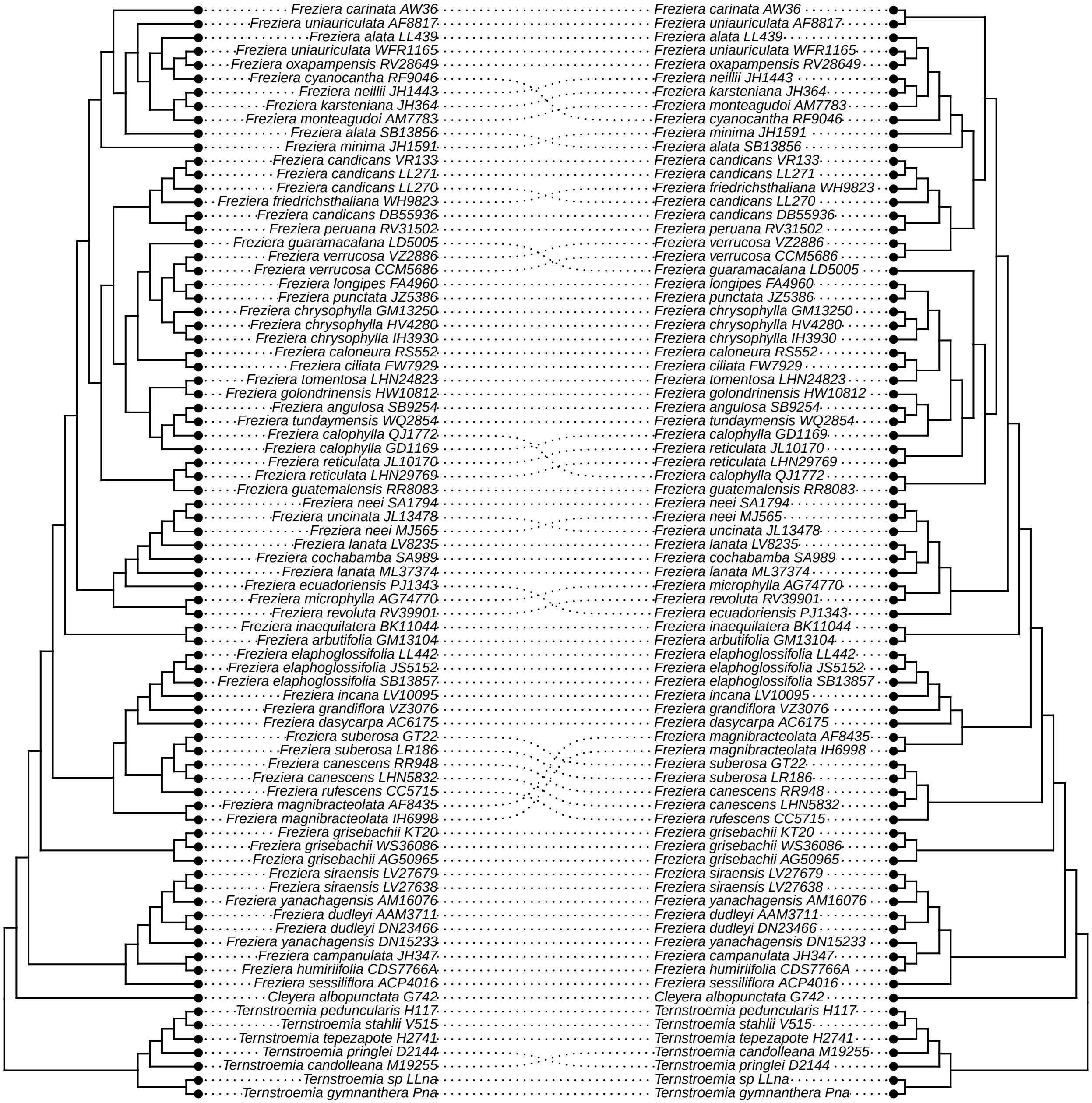

### qq_bipartition_v_internal.pdf

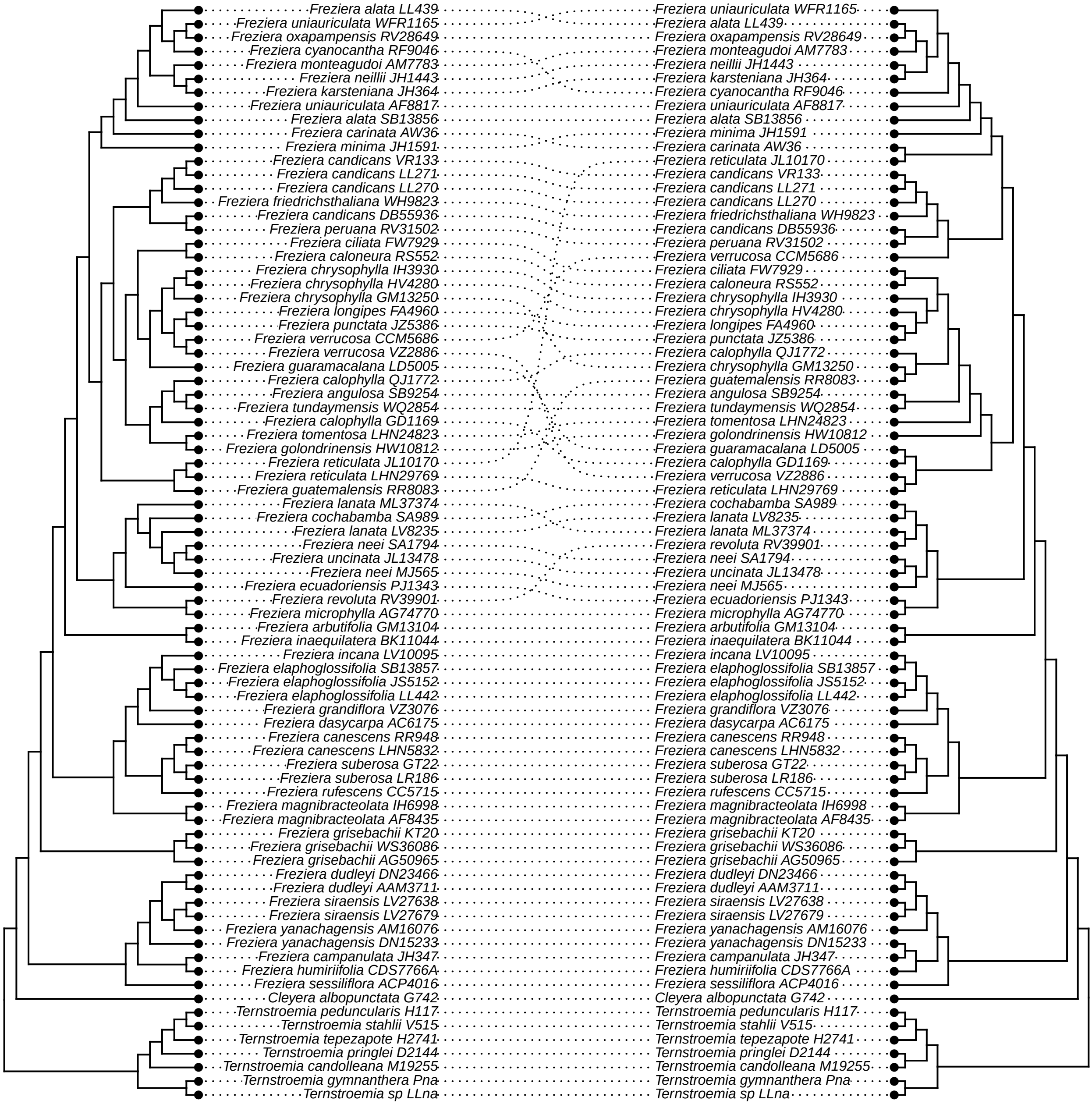

### rr_bipartition_v_avg_BS.pdf

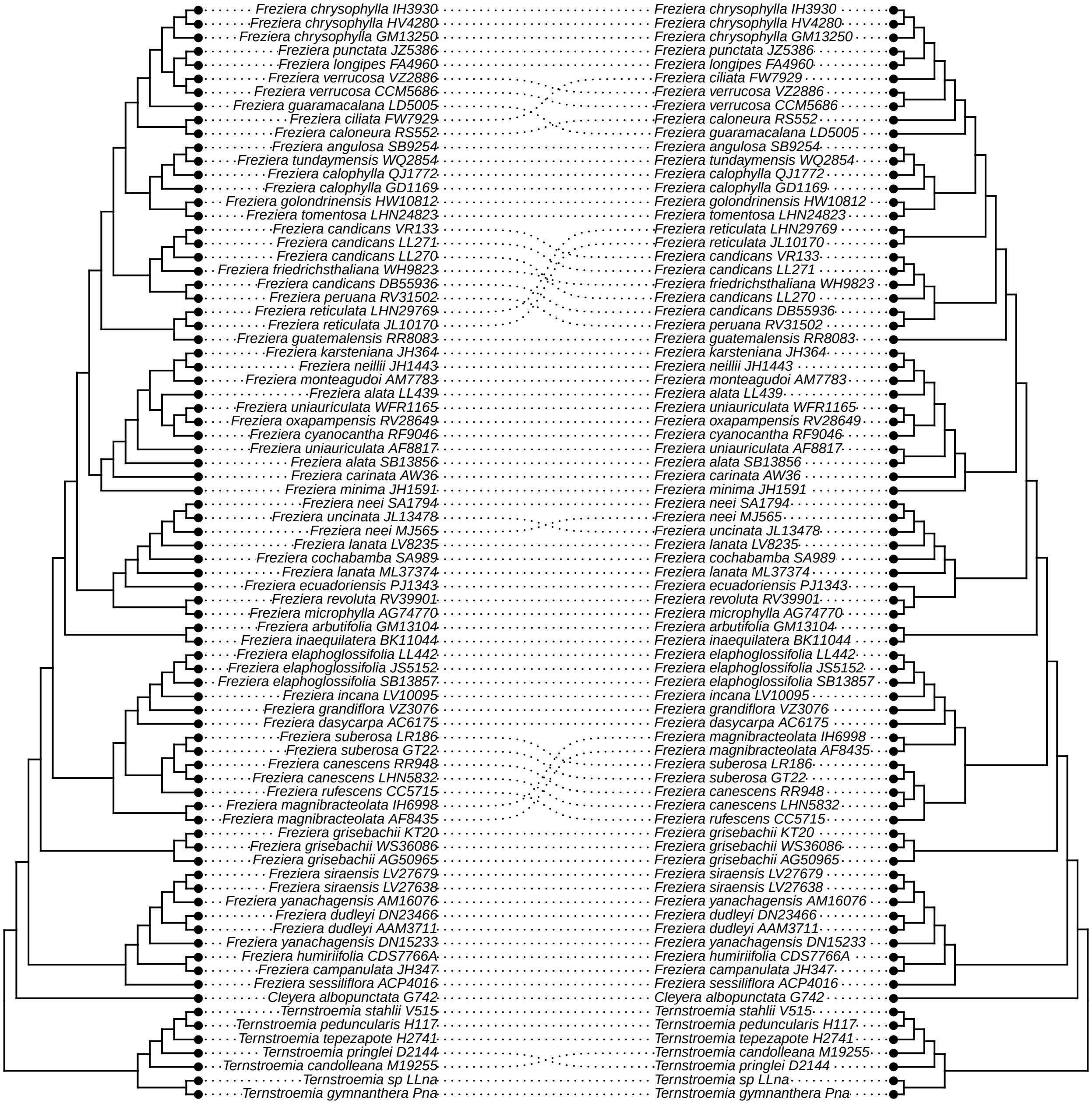

### ss_bipartition_v_tree_length.pdf

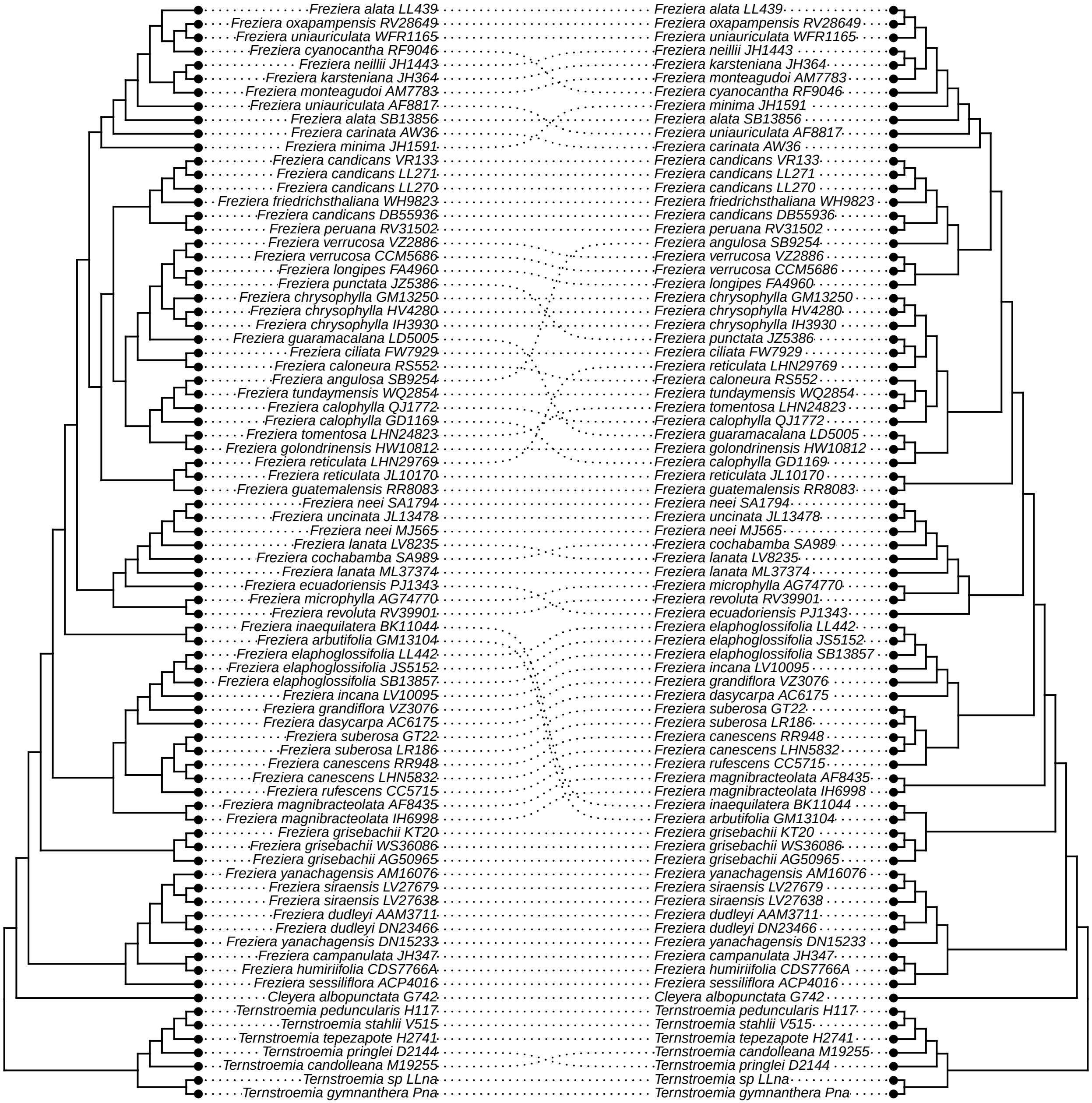

### tt_hp2_long_vs_unfiltered.pdf

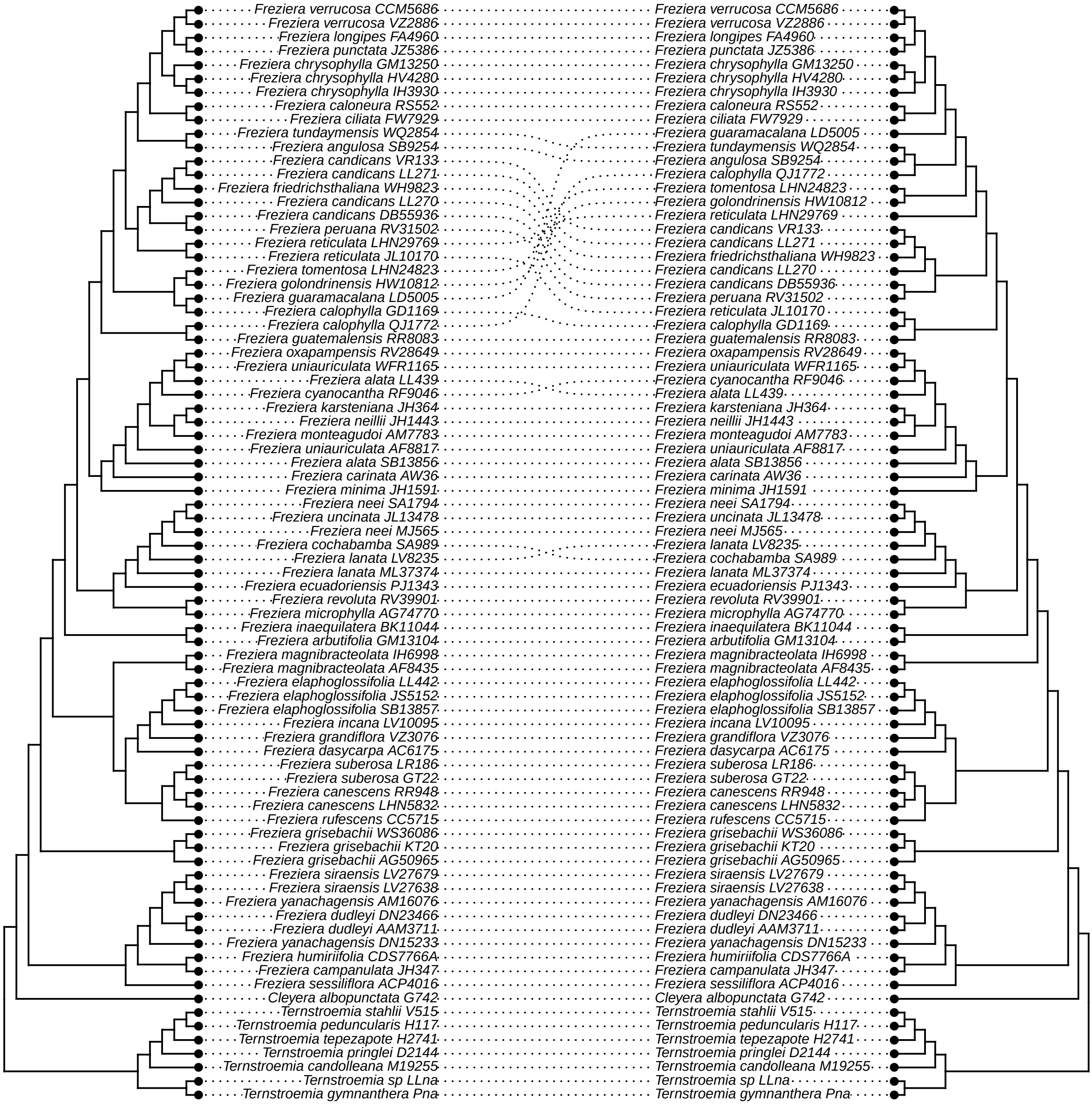

### uu_MO_v_unfiltered.pdf

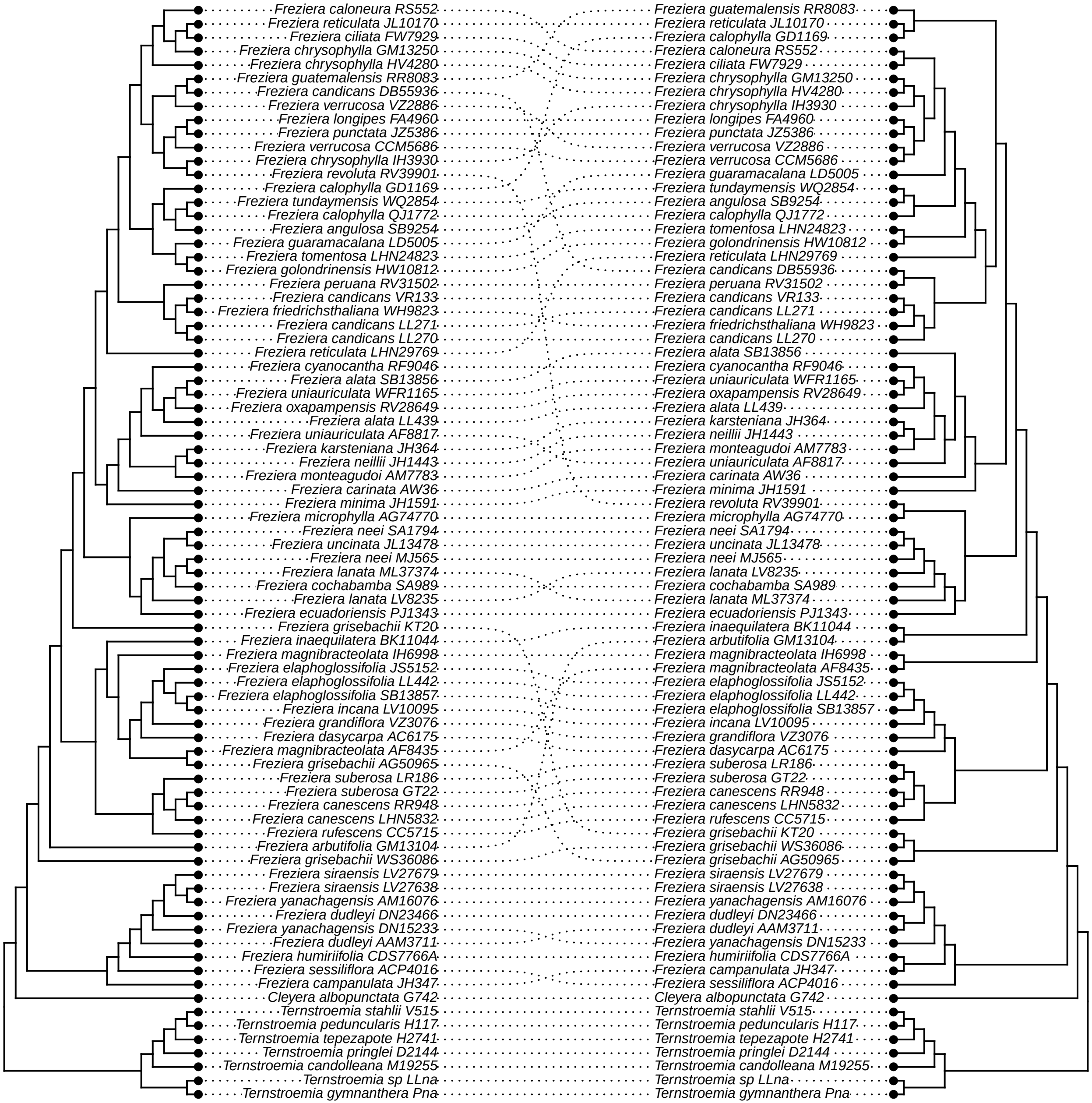

### vv_prop_PI_v_unfiltered.pdf

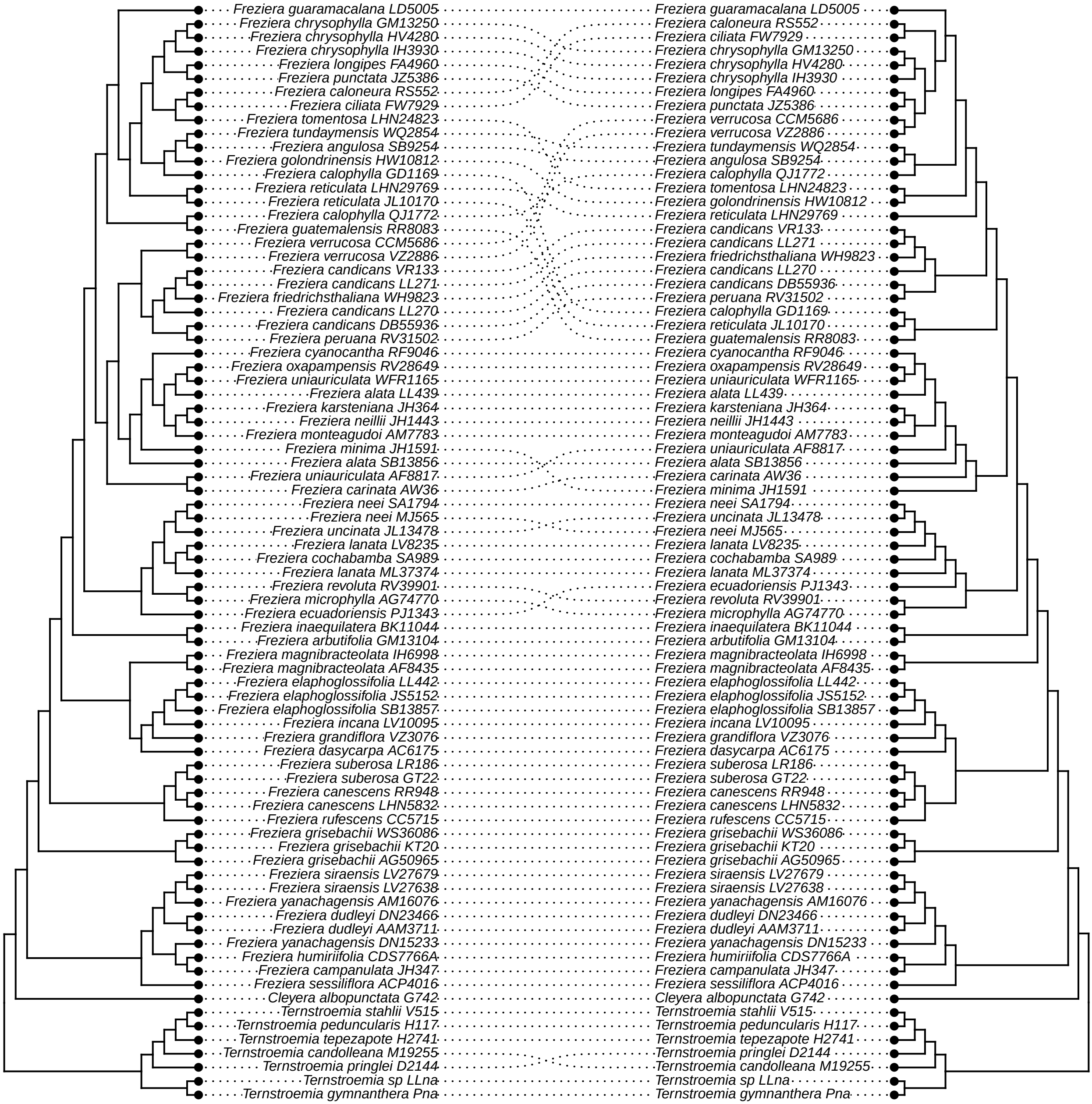

### xx_internal_v_unfiltered.pdf

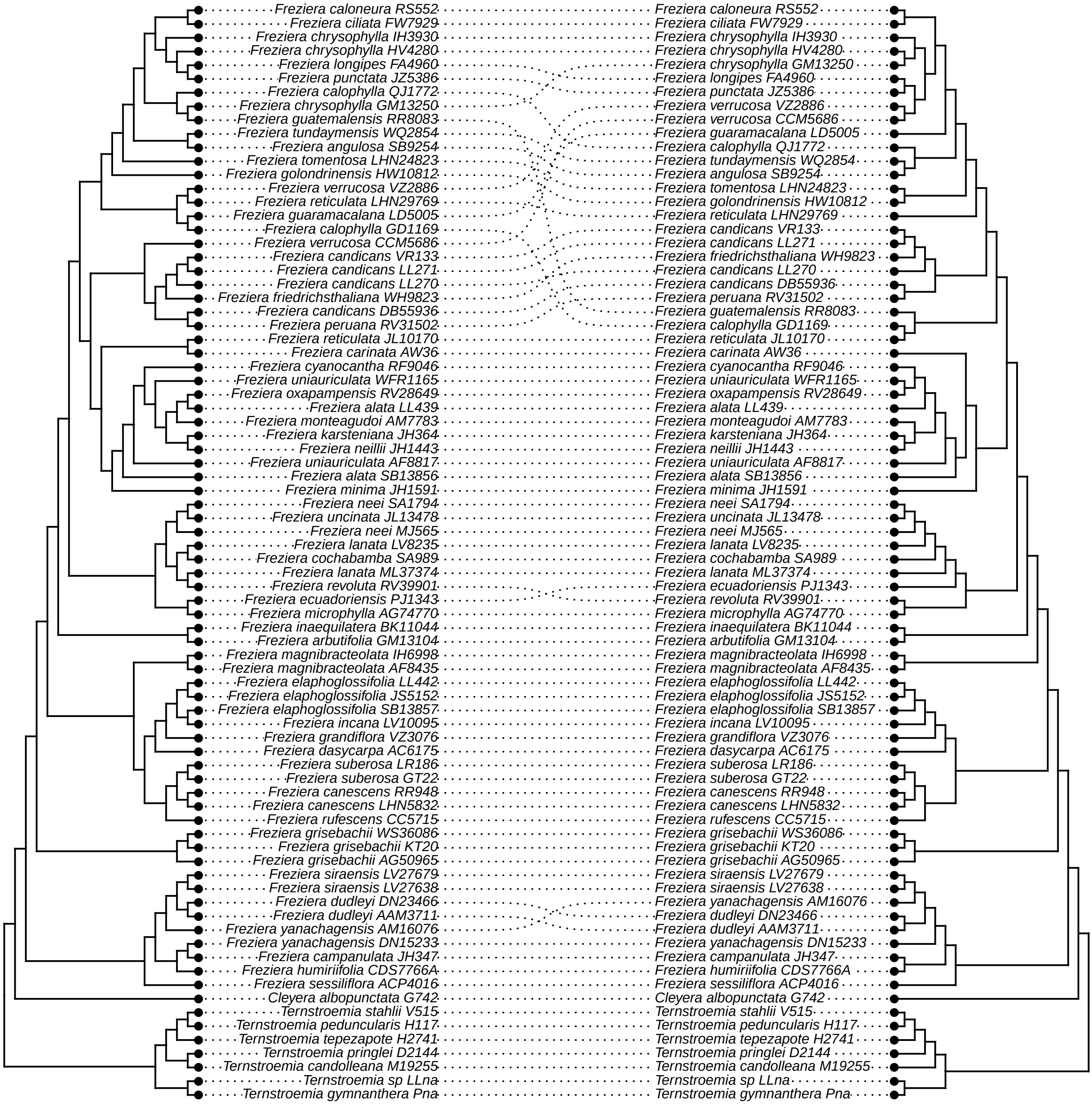

### yy_avg_BS_v_unfiltered.pdf

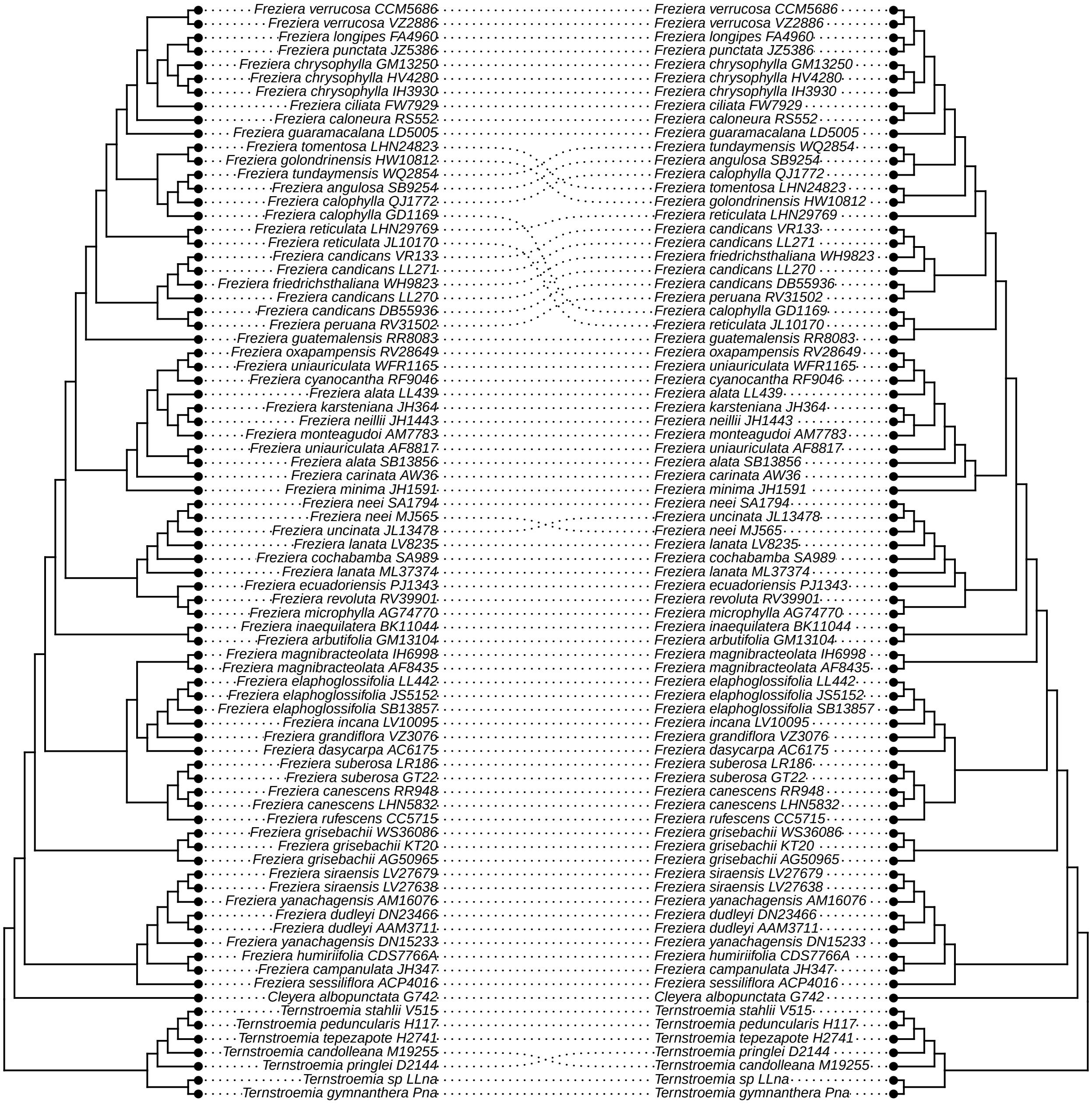

### zz_tree_length_v_unfiltered.pdf

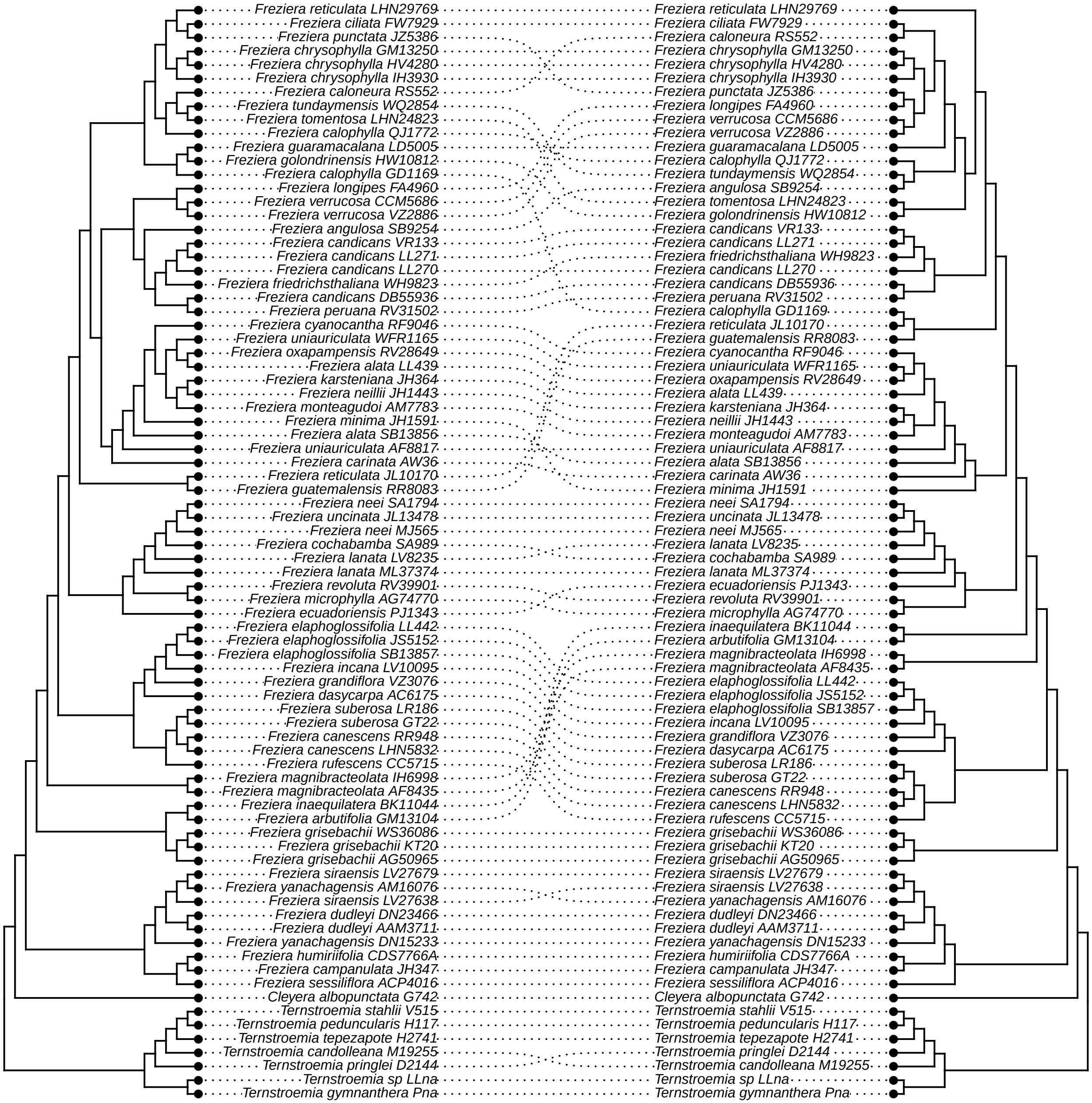
