## Supplementary material for "Strong phylogenetic signal despite high phylogenomic complexity in an Andean plant radiation (*Freziera,* Pentaphylacaceae)": Figure S2

a) ASTRAL-III species tree for *orthologs.unfiltered* dataset

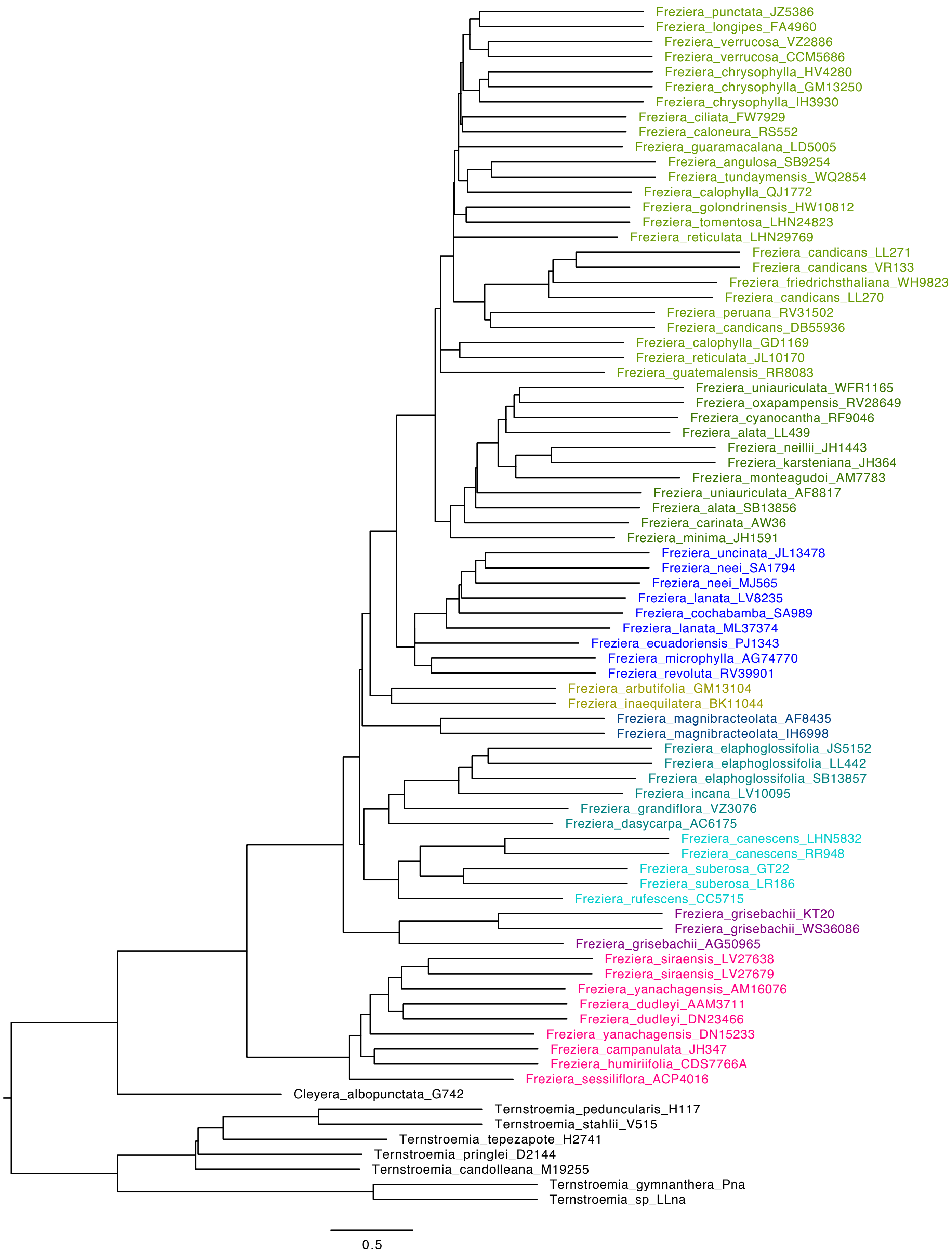

b) ASTRAL-III species tree for *orthologs.by.eye* dataset

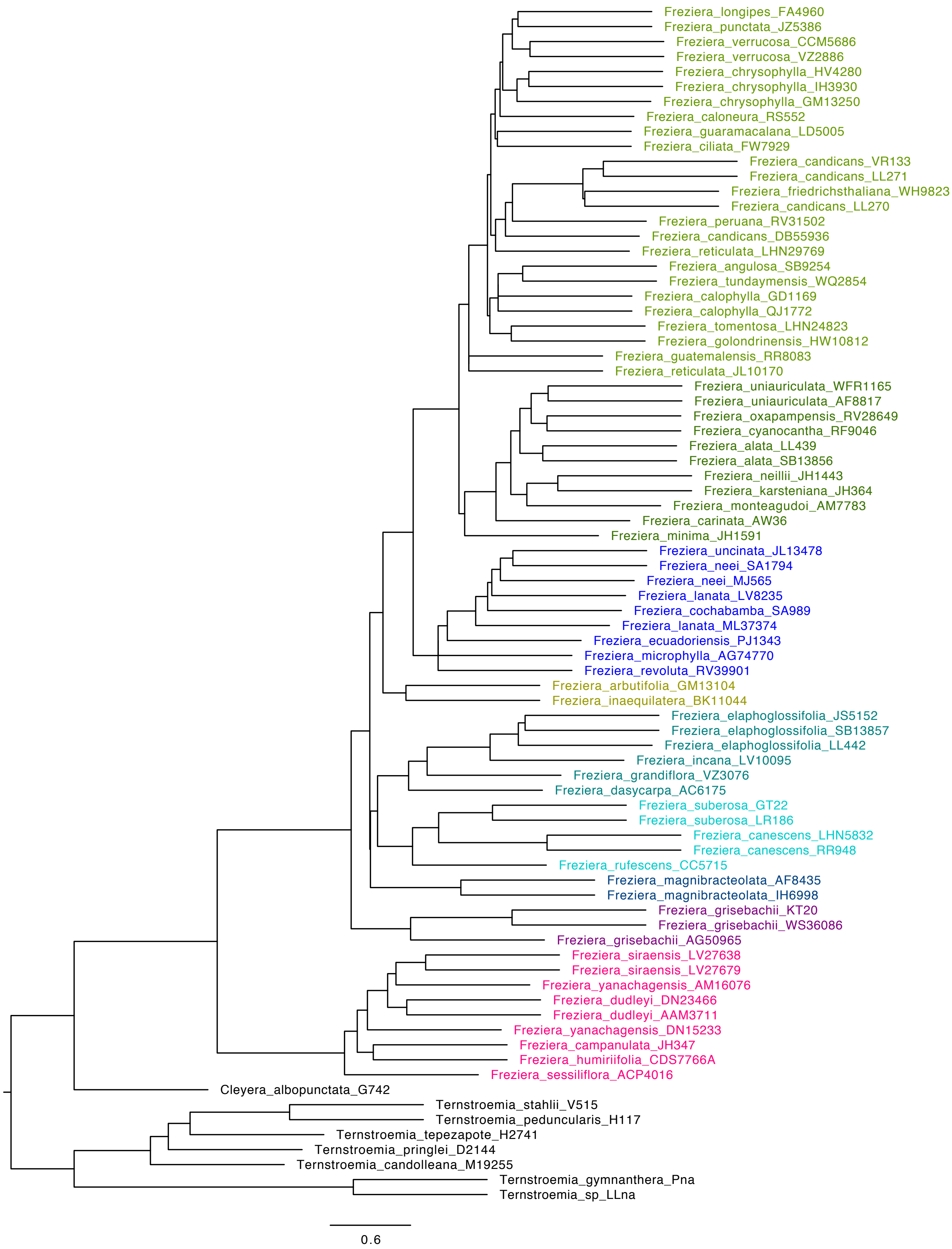

c) ASTRAL-III species tree for *orthologs.HybPiper2.long* dataset

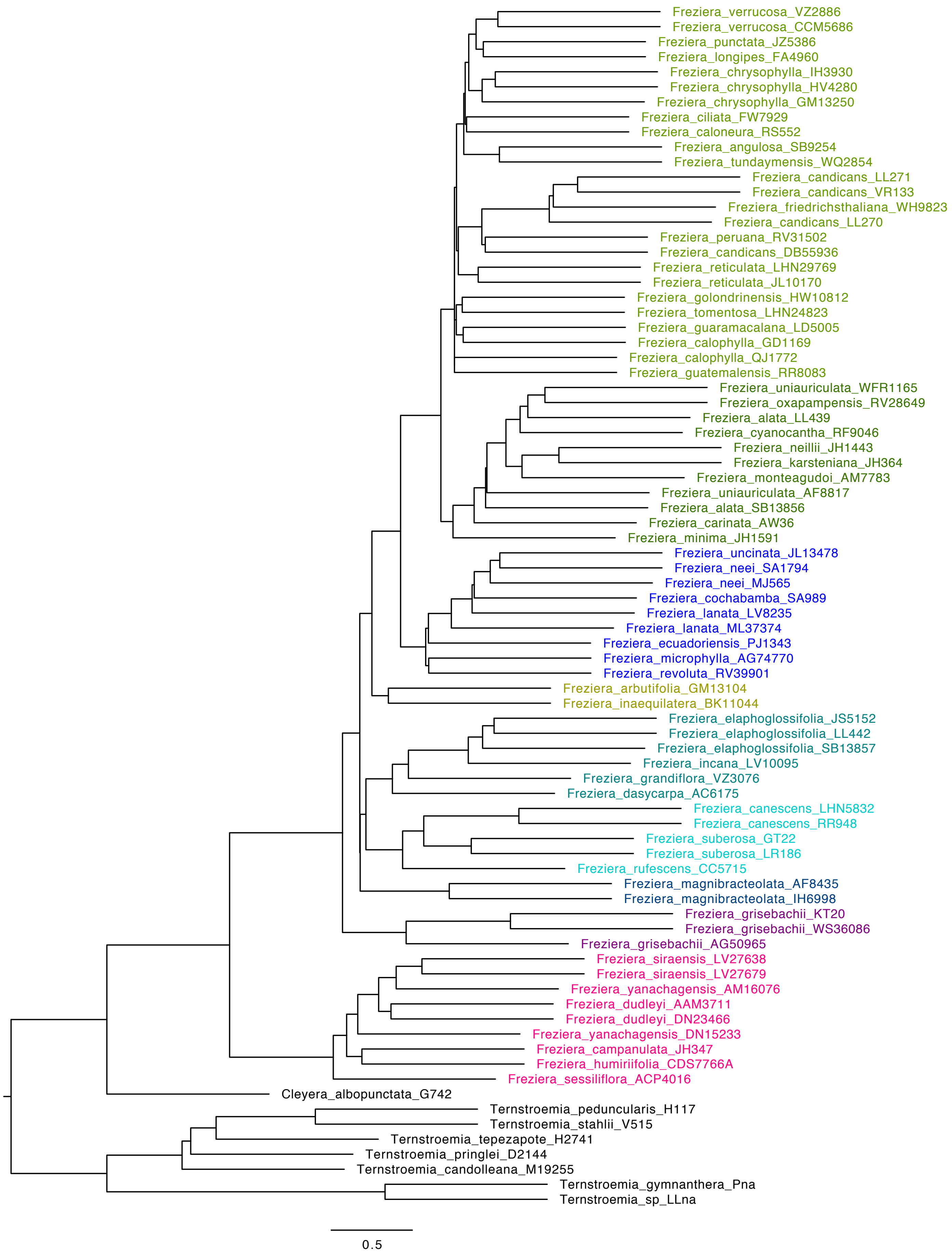

d) ASTRAL-III species tree for *orthologs.HybPiper2.no.warnings* dataset

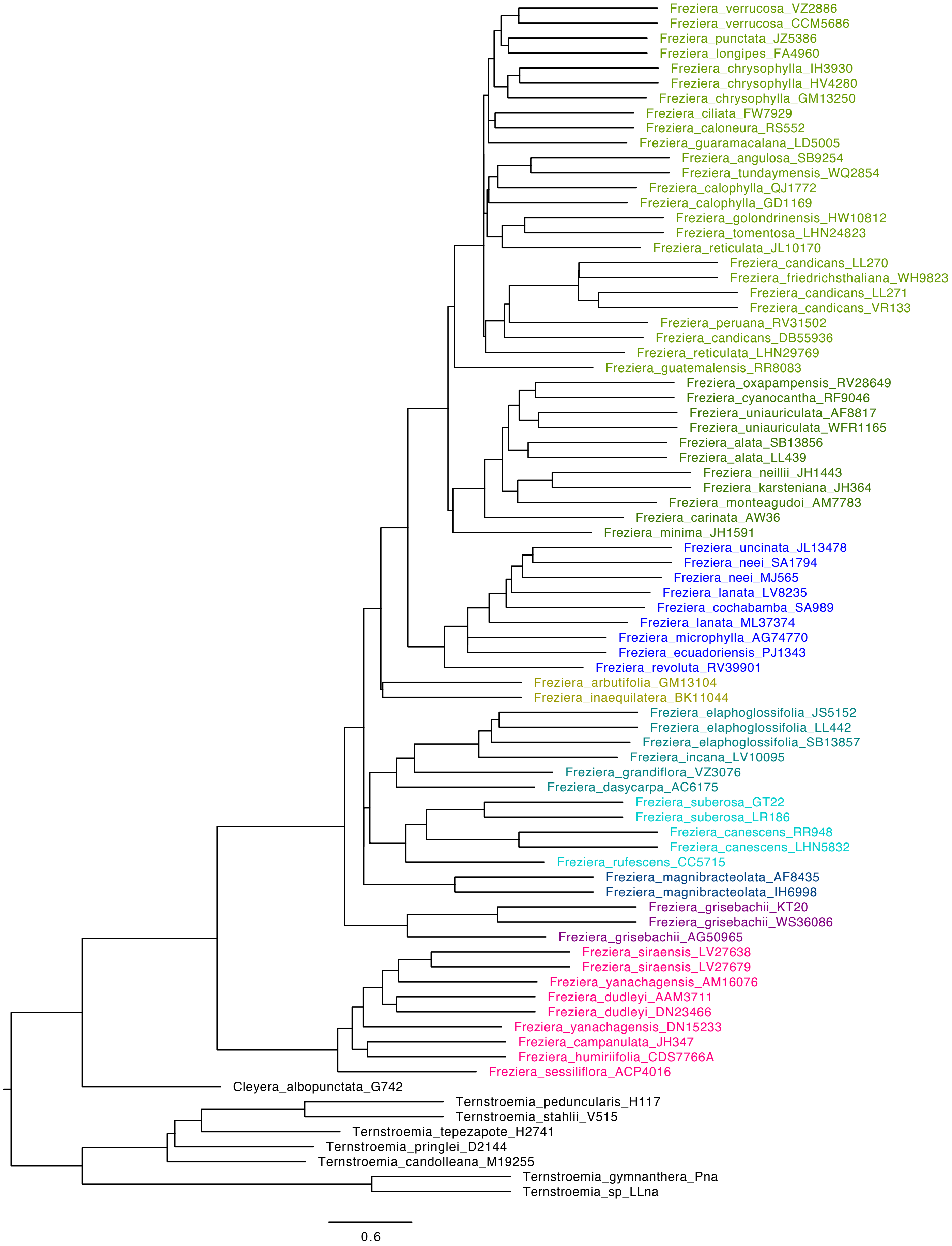

e) ASTRAL-III species tree for *HybPhaser* dataset

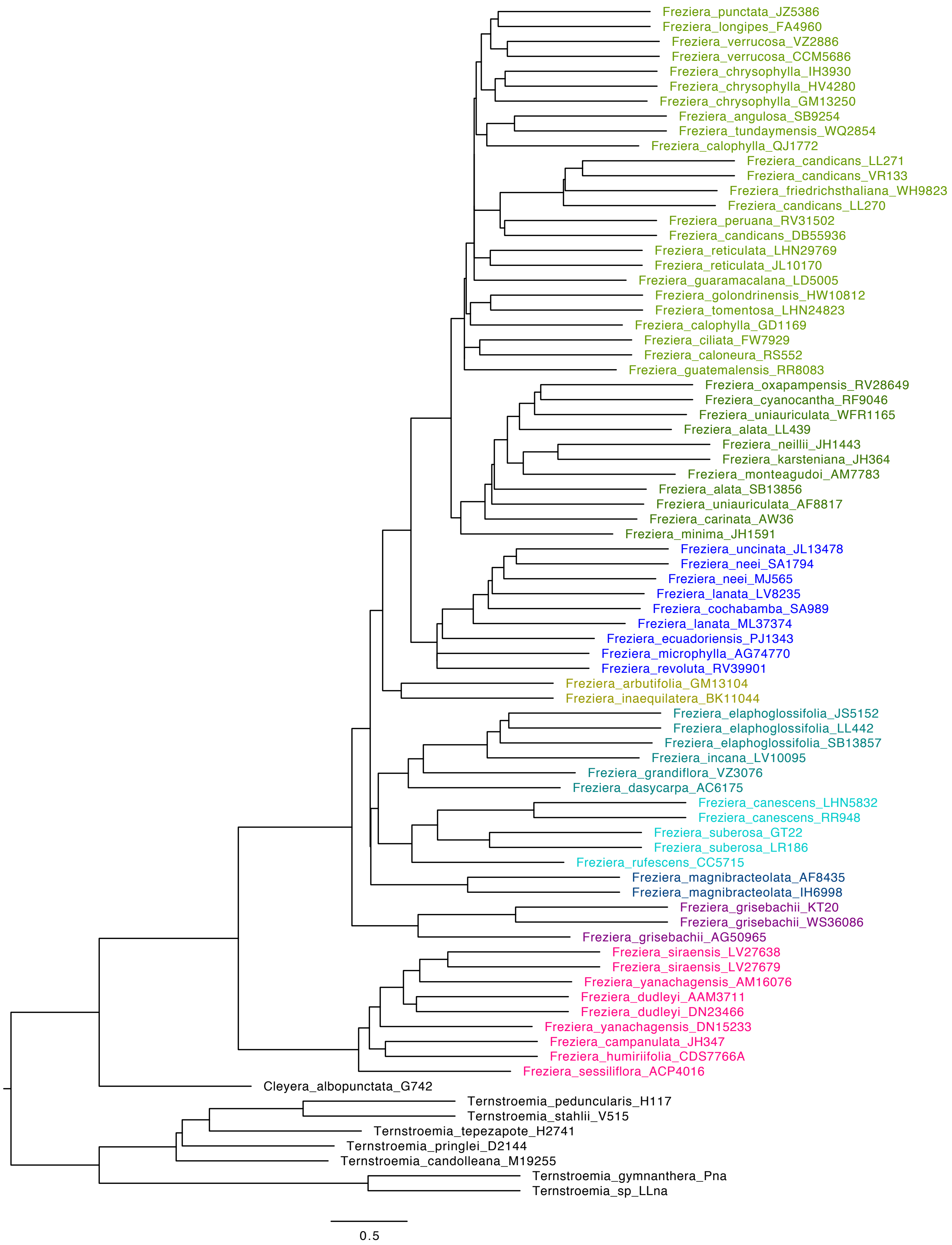

f) ASTRAL-III species tree for *MO* dataset

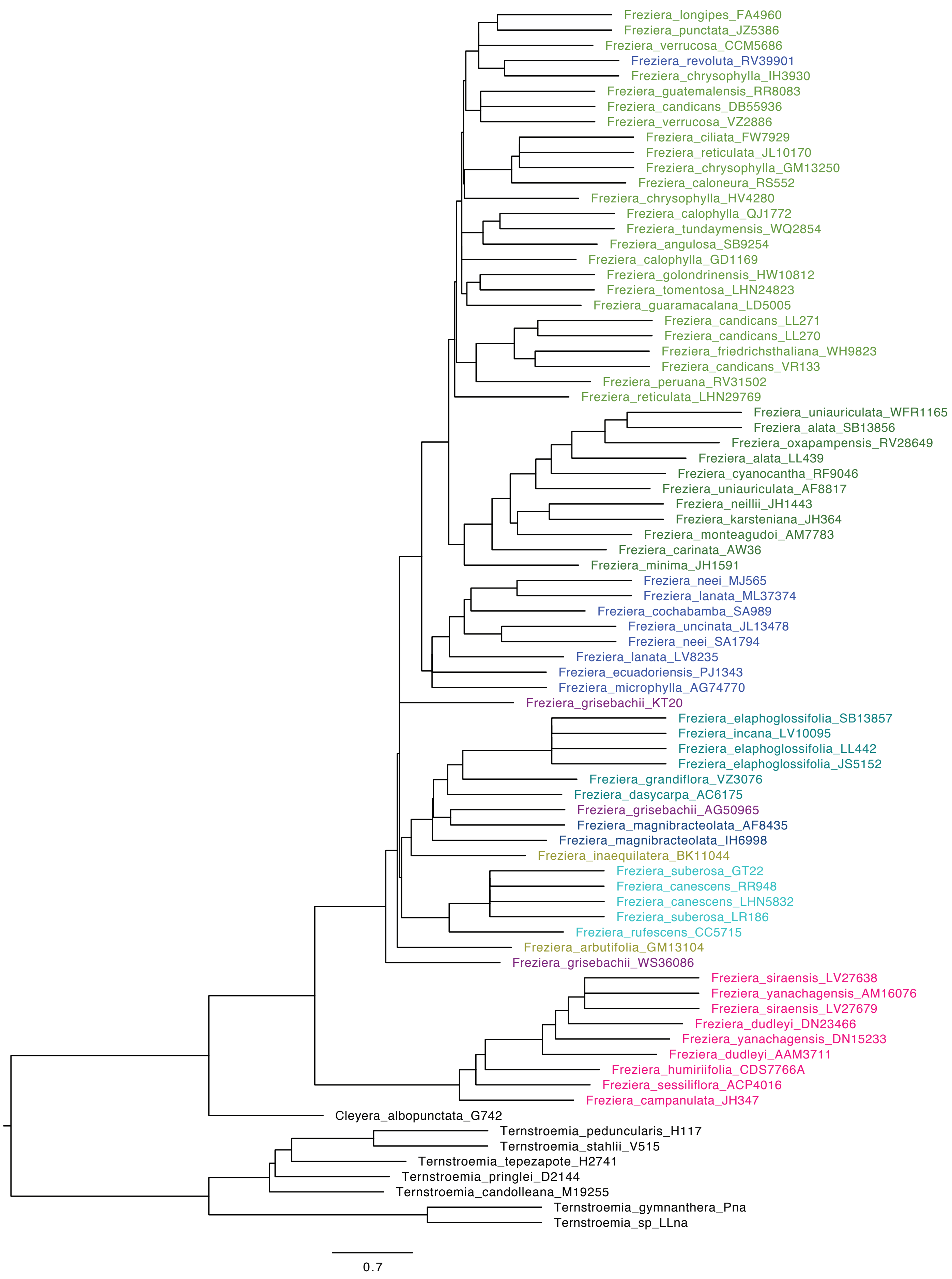

g) ASTRAL-III species tree for *proportion.PI* dataset

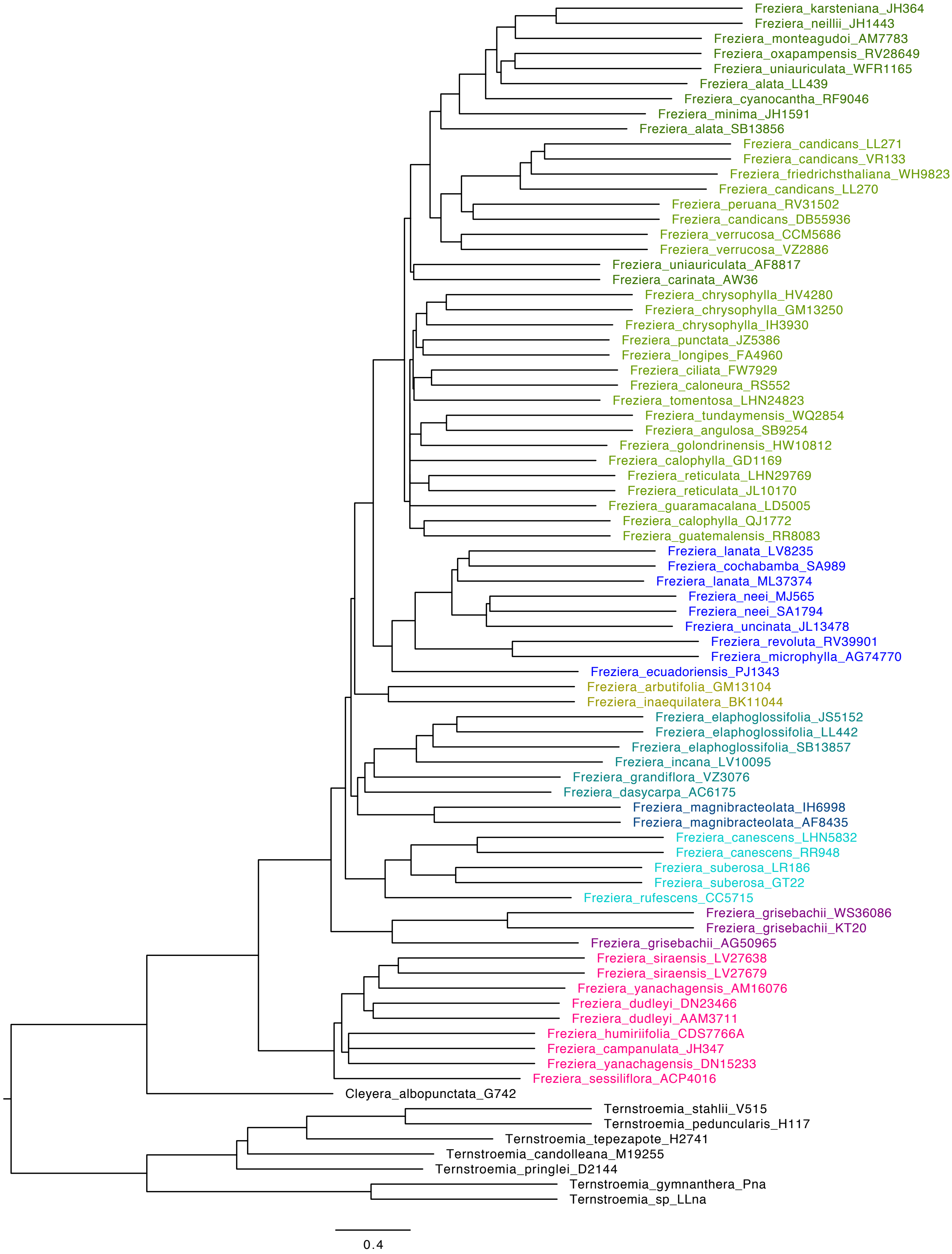

h) ASTRAL-III species tree for *proportion.internal.branch.lengths* dataset

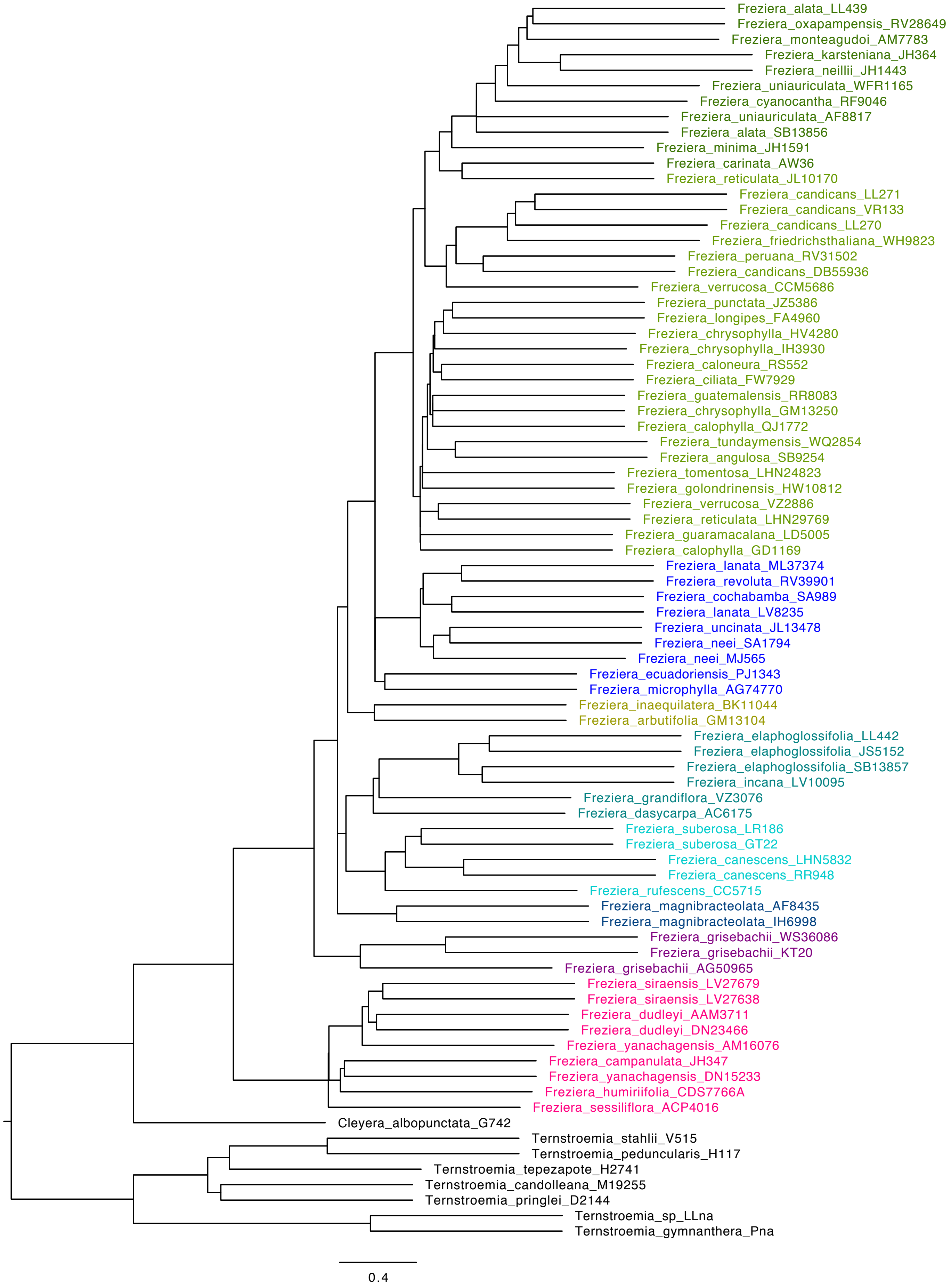

i) ASTRAL-III species tree for *average.BS* dataset

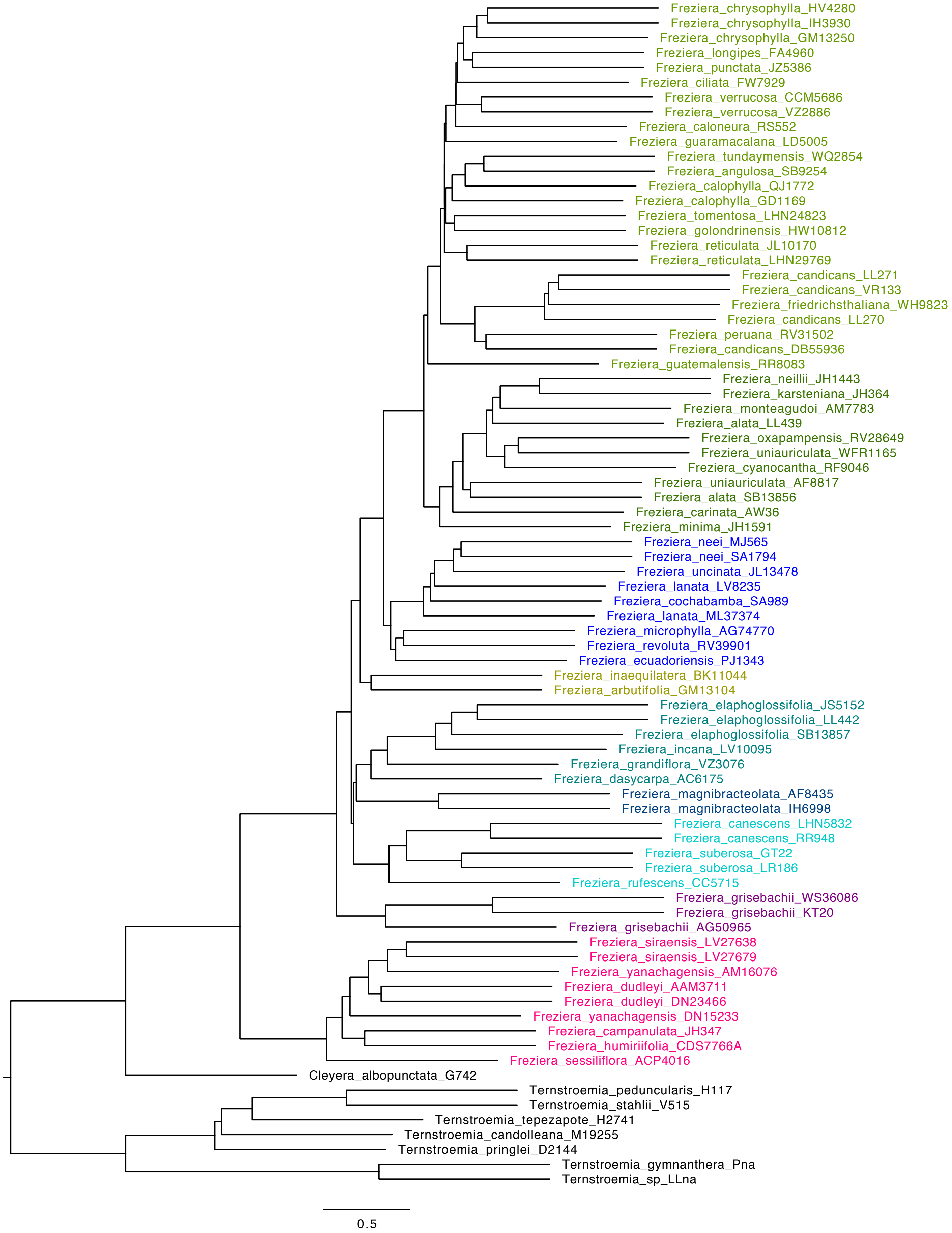

j) ASTRAL-III species tree for *tree.length* dataset

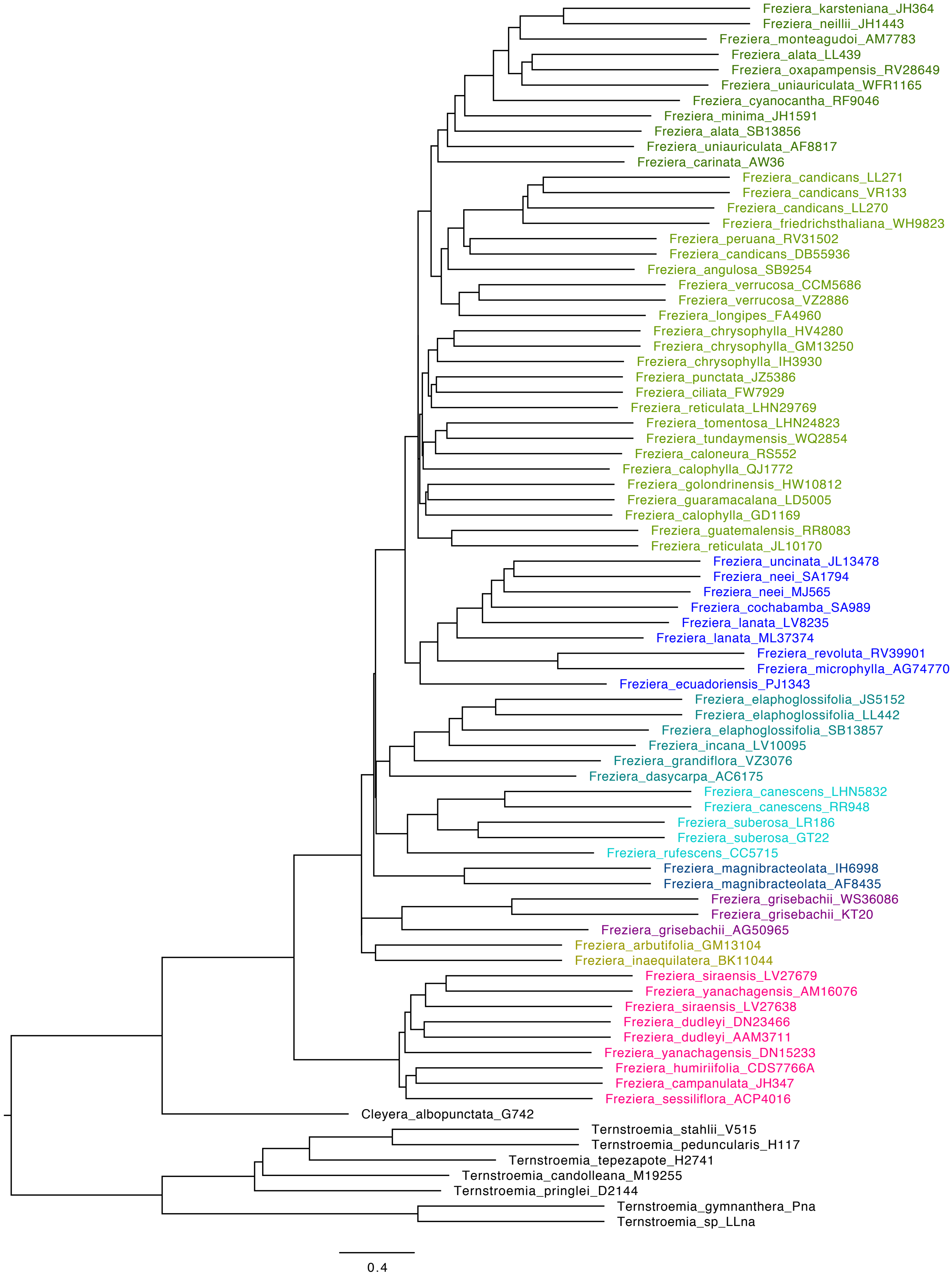

k) ASTRAL-III species tree for *bipartition* dataset

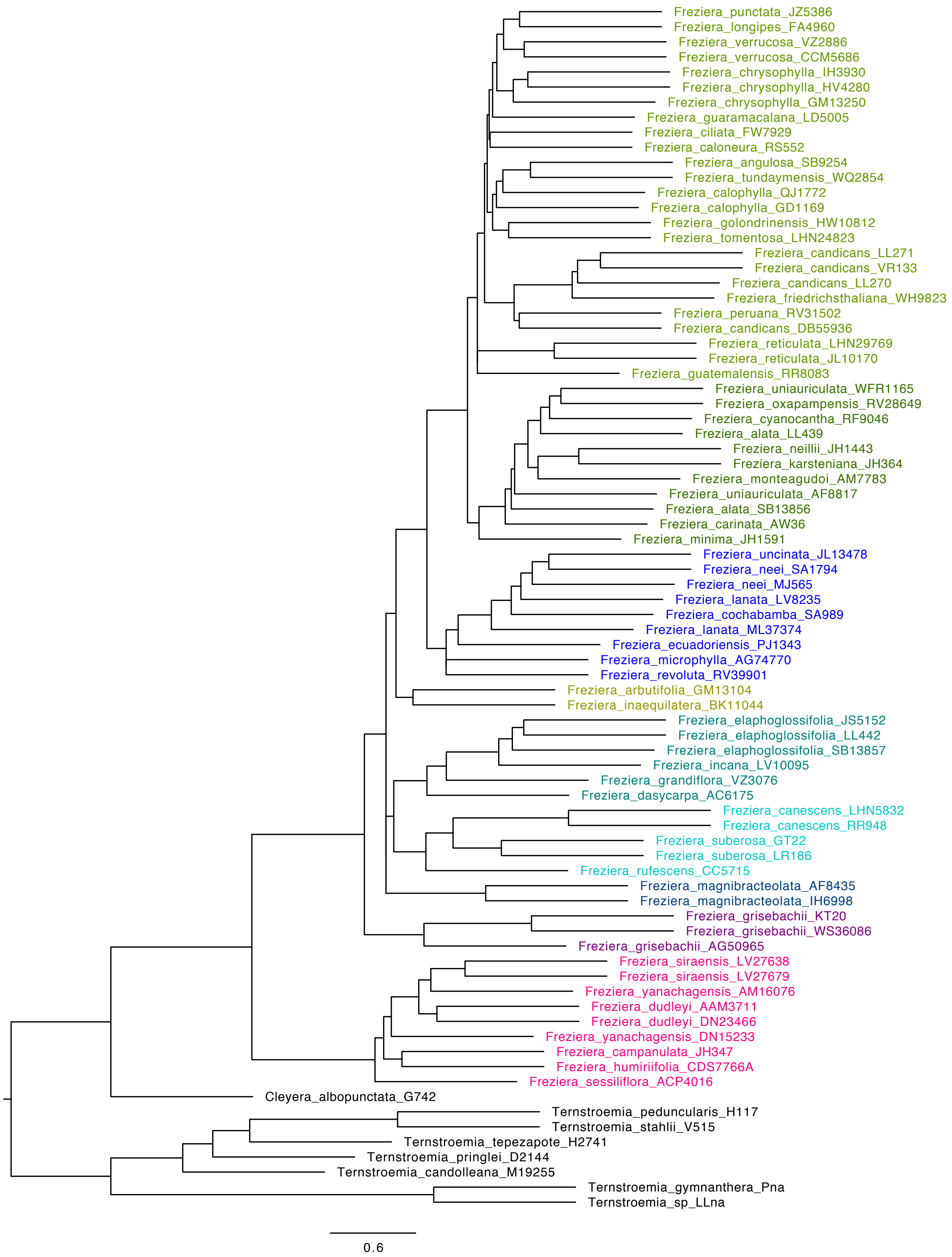
